# Glycan analysis probes inspired by human lectins for investigating host-microbe crosstalk

**DOI:** 10.1101/2024.12.24.630132

**Authors:** Soumi Ghosh, Rajeev Chorghade, Roger C. Diehl, Greg J. Dodge, Sunhee Bae, Amanda E. Dugan, Melanie Halim, Michael G Wuo, Helen Bartlett, Liam Herndon, Laura L. Kiessling, Barbara Imperiali

## Abstract

Human lectins are critical carbohydrate-binding proteins that recognize diverse glycoconjugates from microorganisms and can play a key role in host-microbe interactions. Despite their importance in immune recognition and pathogen binding, the specific glycan ligands and functions of many human lectins remain poorly understood. Using previous proof-of-concept studies on selected lectins as the foundation for this work, we present ten additional glycan analysis probes (GAPs) from a diverse set of human soluble lectins, offering robust tools to investigate glycan-mediated interactions. We describe a protein engineering platform that enables scalable production of GAPs that maintain native-like conformations and oligomerization states, equipped with functional reporter tags for targeted glycan profiling. We demonstrate that the soluble GAP reagents can be used in various applications, including glycan array analysis, mucin-binding assays, tissue staining, and microbe binding in complex populations. These capabilities make GAPs valuable for dissecting interactions relevant to understanding host responses to microbes. The tools can be used to distinguish microbial from mammalian glycans, which is crucial for understanding the cross-target interactions of lectins in a physiological environment where both glycan types exist. GAPs have potential as diagnostic and prognostic tools for detecting glycan alterations in chronic diseases, microbial dysbiosis, and immune-related conditions.

## Introduction

Human soluble lectins (Peiffer, A.L., Dugan, A.E., et al. 2024, Raposo, C.D., Canelas, A.B., et al. 2021, Sahly, H., Keisari, Y., et al. 2008, Schnider, B., M’Rad, Y., et al. 2023) are carbohydrate-binding proteins in the extracellular matrix that play critical roles in host-microbe interactions. These lectins have evolved to recognize distinct glycan structures and drive various physiological functions, including pathogen recognition and innate immune functions. Despite the biological significance of these human lectins, the glycan specificity at the host-microbe interface of many remains unclear, although studies on their specificity towards N- and O-linked mammalian glycans have been reported (Kletter, D., Singh, S., et al. 2013). Currently, there is an incomplete understanding of the binding profiles of soluble human lectins towards microbial glycan targets, which is particularly challenging as microbial cell surface glycans comprise remarkably diverse monosaccharide building blocks and glycosidic linkages absent in the mammalian glycome (Adibekian, A., Stallforth, P., et al. 2011, Griffin, M.E. and Hsieh-Wilson, L.C. 2022, Herget, S., Toukach, P.V., et al. 2008, Imperiali, B. 2019). As human lectins serve as “readers” of complex microbial surface glycans, they represent opportunities for developing reagents for understanding host cell recognition, as well as bacterial colonization and pathogenesis.

Recently, we established the proof-of-concept for producing and utilizing glycan analysis probes (GAPs) from soluble human lectins to decode host specificity to microbes within complex microbiomes. GAPs represent soluble protein reagents that can be engineered with fluorophores or affinity handles, and applied to specifically bind to microbial cell surface glycans, enabling their detection in mixed microbial populations and enrichment for analysis and characterization. GAPs provide capabilities that are complementary to lectin arrays (Benjamin, S.V., Jégouzo, S.A.F., et al. 2024, Hsu, K.-L. and Mahal, L.K. 2006). For example, we previously demonstrated that a GAP based on the human lectin Zymogen granule protein ZG16B selectively binds to a subset of commensal bacteria in the oral cavity, regulating their growth by recruiting oral mucin MUC7 to maintain homeostasis (Ghosh, S., Ahearn, C.P., et al. 2023). The ZG16B-GAP also allowed the identification of the glycan epitope on cell wall polysaccharide on the targeted bacteria involved in the lectin-glycan interaction. In other examples, mannose-binding lectin (MBL) and intelectin-1 (hItln1) GAPs were shown to bind distinct microbial populations in stool samples from healthy individuals and those with inflammatory bowel disease (McPherson, R.L., Isabella, C.R., et al. 2023), suggesting potential applications of GAPs as diagnostic markers for diseases associated with microbial dysbiosis.

Building on these studies we have expanded the suite of GAPs to include an additional ten human soluble lectins, presenting new resources for investigating and understanding human lectin function and providing insights into microbial glycan diversity and glycan-mediated host—microbe interactions. Here, we detail a modular platform (Figure 1A) for tailoring soluble human lectins as glycan analysis probes (GAPs) for visualizing the glycan-binding properties to gain insight into their biological functions. We selected a series of human lectins based on tissue distribution (Figure 1B), serum abundance (Figure 1B), protein structure (Figure 1C), and functional diversity as candidates for conversion into functionalized GAPs. The generalized approach provides lectins bearing a range of functional tags in high yield and purity for use in biophysical and biochemical assays. An essential feature of the GAP production pipeline is its compatibility with newly targeted lectins. This enables rapid screening of expression and purification conditions and co-expression tag optimization.

**Figure 1:**
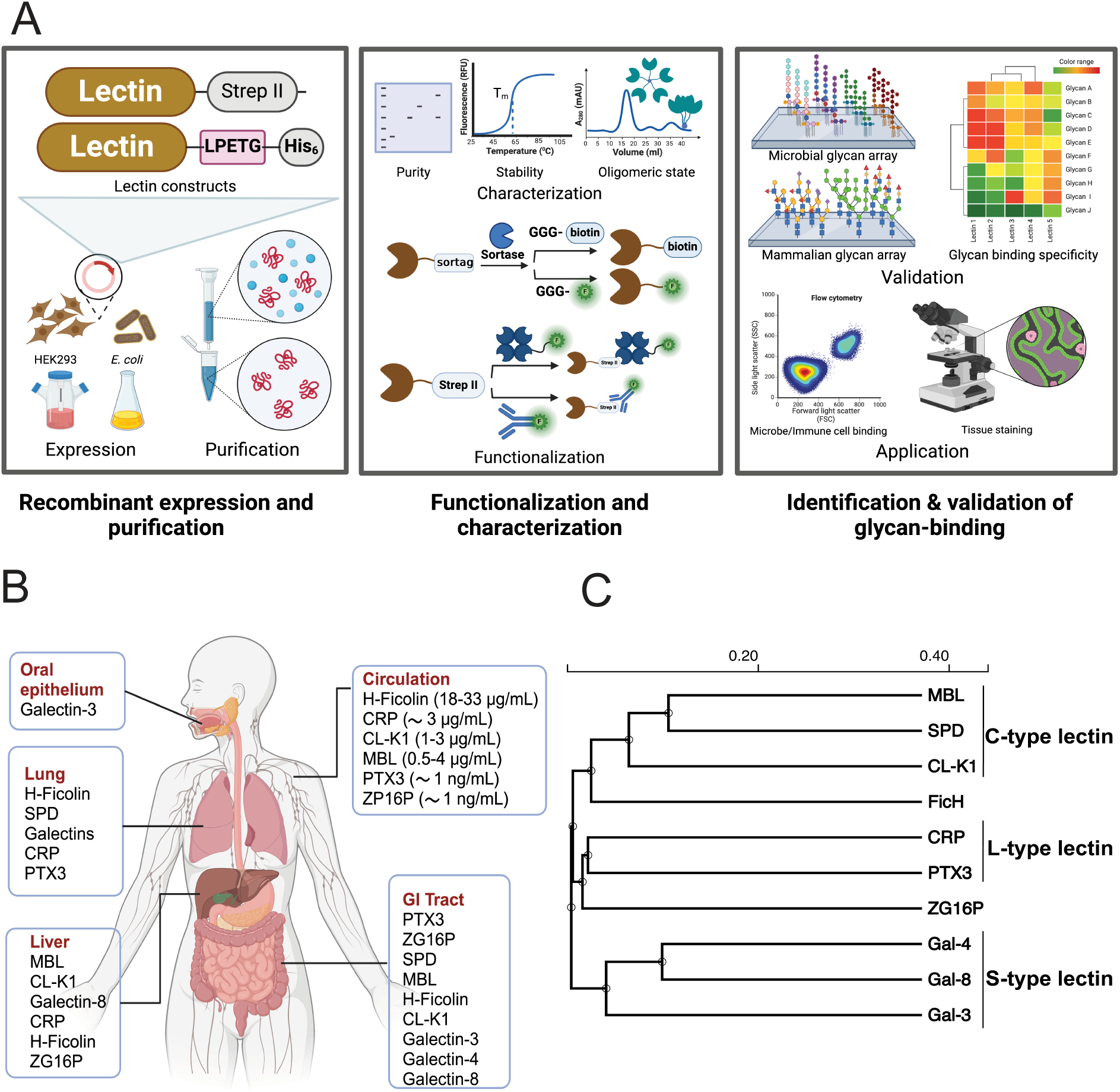
Production of Glycan Analysis Probes (GAPs) from Human Soluble Lectins (A) Schematic diagram of the workflow of developing through recombinant production of human soluble lectins, their functionalization with reporter tags, and identification of their glycan binding partners. (B) Tissue-specific distribution and abundance of human soluble lectins used in this study (adopted from (Peiffer, A.L., Dugan, A.E., et al. 2024). (C) Guide tree representing the structural homology of the human soluble lectins used in the study, generated by multiple sequence alignment in Clustal Omega. The scale indicates the length of the branches, indicating the evolutionary distance between the sequences.

Our comparative analysis reveals distinct binding profiles of lectin-based GAPs assessed by microbial and mammalian glycan arrays and shows applications of the GAPs in mucin binding and tissue staining analyses. As seen with the ZG16B-GAP, such applications are particularly useful for exploring ancillary lectin interactions with host glycoproteins, which play a crucial role in mucosal immunity (Holmgren, J. and Czerkinsky, C. 2005). We present GAPs as a valuable resource that can be employed to delineate the roles of lectins in physiologically relevant environments in host-microbe interactions.

## Results

### Selection of representative lectins as targets for GAP development

We describe the development of a curated set of GAPs from soluble human lectins based on reported microbe binding and participation in innate immunity (see Table I). The lectins feature different tissue-specific distributions (Figure 1B), abundance, oligomeric states, potential ligand-specificity, and reported biological functions. We chose lectins from multiple structural classes (Figure 1C): including the C-type (calcium ion-dependent) lectins, L-type (leguminous lectins), and S-type (galectins) lectins.

**Table 1:**
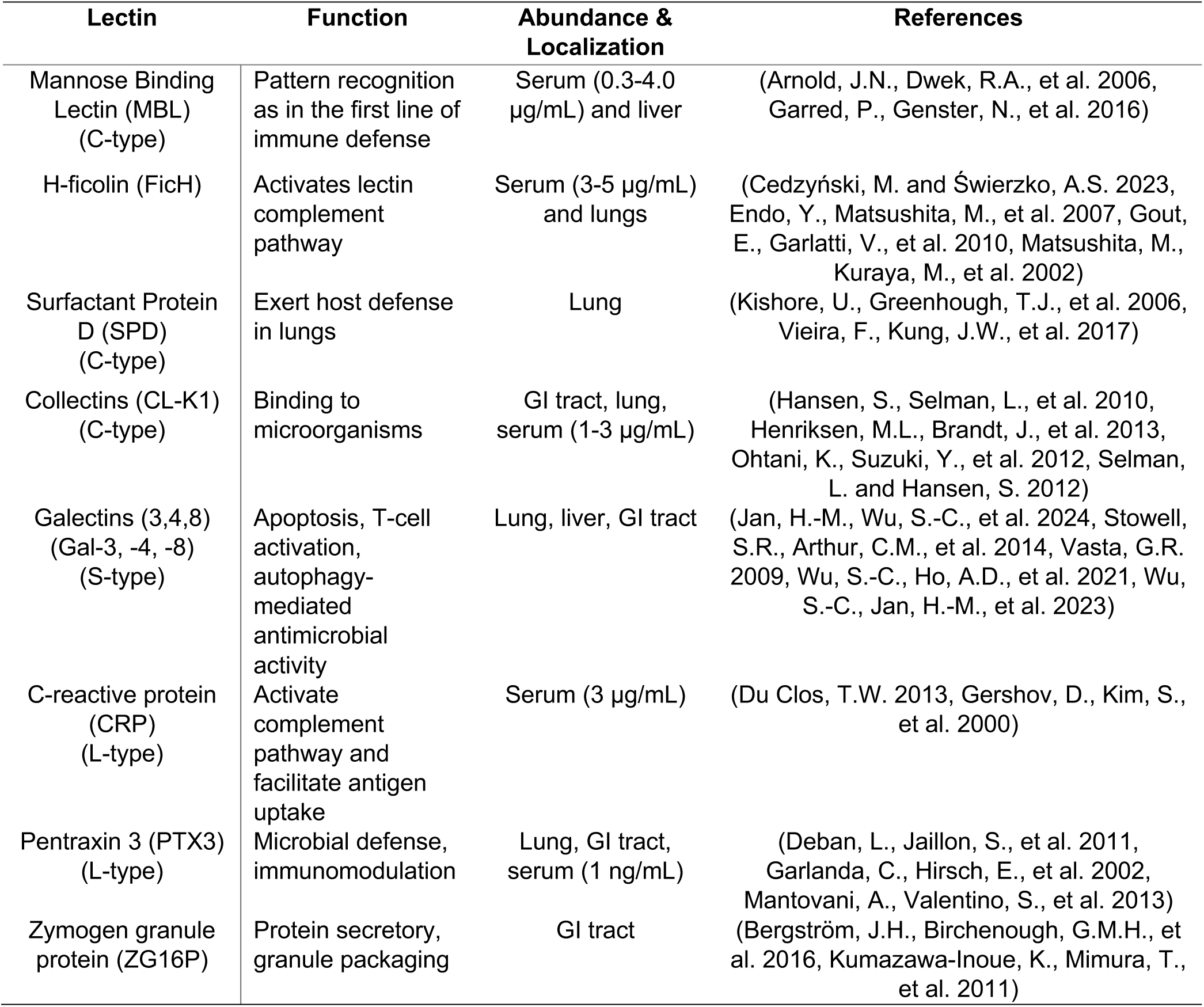
Biological significance and abundance of human soluble lectins chosen for GAP development in this study.

| Lectin | Function | Abundance & Localization | References |
| --- | --- | --- | --- |
| Mannose Binding Lectin (MBL) (C-type) | Pattern recognition as in the first line of immune defense | Serum (0.3-4.0 µg/mL) and liver | (Arnold, J.N., Dwek, R.A., et al. 2006, Garred, P., Genster, N., et al. 2016) |
| H-ficolin (FicH) | Activates lectin complement pathway | Serum (3-5 µg/mL) and lungs | (Cedzyński, M. and Świerzko, A.S. 2023, Endo, Y., Matsushita, M., et al. 2007, Gout, E., Garlatti, V., et al. 2010, Matsushita, M., Kuraya, M., et al. 2002) |
| Surfactant Protein D (SPD) (C-type) | Exert host defense in lungs | Lung | (Kishore, U., Greenhough, T.J., et al. 2006, Vieira, F., Kung, J.W., et al. 2017) |
| Collectins (CL-K1) (C-type) | Binding to microorganisms | GI tract, lung, serum (1-3 µg/mL) | (Hansen, S., Selman, L., et al. 2010, Henriksen, M.L., Brandt, J., et al. 2013, Ohtani, K., Suzuki, Y., et al. 2012, Selman, L. and Hansen, S. 2012) |
| Galectins (3,4,8) (Gal-3, -4, -8) (S-type) | Apoptosis, T-cell activation, autophagy-mediated antimicrobial activity | Lung, liver, GI tract | (Jan, H.-M., Wu, S.-C., et al. 2024, Stowell, S.R., Arthur, C.M., et al. 2014, Vasta, G.R. 2009, Wu, S.-C., Ho, A.D., et al. 2021, Wu, S.-C., Jan, H.-M., et al. 2023) |
| C-reactive protein (CRP) (L-type) | Activate complement pathway and facilitate antigen uptake | Serum (3 µg/mL) | (Du Clos, T.W. 2013, Gershov, D., Kim, S., et al. 2000) |
| Pentraxin 3 (PTX3) (L-type) | Microbial defense, immunomodulation | Lung, GI tract, serum (1 ng/mL) | (Deban, L., Jaillon, S., et al. 2011, Garlanda, C., Hirsch, E., et al. 2002, Mantovani, A., Valentino, S., et al. 2013) |
| Zymogen granule protein (ZG16P) | Protein secretory, granule packaging | GI tract | (Bergström, J.H., Birchenough, G.M.H., et al. 2016, Kumazawa-Inoue, K., Mimura, T., et al. 2011) |

The use of human lectins as glycan analysis probes has remained relatively limited due to technical challenges, including difficulties obtaining pure and natively-folded lectins (Martínez-Alarcón, D., Blanco-Labra, A., et al. 2018, Vorup-Jensen, T. 2014) and their propensity to form oligomeric structures (Sheriff, S., Chang, C.Y., et al. 1994). Thus, our objective was to generate tagged recombinant human lectins tailored for use in multiple applications. We have established robust modular approaches based on bacterial and mammalian cell expression systems emphasizing the following features: 1. Production and purification methods that yield natively-folded, bioactive lectins by extraction of recombinant protein from soluble cellular fractions, rather than renaturation from inclusion bodies; 2. Strategies for the site-specific introduction of reporter tags; 3. Flexibility to substitute native oligomerization domains with alternative scaffolds encoding known oligomerization properties.

Figure 1A illustrates the standardized workflow for developing, validating, and applying GAPs to detect glycans in a complex environment. The applications of the new GAPs include profiling co-existing microbial communities and microbial glycans, comparing the lectin binding specificities, and gaining comprehensive insights into lectin binding to other glycoproteins involved in host defense.

### Production of GAPs from recombinant lectins

#### Recombinant lectin production in bacteria

*Escherichia coli*-based expression was utilized to produce lectins that do not require glycosylation for folding and solubility (Table II). We designed constructs to generate the carbohydrate recognition domains (CRDs) alone or CRDs with oligomerization neck domains to preserve the native fold and binding valency. Additionally, for functionalization, the constructs were engineered with an N-terminal Strep-II tag or the sequence LPXTG for C-terminal sortase-mediated ligation (SML). The LPXTG motif can be treated with sortase for facile and site-specific conjugation of affinity tags (e.g., biotin) and spectroscopic labels (e.g., fluorophores) (Ghosh, S., Ahearn, C.P., et al. 2023, Policarpo, R.L., Kang, H., et al. 2014). The thermostability of purified GAPs was assessed using nano-differential scanning fluorimetry (nDSF) (Table II).

**Table 2:** Details of the glycan analysis probes (GAPs)

| Heterologous expression | Lectin | Domain | M.W (kDa) | pI | $E_{max}$ ($M^{-1}cm^{-1}$) | Expression system | Solubility | Yield (mg/L) | $T_m$ (° C) | Cumulant radius (nm) | Functional Tag |
| --- | --- | --- | --- | --- | --- | --- | --- | --- | --- | --- | --- |
| Mammalian | MBL | Full length | 25.2 | 5.4 | 23865 | HEK 293T | soluble | 2 | 74.8±0.04 | 6.9±0.09 | N-term Strep II |
|  | ΔN-SPD | Collagen-like domain+<br>+Neck<br>+CRD | 34 | 6.3 | 21220 | HEK 293T | soluble | 2 | 66.3±0.1 | 40.8±0.42 | N-term Strep II |
|  | CL-K1 | Full length | 27.6 | 5.2 | 25815 | HEK 293T | soluble | 2.5 | 45.3±0.6† | 17±0.40 | C-term Strep II |
|  | PTX3 | Full length | 41.6 | 4.9 | 65930 | HEK 293T | soluble |  | 58.2±0.01‡ | 12±0.05 | C-term Strep II |
|  | CRP | Full length | 24.5 | 5.5 | 50545 | HEK 293T | soluble | 16 | 60.2±0.01 | 4.5±0.25 | C-term Strep II |
| Bacterial | Gal3 | CRD | 16.6<br>3 | 9.8 | 9970 | BL21(DE3) | soluble | 18 | 59.8±0.1 | 3.3±0.45 | N-term Strep II |
|  | Gal4 | Full length | 38.3<br>6 | 9.3 | 42860 | BL21(DE3) | soluble | 9 | 65.0±0.1 | 3.5±0.63 | N-term Strep II |
|  | Gal8 | Full length | 38.2<br>5 | 8.0 | 32430 | BL21(DE3) | soluble | 12 | 53.2±0.2 | 3.8±0.52 | N-term Strep II |
|  | FicH | neck+<br>CRD | 22.2 | 6.2 | 52035 | BL21(DE3) | Partially soluble | 2.5 | 49.4±0.6 | 5.2±1.12 | Biotin by SML* |
|  | ZG16P | CRD | 17.6 | 9.3 | 29910 | BL21(DE3) | soluble | 6 | 54.6±0.8 | 1.8±0.32 | Biotin by SML* |

†CL-K1 showed a secondary $T_m$ at 60.6 ±0.6 °C
‡PTX3 showed a secondary and tertiary $T_m$ at 65.6 ±0.4 and 70.5±0.2 °C
\*SML: sortase-mediated ligation method

The GAPs produced using this approach facilitate the targeted identification and enrichment of complex glycans from natural sources in various biochemical and cellular assays. Additionally, incorporating a biotin tag allows the multivalent presentation of lectin CRDs on the tetrameric streptavidin scaffold, mimicking avidity effects observed in physiological conditions.

Using the *E. coli* expression system, we produced five new GAPs. One of the GAPs, the intestinal lectin ZG-16P, was generated to complement our previous studies of the oral protein ZG16B. Three of the GAPs are based on galectins (S-type lectins) (Klyosov, A.A., Witczak, Z.J., et al. 2008), an important family of human soluble lectins, which broadly recognize β-galactose-containing glycans and participate in diverse physiological and pathobiological roles (Vasta, G.R. 2009). The last member was based on a highly abundant blood lectin H-ficolin. The rationale for choosing these lectins is rooted in their localization, structural properties, and function.

Building on our previous studies on the oral lectin zymogen granule protein ZG16B (Ghosh, S., Ahearn, C.P., et al. 2023), we generated the paralog intestinal lectin ZG16P (Kanagawa, M., Satoh, T., et al. 2011, Kumazawa-Inoue, K., Mimura, T., et al. 2011) by recombinantly producing the lectin in *E. coli*. The construct encoding the ZG16P GAP excluded the signal sequence and contained both an additional C-terminal sortase tag (LPETG) and a Hisx6 tag (Supporting information, Figure S1C). This construct was expressed, purified, conjugated with biotin (Supporting information, Figure S1D), and subsequently used in binding studies to compare the oral and intestinal proteins.

We tested the ability of our platform to generate tailored galectins. The many roles of galectins include their ability to bind bacteria and mucins, and these intriguing proteins have been shown to bind both bacterial and mammalian glycans (Stowell, S.R., Arthur, C.M., et al. 2014, Wu, S.-C., Ho, A.D., et al. 2021). Unlike most human lectins, galectins are not glycosylated, allowing successful production in *E. coli.* The GAPs are based on CRDs from the chimeric galectin-3, and tandem-repeat galectins, including galectin-4 (Supporting information, Figure S1A) and galectin-8 (Supporting information, Figure S1B). The GAPs featured a C-terminal His6x tag for purification and an N-terminal Strep- II tag, enabling highly specific antibody-based detection in biological and biophysical analyses. The dual-tag constructs yielded pure functionalized lectin CRDs in a one-step metal ion-affinity chromatography-based enrichment. This procedure afforded three proteins from the family, which were further characterized.

The power of our bacterial expression system was highlighted by its ability to yield a GAP based on H-ficolin. This protein is one of the most abundant in serum without significant functional annotation (Matsushita, M., Kuraya, M., et al. 2002). Ficolins are trimeric lectins with an N-terminal collagen-like domain; the Gly-X-Y repeats are joined via a short neck domain to a glycan-binding fibrinogen-like domain, which serves as their CRD (Garlatti, V., Belloy, N., et al. 2007). Trimerization allows for multivalent binding, which enhances the recognition of cell-surface glycans. Despite considerable knowledge of the structure of the ficolins, their glycan specificity remains largely unknown, although some studies support N-acylated carbohydrates as potential ligands (Gout, E., Garlatti, V., et al. 2010).

Recombinant production of ficolins in *E. coli* has been challenging as they require glycosylation (Garlatti, V., Belloy, N., et al. 2007, Hummelshoj, T., Thielens, N.M., et al. 2007) and disulfide bond formation (Ohashi, T. and Erickson, H.P. 2004) for folding, stability, and function. Soluble fusion partners (Watson, A., Sørensen, G.L., et al. 2020) or coexpression with chaperones (Vázquez-Arias, A., Vázquez-Iglesias, L., et al. 2024) have been used to improve the expression and solubility of human lectins in *E. coli.* However, these strategies are often incompatible with producing functional probes for *in vitro* applications.

We successfully expressed an H-ficolin GAP construct with the H-ficolin neck and fibrinogen-like domain (amino acid residues 81-299), and additional C-terminal sortase (LPETG) and His6x tags in *E. coli*. The folded protein was purified from the soluble cellular fraction (Supporting information, Figure S1E) and conjugated with biotin using SML. (Supporting information, Figure S1F). The recombinant lectin oligomerized in solution as observed by the native gel electrophoresis (Supporting information, Figure S1G) and dynamic light scattering (DLS) (Table II, Supporting Information Figure S2) (Garlatti, V., Belloy, N., et al. 2007). The H-ficolin-GAP also showed a discrete folding transition in the nDSF assessment (Supporting information, Figure S3), and, despite the challenges of this protein class, the resulting multimer was sufficiently stable for biochemical studies. We also targeted bacterial production of GAPs using the neck and CRD domains of several representative lectins from the C-type or collectin class (Holmskov, U., Thiel, S., et al. 2003) and L-type or pentraxin class (Deban, L., Jaillon, S., et al. 2011). However, in these cases, the proteins did not retain a functional oligomeric fold, possibly due to non-native disulfide bond formation (Inforzato, A., Rivieccio, V., et al. 2008, Kishore, U., Greenhough, T.J., et al. 2006) and the lack of glycosylation on the N-terminal accessory domains, which influences stability and solubility (Garred, P., Genster, N., et al. 2016, Inforzato, A., Peri, G., et al. 2006, Tsuji, S., Uehori, J., et al. 2001, Wesener, D.A., Dugan, A., et al. 2017). To overcome these problems, a eukaryotic expression system was used for a subset of lectins that relied on posttranslational modifications for proper folding and function.

#### Recombinant lectin production in mammalian cells

The folding of many lectins depends on post-translational modifications that occur in eukaryotic cells. Specifically, lectins often require disulfide bonds, which can occur within their carbohydrate-binding domains or between domains to stabilize their oligomeric state. Collectins (members of the C-type lectin class) and pentraxins (L-type lectins) have multiple disulfide bonds that contribute to their stability. In addition, the collectins have collagenous domains that should require proline hydroxylation to promote folding. Finally, glycosylation can promote the folding of members of these two lectin classes. Thus, to ensure proper folding of GAPs based on collectins and L-type lectins (Supporting information, Figure S4), full-length lectins were expressed in an HEK-derived mammalian cell line (Table II). The lectins were engineered for secretion to facilitate isolation. An N-terminal Strep-II tag was added to the constructs for direct antibody-based detection. The formation of higher-order oligomers of the full-length recombinant lectins was evaluated by dynamic light scattering (DLS) measurements (Table II, Supporting information, Figure S5) and western blot analysis under non-reducing conditions. As with all of our recombinant lectins, their stability in solution was assessed using nDSF (Table II, Supporting information, Figure S6) to ensure a cooperative folding transition indicative of a properly folded protein.

We focused on three members of the human soluble collectin family (Håkansson, K. and Reid, K.B.M. 2000): mannose-binding lectins (MBL) and collectin kidney-1 (CL-K1), which are predominantly found in blood, and pulmonary lectin surfactant-associated protein D (SP-D), which is found in the lung and gut. These are all representative effector proteins of the innate immune system (Casals, C., García-Fojeda, B., et al. 2019, Cedzyński, M. and Świerzko, A.S. 2023).

Collectins share a common fold but show different extents of oligomerization. MBL and SPD form homo-oligomers of the basic trimer (Sheriff, S., Chang, C.Y., et al. 1994, Vieira, F., Kung, J.W., et al. 2017, Watson, A., Sørensen, G.L., et al. 2020). CL-K1 forms homo-oligomers of a trimeric subunit or hetero-oligomers with other related collectins (Henriksen, M.L., Brandt, J., et al. 2013). In our study, the representative N-terminal Strep-II tagged GAPs (Supporting information, Figure S4A-C) from these collectins also varied in their cumulant radii and multimerization as observed in DLS (Table II, Supporting information, Figure S5), DSF (Table II, Supporting information, Figure S6), and gel electrophoresis analysis (Supporting information, Figure S4A-C).

The L-type lectins or pentraxins are critical in the complement pathway-mediated pathway, a fundamental component of innate immunity (Deban, L., Jaillon, S., et al. 2011, Inforzato, A., Peri, G., et al. 2006). The pentraxins, which are abundant in the blood, enhance opsonophagocytosis (Mold, C., Gresham, H.D., et al. 2001), and tune inflammation (Du Clos, T.W. 2013, Gershov, D., Kim, S., et al. 2000). We generated the full-length short pentraxin C-reactive protein (CRP) (Supporting information, Figure S4D) and long pentraxin (PTX3) (Supporting information, Figure S4E) as representative L-type lectins for GAP development. Both are upregulated upon inflammation, with CRP functioning as a critical biomarker. We sought to evaluate their binding interactions as the ligand specificity of these lectins is not fully annotated.

Both CRP and PTX3-GAP showed multimer formation in the solution when assessed by gel electrophoresis under non-reducing conditions (Supporting information, Figure S4D, E) and DLS analysis (Table II, Supporting information, Figure S5). Notably, the long pentraxin PTX3, which has been predicted to adopt an octameric structure in solution (Inforzato, A., Peri, G., et al. 2006), showed a different oligomerization state relative to the short pentraxin CRP, which is reported to form a pentamer (Osmand, A.P., Friedenson, B., et al. 1977).

Successful production of GAPs that adopt native-like conformations and oligomerization states provide the means to investigate cognate, biologically relevant microbial glycan targets. Additionally, using the GAPs in various experimental formats (e.g., arrays, mucin-binding, and tissue staining) will reveal previously unrecognized glycosylation patterns, identify knowledge gaps in our understanding of glycobiology, and identify potential biomarkers for various conditions.

### Glycan-binding specificity of GAPs

#### GAP analysis with bacterial glycan microarrays

With access to GAPs that adopt native-like conformations and oligomerization states, we next assessed their glycan-binding specificities. Glycan arrays enable the detection of a range of potential glycan ligands with limited protein quantities and represent a powerful high-throughput method for identifying glycan-binding specificity. For the array analysis, we deployed both microbial and human glycan arrays. To determine the ligand specificity of the functionalized lectin CRDs, we first used a custom bacterial glycan array from the Scripps Research Institute Microarray Core Facility, printed with distinct polysaccharides isolated from a range of microbes (Stowell, S.R., Arthur, C.M., et al. 2014). The bacterial glycan array includes over 300 microbial polysaccharides printed on an amine-reactive N-hydroxysuccinimidyl ester (NHS) activated glass slide in a 4*12 grid matrix, each grid having six replicates of each glycan. The array features glycans primarily isolated from gram-negative bacteria belonging to the order *Enterobacteriales* (Stowell, S.R., Arthur, C.M., et al. 2014), including the genera *Escherichia, Klebsiella, Proteus, Providencia, Salmonella, Shigella,* and *Yersinia*. Capsular polysaccharides derived from the gram-positive bacteria *Streptococcus pneumoniae* are also included in the microarray. The printed polysaccharides are enriched in non-mammalian monosaccharides, such as FucNAc, D-FucNAc, ManNAc, 3’-deoxy sugars (KO and KDO), and heptoses. A similar bacterial glycan array has previously been used to predict and validate host-microbe interactions. For example, the binding of Galectin-8 to immobilized glycan epitopes derived from the gram-positive microbe *Streptococcus pneumoniae* predicted strain-specific binding and antimicrobial activity against whole microbial cells, supporting that the array could provide insight into physiologically relevant glycan-binding properties (Wu, S.-C., Jan, H.-M., et al. 2023). A notable limitation of the bacterial glycan array is that it is generated using a limited set of glycans isolated from natural resources and lacks glycans isolated from species other than pathogenic bacteria. Importantly, it does not encompass the diversity of monosaccharide building blocks or glycoside linkages of the constituent glycans found in microbial glycomes that can be achievable through combined synthetic strategies and extraction from natural resources (Geissner, A., Reinhardt, A., et al. 2019). We evaluated the binding of soluble human lectins from the GAP production platform outlined previously on the bacterial glycan array. These included: S-type lectins (Galectin- 3, -4, -8), pentraxins (CRP, PTX3), collectins (MBL, SPD, CL-K1), as well as H-ficolin and ZG16P. The binding intensity of the GAPs to each glycan sample was calculated as an average of integrated spot intensities from six replicates. Internal grid controls, including dye samples and printed null spots, aided in validating the consistency of the assay results.

Due to the lot-to-lot variability in the construction of arrays and fluorescence readings across experiments, we normalized binding data for each lectin such that the highest binding value above our binding signal threshold was set to 100%, and binding to every glycan was plotted as a percentage of binding relative to this value (Figure 2A, Supporting information, Spreadsheet-1). We set the top 10% of the highest normalized fluorescence intensity (100%) as the threshold to identify binders of GAPs on the bacterial glycan array (Supporting information, Spreadsheet 1) (Cao, Y., Park, S.-J., et al. 2019). This approach enabled independent determination of the binders for each GAP, even when the binding strength was relatively weak. Additionally, the total fluorescence-binding intensity is considered to assess the binding profile of GAPs for one-to-one comparison (Figure 2B, Spreadsheet-2).

**Figure 2:**
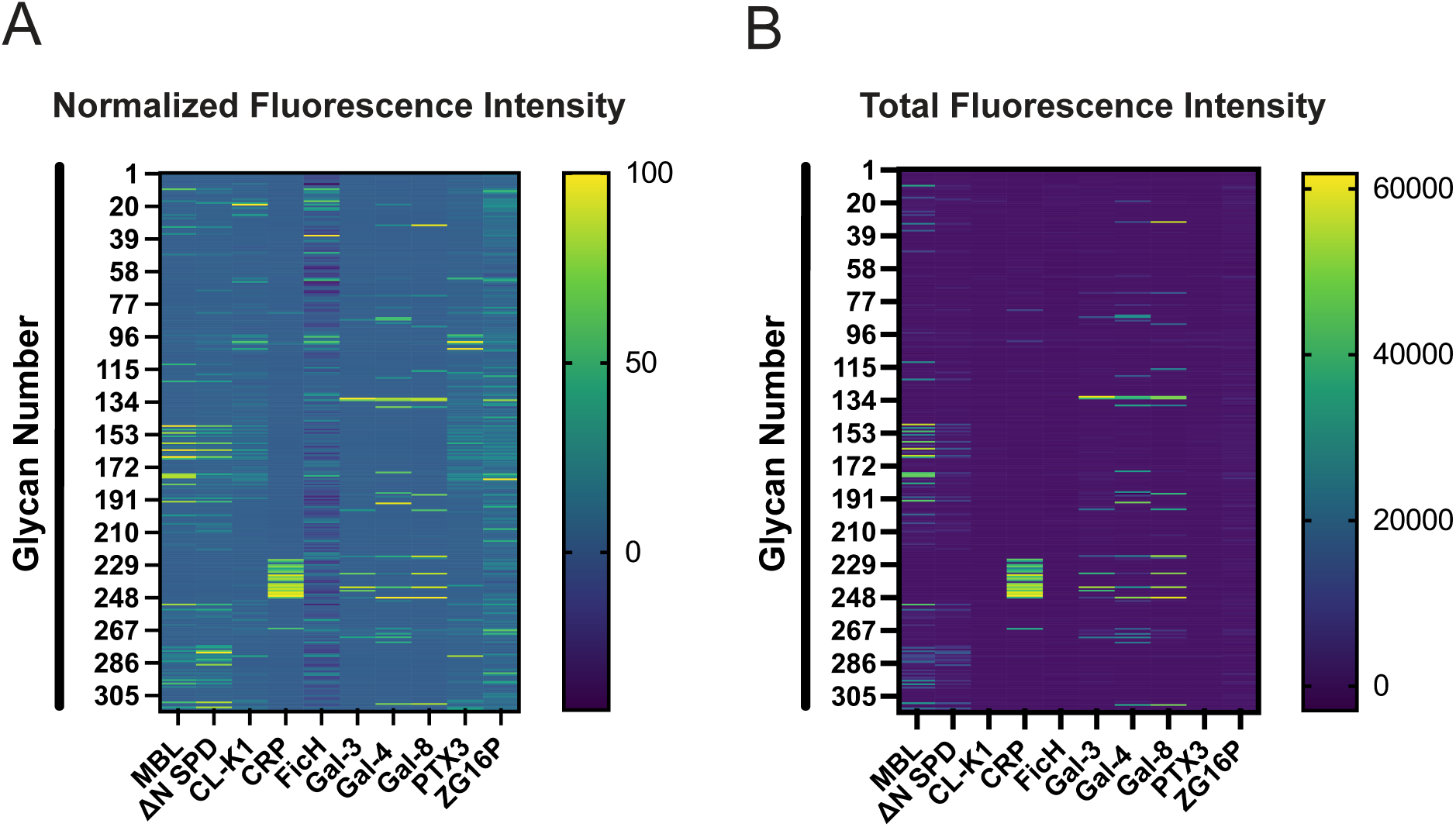
Microbial glycan binding specificity of GAPs: Glycan binding profile of GAPs on custom bacterial glycan array (from TSRI MicroArray Core Facility, The Scripps Research Institute) represented as heatmaps. (A) Heatmap generated based on the normalized values of relative fluorescence intensity for each lectin considering the highest binding value as 100% binding (on the scale). Data are presented as the mean ± s.d. (*n*=6 technical replicates). (B) Heatmap generated based on the total binding intensity quantified as relative fluorescence unit (on the scale).

We observed distinct binding profiles for the GAPs on the bacterial glycan arrays (Figure 2A, B, Supporting information, Figure S7), supporting the specificity of the GAP-glycan interactions. Several GAPs, including MBL, SPD, CRP, Galectin-3, -4, -8 exhibited more readily apparent binding to multiple microbial glycans than other GAPs (Figure 2B, Supporting information, Spreadsheet-2). We also noted that C-type lectins, such as MBL and SP-D, preferred similar microbial glycans, including the lipopolysaccharides (LPS) from pathogenic strains of gram-negative *Yersinia pestis* containing heptose KO and KDO (Supporting information, Figure S7, Spreadsheet 1, 3). These lectins also showed a broad range of binding to O-specific polysaccharides (OPS) or O-antigens from other pathogenic strains of *Shigella flexneri*, *Escherichia coli,* and *Klebsiella pneumoniae* (Supporting information, Figure S7, Spreadsheet 1). In contrast, CRP, a L-type lectin, specifically interacts with capsular polysaccharides (CPS) from *Streptococcus pneumoniae*. Galectin-3, 4, and 8 bound a diverse set of glycans but also shared similarities in their binding profiles, showing a strong preference for OPS and LPS containing colitose, D-glucofuranose, and D-hexose (Supporting information, Figure S7, Spreadsheet 1, 3).

GAPs developed from CL-K1, H-Ficolin, PTX3, and ZG16P exhibited weaker signals on the array, likely due to the absence of their cognate targets on the bacterial glycan array (Figure 2B). This result highlights the need for screening the GAPs on more diverse glycan arrays. Despite the need for arrays that are more representative of the diversity of microbial glycans, even GAPs with weak microbial glycan binding revealed intriguing profiles in the normalized analysis (Figure 2A). For example, the long pentraxin PTX3 has CRD that is structurally similar to that of CRP, yet the binding profiles differ; PTX3 bound to a broad range of glycans with a preference for OPS from *Proteus mirabillis* and *Shigella flexneri* containing ethanolamine-modified sugars (Supporting information, Figure S7, Spreadsheet 1, 3) (Toukach, F.V., Arbatsky, N.P., et al. 2001). We mainly detect H-ficolin binding to OPS from different O-serotypes of *Proteus mirabillis* containing a D-galacturonic or D-glucuronic acid, often attached to an L-amino acid via an amide bond (Supporting information, Figure S7, Spreadsheets 1 and 3) (Arbatsky, N.P., Shashkov, A.S., et al. 1999).

The binding profiles of the monomeric intestinal lectin ZG16P were broad, although we speculate that many may be non-specific. Previously, ZG16P was reported to bind to several commensal gram-positive bacteria, including *Lactobacillus jensenii and Enterococcus faecalis* (Bergström, J.H., Birchenough, G.M.H., et al. 2016). The related oral lectin ZG16B binds to a set of oral gram-positive bacteria, including *Streptococcus vestibularis* (Ghosh, S., Ahearn, C.P., et al. 2023). Therefore, with a limited set of glycans from commensal bacteria on the array, the physiologically relevant glycan targets of ZG16P remain unclear.

#### GAP analysis with mammalian glycan microarrays

To complement the bacterial glycan array analysis, we used the commercially available mammalian glycan array 300 (Ray Biotech). The array includes 300 mammalian glycans comprising N-glycans, glycolipid glycans, human milk oligosaccharides, and sialic acid-rich tandem epitopes. Although the glycan array provides sufficient diversity for determining lectin binding, the limited availability of O-linked glycans and glycosaminoglycans (GAGs) makes it difficult to completely rule out the possibility that some GAPs may bind to both host glycans and microbial targets. Indeed, we have previously observed simultaneous binding to both highly glycosylated mucins and bacteria (Ghosh, S., Ahearn, C.P., et al. 2023).

In general, GAPs exhibited a weaker affinity for mammalian glycans compared to microbial glycans (Supporting information Figure S8, Spreadsheet 4). This selectivity highlights the propensity of soluble human lectins to bind microbial glycans. However, various GAPs, particularly those based on galectins, bound selected mammalian glycans (Supporting information Figure S8, Spreadsheet 4 and 5). This observation is consistent with the multiple functional roles of galectins (Bergström, J.H., Birchenough, G.M.H., et al. 2016, Ghosh, S., Ahearn, C.P., et al. 2023).

#### GAP binding to human host mucins

We also explored GAP binding to other host glycans in physiologically relevant niches. This approach aims to provide insights into the role of lectins in host-microbe interactions. The mucosal barrier is central to regulatory activities in host-microorganism interactions, including receptor-mediated endocytosis, cellular recognition, and microbial adhesion (Frenkel, E.S. and Ribbeck, K. 2015, Jeffers, F., Fuell, C., et al. 2010). As a component of mucosal immunity, some lectins can interact with the highly glycosylated mucins, the glycoproteins with O-linked glycans in the mucus layer (Peiffer, A.L., Dugan, A.E., et al. 2024). Mucins can modulate inflammatory responses by forming mucus layers on epithelial cell surfaces that protect tissues from bacterial invasion (Frenkel, E.S. and Ribbeck, K. 2015). The crosstalk of soluble lectins and mucins often influences microbial colonization and the bacterial growth regulatory effects of the mucosal barrier.

In recent studies, Galectin-3 has been shown to bind the gastrointestinal mucin MUC2 and respiratory tract mucins MUC5AC and MUC5B (Diehl, R.C., Chorghade, R.S., et al. 2024) when regulating innate immune properties of airway mucosal surface; ZG16P prevents bacterial aggregation on colon epithelial cells by working together with the colon mucin MUC2 (Bergström, J.H., Birchenough, G.M.H., et al. 2016); the oral lectin ZG16B recruits oral mucin MUC7 on opportunistic pathogens in the oral cavity and maintains microbial homeostasis (Ghosh, S., Ahearn, C.P., et al. 2023). However, investigating lectin-mucin interactions *in vitro* is challenging due to the high complexity of O-glycosylated mucins present and the multivalent, cooperative nature of these interactions. The GAPs represent physiologically relevant lectin-based analytical tools to address these challenges. We employed biotinylated or Strep-II-conjugated GAPs to capture the glycan-specific interaction between mammalian mucins and human soluble lectins. GAPs can be used to identify specific glycan epitopes on the mucins that can modulate the mucosal barrier properties, influencing microbial adhesion, colonization, and potentially pathogenicity. Additionally, the biotinylated GAPs can also be used to create the multivalent display required for these interactions, mimicking aspects critical for physiological binding.

GAPs produced from *E. coli* were screened for their interaction with crude porcine mucins or extracted porcine mucins MUC2, MUC5AC, and MUC5B by a dot-blot assay. The relative binding of GAPs to different mucins was normalized to the highest intensity integrated from the dot blot signals. GAPs from Galectin-4 (Gal-4) and -8 (Gal-8) bound strongly to MUC2, MUC5AC and MUC5B (Figure 3A, B, Supporting information, Figure S9A, B). The binding of Gal-8 was inhibited by the addition of 50 mM lactose as a competitive inhibitor, indicating that the interaction is mediated through mucin glycan recognition (Carlsson, S., Öberg, C.T., et al. 2007). However, the interaction of Gal-4 with the mucins was not disrupted by lactose, likely due to the stronger affinity of Gal-4 for the multivalent N-acetyllactosamine (LacNAc) repeats present in the colonic mucins relative to lactose, as shown previously (Elzinga, J., Narimatsu, Y., et al. 2024, Holmén Larsson, J.M., Karlsson, H., et al. 2009).

**Figure 3:**
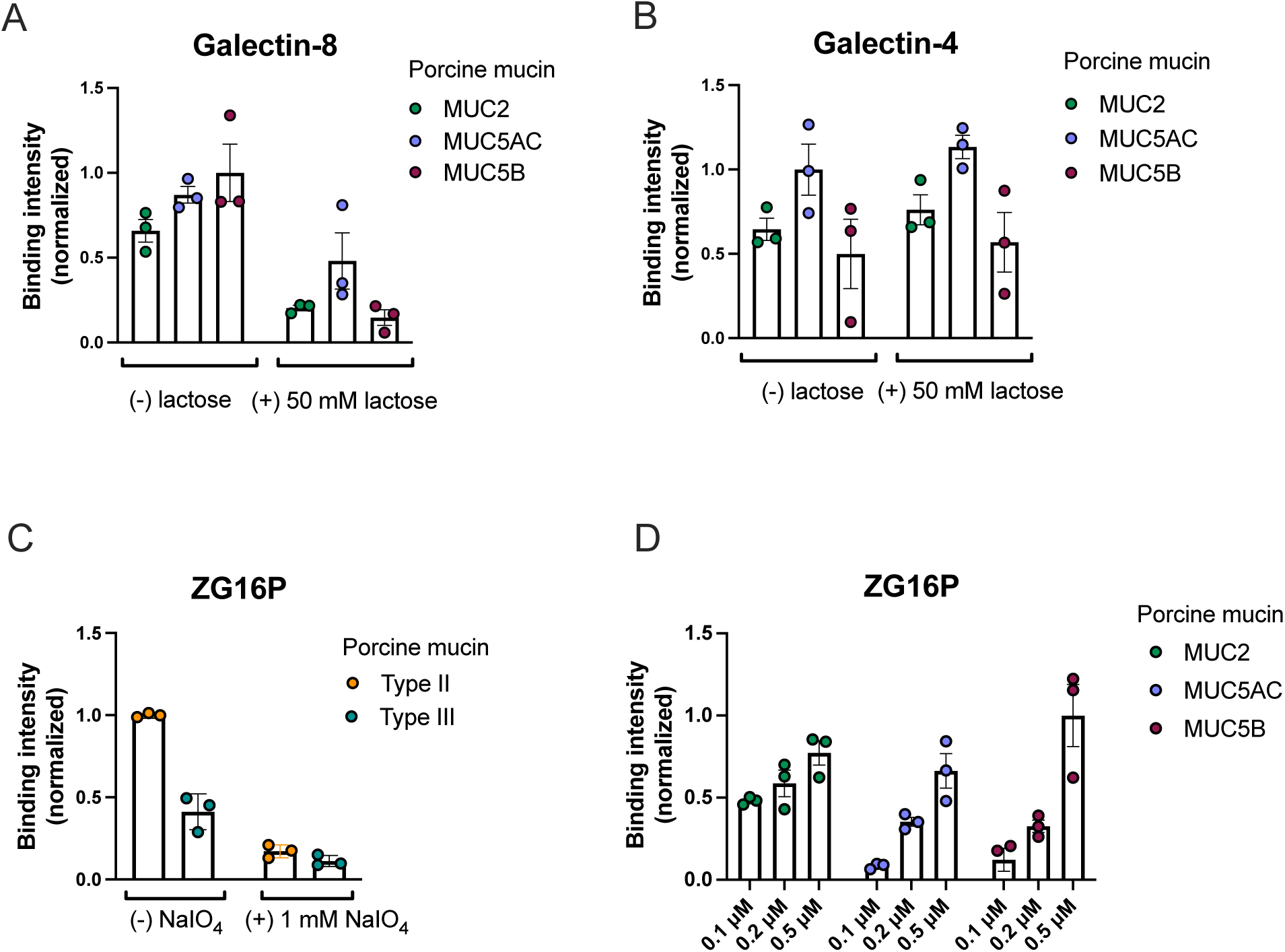
Mucin binding of GAPs: Binding profile of (A) Galectin-8 GAP and (B) Galectin- 4 GAP with purified porcine mucins MUC2, MUC5AC, and MUC5B. The binding intensity was quantified from the mucin dot blots probed with Strep-II tagged GAPs at 250 nM concentration, followed by washing with mucin binding buffer with or without 50 mM lactose as a competitive inhibitor. (C) Binding of ZG16P-biotin GAPs at a 100 nM concentration to crude porcine mucins, treated with 1 mM sodium periodate to oxidize the vicinal-diols of mucin O-glycans. Data are represented as mean± SEM. (*n*=3 technical replicates). (D) Binding of ZG16P-biotin GAP to purified porcine mucins MUC2, MUC5AC, and MUC5B at different concentrations. Data are represented as mean± SEM. (*n*=3 technical replicates).

The ability of GAPs to uniquely bind different mammalian mucins was assessed for ZG16P and H-ficolin. As these lectins are not yet known to bind mucin glycans, we initially tested them on commercially available porcine Type-II and Type-III mucins. The ZG16P- GAP showed binding to the crude porcine mucins (Figure 3C, Supporting information, Figure S9C). The lectin binding was reduced when mucins were treated with 1 mM sodium periodate, suggesting that mucin O-glycans mediated the binding. When screened for binding to purified mucins, ZG16P showed binding to MUC2, MUC5AC, and MUC5B in a concentration-dependent manner (Figure 3D, Supporting information, Figure S9D). In contrast, the GAP from H-ficolin did not show glycan-dependent binding to crude porcine mucins (Supporting information, Figure S10A, B) or concentration-dependent binding to the purified porcine mucins (Supporting information, Figure S10C, D). These differences in mucin binding are consistent with the localization of the lectins: H-ficolin is typically found in the blood, while ZG16P is present in the gut. Thus, only the latter lectin resides at a mucosal barrier. Similar reasoning can be applied to rationalize the binding difference between the galectins and ZG16P. The galectins showed stronger binding to airway pathway mucins (MUC5AC and MUC5B), while the latter showed a preference for intestinal mucin (MUC2). The O-glycan structures in MUC2, MUC5AC, and MUC5B differ in overall complexity and length of their glycan chains. MUC2 (Thomsson, K.A., Holmén-Larsson, J.M., et al. 2012) typically has shorter, less extended O-glycans than the longer and more complex structures found on MUC5AC and MUC5B (Holmén , J.M., Karlsson, N.G., et al. 2004). This difference in glycosylation pattern can contribute to the distinct and varied mucin-binding capabilities of GAPs and underscore the specialized roles of different lectins in mucin recognition and interaction.

#### Application of GAPs in tissue analysis

The GAPs also serve as tools for tissue staining. They can detect mucins or bacteria in relevant physiological contexts. In addition, they can be used to probe changes in microbial communities or altered glycosylation of the mucus layer, which is common in various colonic and respiratory conditions (Chatterjee, M., Putten, J.P.M.v., et al. 2020, Grondin, J.A., Kwon, Y.H., et al. 2020). Therefore, GAPs can reveal new aspects of the organization of mixed host and microbial cells and as non-invasive agents for differentiating diseased from healthy tissues.

We used a FITC-conjugated Galectin-3 (Gal-3) GAP for fluorescent lectin histochemistry (Figure 4). Fixed murine ileum and colon tissues were sectioned, mounted on a glass slide, and incubated with FITC-conjugated Gal-3 GAP and the 4’,6-diamidino-2- phenylindole (DAPI) stain was used as the nuclear counterstain for the tissue section. The Gal-3 GAP readily stained the extracellular matrices of mucus-producing goblet cells in murine intestinal tissue. The goblet cells form the inner mucus layer rich in MUC2 that separates the colonizing bacteria in the outer mucus membrane from the epithelium (Kim, Y.S. and Ho, S.B. 2010, Pelaseyed, T., Bergström, J.H., et al. 2014). Mice with a disrupted inner mucus layer, lacking mucin MUC2, showed symptoms of colitis due to penetration of bacteria from the outer to the inner mucus layer (Johansson, M.E., Phillipson, M., et al. 2008, Van der Sluis, M., De Koning, B.A., et al. 2006). These observations suggest that the Gal-3 GAP may recognize the mucin glycans in the inner mucus scaffold on goblet cells, which can effectively reveal the organization of the intestinal tissues (Diehl, R.C., Chorghade, R.S., et al. 2024).

**Figure 4:**
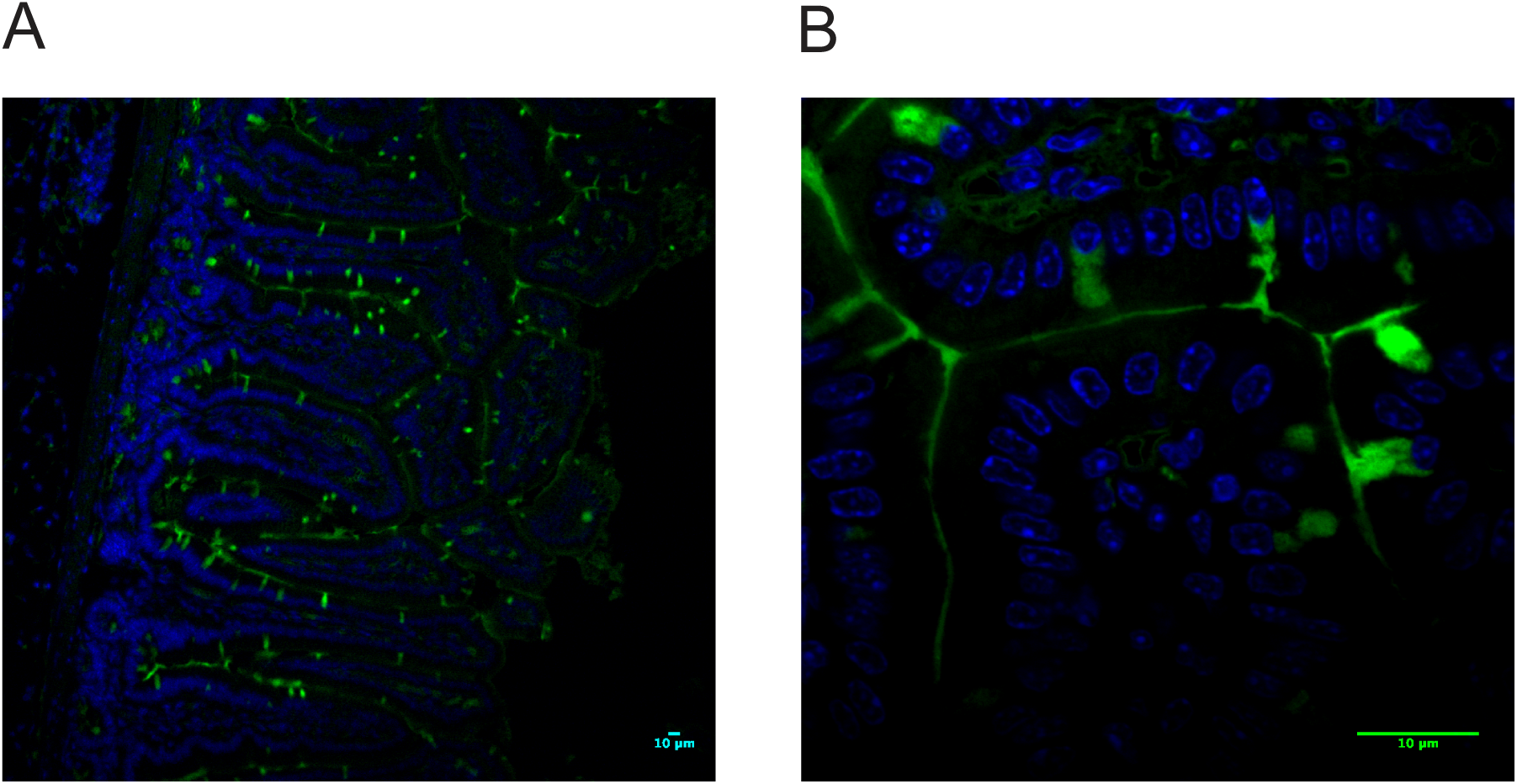
Staining of the extracellular matrices of mucus-producing goblet cells in murine intestinal tissue by FITC conjugated Galectin-3 GAP. Surgical murine ileum and colon tissue were probed with Galectin-3-FITC (shown in green) and visualized with the RPI Spinning disc confocal microscope at (A) 10x and (B) 63x objectives. The GAP stained the inner mucus layer produced by the goblet cells. Nuclei are stained with DAPI (blue).

## Discussion

A significant current challenge in glycoscience is elucidating the details of microbial cell wall glycoconjugates that mediate host cell adhesion and drive pathogenesis. Plant and fungal lectins (Rüdiger, H. and Gabius, H.-J. 2001, Tsaneva, M. and Van Damme, E.J.M. 2020) have been widely used to characterize mammalian glycans. However, due to the greatly expanded diversity of bacterial glycomes (Imperiali, B. 2019), many complementary tools are needed to reliably detect and characterize physiologically relevant glycan epitopes on the microbial glycocalyx and effectively distinguish them from host glycans.

Human lectins, with their unique capacity to recognize distinct glycan epitopes, especially within diverse and complex microbiomes, offer a potential solution to the limitations of current lectin-based approaches. Motivated by this insight, our strategy of developing glycan analysis probes (GAPs) from human soluble lectins establishes a foundational toolset for advancing our understanding of the precise protein-glycan interactions in host-microbe communication.

The development of GAPs based on human lectins will enhance our understanding of the characteristics of human soluble lectins across different structural classes, tissue-specific expression, roles in immune responses, and adaptive divergence in molecular recognition for diverse glycans. The presented experimental strategy is also generalizable, making it amenable to expansion to other lectins or glycan-binding proteins. Furthermore, we demonstrate that the engineered lectin-based probes are isolated in their correct folded and oligomeric states. The ability to produce natively folded GAPs and precise functionalization for glycan detection marks a practical advancement in studying complex biological interactions based on bacterial glycoconjugates. Critically, GAPs offer a new platform for identifying pathogens and defining clinically significant glycan epitopes for antibiotic development.

One key advantage of using GAPs in decoding host-microbe interactions is the potential to perform unbiased searches for ligand specificity at a high throughput scale. For example, GAPs developed from human lectins ZG16B, MBL, and Intelectin 1 (ITLN1) have been used to recognize and isolate the microbial targets of the lectin from complex microbiota (Ghosh, S., Ahearn, C.P., et al. 2023, McPherson, R.L., Isabella, C.R., et al. 2023). Furthermore, the GAP based on ZG16B could precisely identify cell wall polysaccharides on the surface of oral commensals that mediate specific lectin-microbe interactions (Ghosh, S., Ahearn, C.P., et al. 2023). Therefore, GAP applications provide top-down approaches for glycan identification, facilitating analysis of glycan binding across different species and in unfractionated microbiome samples.

GAPs offer a complementary approach for glycan profiling with the recently developed human lectin-based arrays (Benjamin, S.V., Jégouzo, S.A.F., et al. 2024). These arrays have proven crucial in high throughput screening of lectin interactions with isolated microbial and fungal polysaccharides, as well as viral glycoproteins, revealing unique binding targets of the lectins. The approaches share some lectin representatives, such as MBL, SP-D and galectins, but provide distinct insights: while human lectin arrays reveal the overall glycosylation profile of a sample, GAPs identify specific glycan structures in heterogeneous samples. Therefore, by combining GAPs with human lectin arrays, researchers can achieve a comprehensive understanding of lectin specificity across a wide range of microbial and mammalian glycans in complex systems.

In this study, we deployed the GAPs on bacterial glycan arrays to identify specific microbial targets from a bottom-up approach. Additionally, we used them on mammalian glycan arrays to understand the cross-reactivity of human lectins across diverse glycomes. Glycan array screening is important to ensure that the glycosylation of the GAPs does not confound recombinant GAP recognition. This issue is especially relevant for GAPs produced in mammalian cell culture, as they are secreted proteins and, therefore, glycosylated. Despite these concerns, we have observed that the selectivity of the different GAPs on glycan arrays indicates that their binding is driven by recognition of the immobilized glycans and not merely by protein glycosylation.

The custom bacterial glycan array used in this study features more than 300 naturally isolated microbial glycans comprising O-antigens from pathogenic gram-negative bacteria and cellular polysaccharides from selected gram-positive bacteria. Given the diversity of microbial glycans (Imperiali, B. 2019), even these valuable arrays can only represent a small fraction of the available glycans. However, despite the relatively low diversity score of the custom bacterial glycan array relative to other reported arrays (Geissner, A., Reinhardt, A., et al. 2019), several GAPs still displayed potential binding candidates, underscoring the effectiveness of the probes in targeting specific glycans even in less diverse environments. Additionally, GAPs can distinguish between various glycan structures, implying their pathobiological significance of human lectins in identifying a specific set of microbes in a complex microbiome. More diverse glycan arrays, comprising glycans from commensal and pathogenic bacteria, will be very valuable for determining the ligand binding specificity of other GAPs and, importantly, for identifying differentiating characteristics between commensals and pathogens.

Screening GAPs on bacterial and mammalian glycan arrays enabled an extensive cross-reactivity assessment, a critical consideration in the contexts of host-microbe interactions. Furthermore, testing GAPs for mucin binding illustrated their utility in identifying host glycan signatures. This screening allowed for the prediction of interactions across various glycan types in complex biological environments where microbial and host glycans coexist, such as the oral cavity and digestive tract. Additionally, GAP-mediated histochemistry provided a potential diagnostic and prognostic alternative to immunostaining in differentiating diseased vs healthy tissues.

Collectively, GAPs are valuable resources to aid in therapeutic interventions. GAPs can be beneficial in building a comprehensive database of glycan epitopes involved in host-microbe interactions. The curated databases, combined with machine learning (Bojar, D., Meche, L., et al. 2022), can predict unknown glycan-mediated interactions in host-microbe dynamics, advancing the development of glycoconjugate-based therapeutics. Additionally, GAPs can serve as a platform for developing peptide-based inhibitors or small molecules that disrupt native lectin-glycan interactions, offering a promising strategy to block pathogen adhesion and infection.

## Materials and Methods

The recombinant production and characterization of GAPs and screening of GAPs on mammalian glycan arrays are discussed in detail in the supporting information text.

## Screening of GAPs on custom bacterial glycan array

The slide printed with microbial glycans (from TSRI MicroArray Core Facility, The Scripps Research Institute) was equilibrated at room temperature after taking out of -20 ^0^C for 20- 30 minutes. The slide was placed on a slide holder tube and soaked in wash buffer (20 mM HEPES; pH7.5, 150 mM NaCl, 10 mM CaCl2) for 10 minutes. The slide was placed on a raised slide holder platform in a humidification chamber with wet paper towels and the printed area (no barcode side) was sealed by placing a rubber guard around it. 800 µL of Strep-II tagged lectin, diluted to 5µg/mL (for Galectin-3,4,8), 10µg/mL (for full-length MBL, and N-terminal truncated SPD) or 50 µg/mL (for full-length CL-K1 and PTX3) in protein binding buffer (20 mM HEPES; pH7.5, 150 mM NaCl, 10 mM CaCl2, 0.1% BSA, 0.1% Tween 20) was added on the slide and incubated for 1 h at room temperature with gentle rocking inside the humidification chamber. Biotinylated ZG16P (800 µL) at a concentration of 10 µg/mL (either free or pre-complexed with streptavidin-Cy3) in protein binding buffer (20 mM HEPES; pH 7.5, 150 mM NaCl, 10 mM CaCl2, 0.1% BSA, 0.1% Tween 20) was added to the slides. Meanwhile, the Dy549-conjugated strep-monoclonal antibody (StrepMAB) was prepared by diluting the stock solution (from IBA-Lifesciences) of 0.5 mg/mL in 750 µL protein binding buffer at a final concentration of 4ug/mL. For biotinylated GAPs, the Streptavidin conjugated Cy3 (RayBiotech 300 glycan array kit) was prepared according to the manufacturer’s protocol. After the incubation, the slide was washed in protein binding buffer, wash buffer, and double-deionized water consecutively, four times in each wash medium. The slide was placed back in the chamber, sealed with the rubber guard and 750 µL of Dy549-conjugated strep-monoclonal antibody was added to the slide and incubated for 1 h at room temperature with gentle rocking. The slide was then washed in protein binding buffer, wash buffer consecutively, four times in each wash medium, and lastly in double-deionized water, thrice each time in two separate wash containers. The slide was dried in a slide spinner and scanned at the Genepix4000 laser scanner using the 532 nm laser.

The data obtained was analyzed by using Microsoft Excel to average the fluorescence intensities of the best four of six replicates for each glycan. The average intensity was plotted against the sample ID number.

## Mucin binding assay

Lyophilized MUC2, MUC5AC, and MUC5B were purified from porcine sources as previously described and were generously provided by Prof. Katharina Ribbeck. These mucins were reconstituted to 10 μg/μL overnight at 4 °C in mucin assay buffer (20 mM HEPES, 150 mM sodium chloride, pH 7.4) as previously described, then diluted to a concentration of 1 μg/μL in the same buffer. The porcine mucins type II and type III (Sigma) were diluted at the same concentration and treated with 1 mM sodium periodate at 37°C overnight, protected from light. One µg of the mucin samples was dotted on nitrocellulose or PVDF membranes in triplicate. The membranes were incubated with biotinylated or Strep-II-tagged GAPs at appropriate concentrations. For galectins, membranes were washed with mucin assay buffer supplemented with 50 mM lactose after incubating with GAPs. The blots were visualized with appropriate detection methods at different conditions (SI text) and analyzed using the Fiji image suite.

## Fluorescence Immunohistochemistry using GAP

Surgical tissue of mouse ileum and colon, fixed in Carnoy’s fixative, were paraffin-embedded and sectioned (4–5 μM) and mounted on X-tra™ positive-charged slides (Leica Biosystems, Buffalo Grove, IL). Sections were deparaffinized, rehydrated in gradient ethanol, H2O, and blocked 5% BSA (in 1 × PBS) for 60 min before overnight incubation at 4°C with FITC conjugated Galectin-3 GAP. Following overnight incubation with GAP, samples were counter-stained with DAPI (10 µg/mL) and mounted with ProLong Diamond antifade mountant (Thermo) with a 24 hour cure time. Images were captured using RPI spinning disk confocal microscope.

## Acknowledgments

We thank Dr. Ryan McBride for providing the Custom Bacterial Glycan Arrays. We thank Dr. Shanu Mehta of the Nanowell Cytometry Core Facility at the Koch Institute for technical support with Genepix microarray scanner. We thank Prof. Katharina Ribbeck, Dept. of Biological Engineering, MIT for generously providing us with MUC2, MUC5AC, and MUC5B enriched samples.

