## Supplementary Information for "Glycan analysis probes inspired by human lectins for investigating host-microbe crosstalk"

Supporting information text and figures include:

- Materials and methods
- Figure S1 to S10

#### Materials and Methods

**Cell line and strains:** BL21(DE3) competent cells were obtained from Agilent. HEK293D cells were purchased from ATCC.

**Plasmids for bacterial expression of mGAPs:**

*Plasmid for galectins:*

The coding sequence of human galectin-3 residues 113-250 (UniProt: P17931), and full-length galectin-4 (UniProt: P56470) and -8 (UniProt: O00214) were obtained as gene products from Integrated DNA Technologies (Coralville, IA) and inserted into a pET24a vector (Novagen, Madison, WI) by the Gibson assembly method. This vector was amplified in *Escherichia coli* DH5α cells, then used to transform DE3 Tuner cells for protein expression.

*G-block DNA assembly of ZG16P and* H-ficolin *constructs in pET24a:* The codon-optimized constructs for BL21(DE3) expression of human zymogen granule protein 16 homolog P (ZG16P) (UniProt: O60844) and H-ficolin (UniProt: O75636) from *H. sapiens* were obtained from IDT as synthetic gBlocks. Residues 21-159 of ZG16P comprising the CRD and C-terminal LPETG linker and 6x Histidine tag were subcloned into pET24a between BglII and XhoI sites. Similarly, residues 133-299 of H-ficolin comprising neck domain, CRD, C-terminal LPETG linker, and 6x Histidine tag were subcloned into pET24a between BglII and XhoI sites.

**Plasmid construction for mammalian expression of mGAPs:**

Human MBL (UniProt: P11226), ∆N-SPD (construct excludes 24-45 amino acid residues) (UniProt: P35247), CL-K1(UniProt: Q9BWP8), CRP(UniProt: P02741) and PTX3 (Uniprot: P26022) gBlocks DNA Fragment (Integrated DNA Technologies) was Gibson ligated into a linearized pcDNA4 plasmid [primers reverse (5′-GGTGAAGCTTAAGTTTAAACG-3′) and forward (5′-TGAGGATCCACTAGTCCAGTG-3′)]. StrepII tag (5′-TGGAGCCATCCGCAGTTTGAAAAG-3′) was inserted C-terminal of amino acid 20 into pcDNA4-hMBL2 using inverse polymerase chain reaction (PCR) mutagenesis using forward primer (5′-TGGAGCCATCCGCAGTTTGAAAAGGAAACTGTGACCTGTGAGG-3′) and reverse primer (5′-TTCTGAGTAAGACGCTGCC-3′). Correct insertion was verified by DNA sequencing (Quintara Biosciences).

**Expression and purification of galectins:**

Galectin-3, -4 and -8 were expressed in *Escherichia coli* DE3 from a pET-24a vector. Liter-scale cultures were grown to mid-log phase ($OD_{600}$=0.3-0.5) and induced with 0.4 mM isopropyl β-thiogalactoside (IPTG). Cultures were then grown for 16 hours at 18 °C and pelleted by centrifugation (20 min at 3000xg), and were stored at -20 °C for up to 2 months. Pellets were resuspended in Ni-NTA Loading Buffer (20 mM sodium phosphate, 500 mM sodium chloride, 20 mM imidazole, pH 7.4), lysed by either sonication (3 x 40 seconds), French press (two cycles, maximum pressure 1500 bar), or BPER-II protein lysis reagent (Thermo Fisher 78260), and cell debris was removed by centrifugation (60 min at 24,000 xG). Filtered supernatants were then loaded onto a Ni-NTA affinity column

(BioRad, Hercules, CA), washed in Ni-NTA Loading Buffer, and then eluted on a gradient to Ni-NTA Elution Buffer (20 mM sodium phosphate, 500 mM sodium chloride, 500 mM imidazole, pH 7.4). Fractions were analyzed on stain-free tris-glycine gels. Following purification, proteins were assessed by differential scanning fluorimetry using Sypro Orange dye to ensure that the constructs were folded.

**Expression and batch purification of ZG16B and ZG16P:**

The ZG16B and ZG16P constructs were transformed into BL21(DE3) cells and a 5 mL seed culture, supplemented with 30 µg/mL Kanamycin was grown in Luria-Bertani medium at 37°C for 18 hours at 225 rpm for each lectin. The overnight seed cultures were inoculated in 500 mL autoinduction media (0.1% (w/v) tryptone, 0.05% (w/v) yeast extract, 2 mM $MgSO_4$, 0.05% (v/v) glycerol, 0.005% (w/v) glucose, 0.02% (w/v) α-lactose, 2.5 mM $Na_2HPO_4$, 2.5 mM $KH_2PO_4$, 5 mM $NH_4Cl$, 0.5 mM $Na_2SO_4$) in a baffled flask, supplemented with 90 µg/mL kanamycin. The 500 mL cultures were incubated at 37°C for 4-5 hours at 225 rpm until bacterial growth reached log phase ($OD_{600}$ ~ 0.8-1). The incubation temperature was reduced to 18°C for the autoinduction of the protein expression for 18 hours at 225 rpm. The cells were harvested at 3000 rpm for 25 minutes at 4°C. The pellets were washed with 15 mL phosphate buffer saline and saved at -80°C.

The cell pellet (weight ~8.0 g for ZG16B and ZG16P) was resuspended in ~30 mL of lysis buffer (50 mM HEPES; pH 7.5, 300 mM NaCl, 20 mM imidazole, 10% glycerol, 5 mg lysozyme, 2 mg DNaseI, 2 mM $MgCl_2$) by rotating for 1 h at 4 °C. The cells were homogenized and lysed by sonicating twice for 90 s (1s ON and 2s OFF cycle) at 50% power output. The lysed cells were centrifuged at 30,000 RPM at 4°C for 60 min inTi45 rotor. The supernatant was collected, filtered through a 0.22 μm PES membrane filter and loaded to 3 mL Ni-NTA resin, pre-equilibrated with 10 CV buffer A (50 mM HEPES; pH 7.5, 300 mM NaCl, 5 mM $CaCl_2$, 20 mM Imidazole, 10% glycerol). The supernatant was batch-bound for 90 minutes at 4°C with gentle rotation. The unbound protein was collected by flowing through the protein-resin complex in a gravity column; the supernatant was flowed through thrice to ensure maximum binding of the lectin to the Ni-NTA beads. The 3 mL resin was washed with buffer A, containing 20, 50, 100, 200, and 400 mM of imidazole, 15 mL in each wash. The wash fractions were checked on 4-20% polyacrylamide gel and those containing ZG16B or ZG16P were pooled, and flash-frozen at -80 °C.

Fractions containing Ni-NTA purified ZG16B or ZG16P were thawed on ice. Meanwhile, the 3x 5 mL prepacked Hitrap desalting columns with Sephadex G-25 resin were equilibrated with 3CV of dialysis buffer (25 mM HEPES, pH 7.5, 150 mM NaCl, 10% glycerol), attaching to the AKTA purifier FPLC. The purified lectins are injected into the FPLC using a 5 mL loop and buffer exchanged at the flow rate of 1 mL/min, washing the columns with 4 CV of dialysis buffer. The desalted and buffer exchanged fractions were collected, checked on a 4-20% polyacrylamide gel, and stored at -80°C.

**Expression and purification of H-ficolin:**

The H-ficolin construct was transformed into BL21(DE3) and expressed in Terrific broth media in the same way as in ZG16B expression. The cells were harvested and saved at -80 $^0$C.

The cell pellet was resuspended in (4 mL/ g of pellet) lysis buffer (50 mM HEPES; pH 7.5, 300 mM NaCl, 5 mM $CaCl_2$, 20 mM Imidazole, 10% glycerol, 5 mg lysozyme, 2 mg DNaseI, 2mM $MgCl_2$ and 0.1% TritonX-100) by rotating for 1 h at 4 $^0$C. The cells were homogenized and lysed by sonicating twice for 90s (1s ON and 2s OFF cycle) at 50% power output. The lysed cells were centrifuged at 30,000 RPM at 4 $^0$C for 60 min inTi45 rotor. The supernatant was collected and filtered through a 0.22 µm PES membrane filter. The pellet was saved at -80 $^0$C for later use. The protein was incubated with 3mL Ni-NTA resin, pre-equilibrated with 15 mL of buffer A (50 mM HEPES; pH 7.5, 300 mM NaCl, 20 mM Imidazole, 10% glycerol) and batch bound for 90 min at 4 $^0$C with gentle rotation. The unbound protein was collected by flowing through the protein-resin complex in a gravity column. The 3 mL resin was washed with buffer A, containing 20, 50, 100, 200 and 400 mM of imidazole, 15 mL in each wash. The wash fractions were checked on 4-20% polyacrylamide gel and those containing H-ficolin were pooled and dialyzed overnight against 2L dialysis buffer (25 mM HEPES; pH 7.5, 150 mM NaCl, 5mM $CaCl_2$, 10% glycerol) at 4$^0$C.

The dialyzed fractions were further purified using a 5 mL pre-packed HiTrap Q HP ion-exchange chromatography column. The column was equilibrated with 3 CV of low salt buffer (50 mM HEPES; pH 7.5, 50 mM NaCl, 5 mM $CaCl_2$). The protein sample was injected through a 5 mL loop and washed with a 0-50% gradient of high salt buffer (50 mM HEPES; pH 7.5, 2 M NaCl, 5 mM $CaCl_2$) over 10 CV. The column was then washed with 5 CV 100% high salt buffer. The fractions containing H-ficolin were checked on a 4-20% polyacrylamide gel, pooled, concentrated and saved at -80$^0$C.

**Expression and purification of Strep-II tagged lectins from HEK293T:**

Strep-II tagged lectins were each expressed by transient transfection of suspension-adapted human embryonic kidney (HEK) 293T cells. Cells were transfected at 1.8 × $10^6$ cells/ml in growth medium [Dulbecco's modified Eagle's medium (Thermo Fisher Scientific, catalog no. 11995) supplemented with 10% heat-inactivated fetal bovine serum (FBS), penicillin-streptomycin (50 U/ml), 4 mM L-glutamine, and 1× nonessential amino acids] using Lipofectamine 2000 (Thermo Fisher Scientific) following the manufacturer's protocol. Six hours after the transfection, culture medium was exchanged to FreeStyle F17 expression medium (Thermo Fisher Scientific) supplemented with penicillin-streptomycin (50 U/ml), 4 mM L-glutamine, 1× nonessential amino acids, 0.1% heat-inactivated FBS, and 0.1% Pluronic F-68 (Thermo Fisher Scientific). Transiently transfected cells were cultured for up to 4 days or until viability was below 60%. The conditioned expression medium was then harvested by centrifugation and sterile filtration.

For purification of MBL, SPD, and CL-K1, $CaCl_2$ was added to the harvested conditioned expression medium to a final concentration of 10 mM, and avidin (7 mg/ml) was added at 12 μl per ml of expression media (IBA, catalog no. 2-0204-015; per the IBA protocol). Protein was captured onto 2 ml of Strep-Tactin Superflow High-Capacity resin (IBA Lifesciences, catalog no. 2-1208-002) equilibrated with Hepes-Ca buffer [20 mM Hepes (pH 7.4), 150 mM NaCl, and 10 mM $CaCl_2$]. The resin was then washed with Hepes-EDTA buffer [20 mM Hepes (pH 7.4), 150 mM NaCl, and 1 mM EDTA] and eluted with 5 mM *d*-desthiobiotin (Sigma-Aldrich) in Hepes-EDTA buffer. StrepII-tagged hItln1 was concentrated with a 30,000–molecular weight cutoff (MWCO) Amicon Ultra centrifugal filter. StrepII-tagged MBL was concentrated with a 10,000-MWCO Vivaspin 6 centrifugal filter (GE). All proteins were buffer exchanged to Hepes-EDTA buffer for storage. Protein concentrations were determined by absorbance at 280 nm.

For the purification of CRP and PTX3, a similar protocol was followed without the addition of calcium.

**Gravity flow-based sortase-mediated ligation:**

The gravity flow method of sortase-mediated ligation (SML) was developed based on the previously published syringe-pump flow method with minor modifications (1). Briefly, 40 µg of 6x histidine-tagged sortase A protein was incubated with 200 µL Ni-NTA resin, pre-equilibrated with sortase reaction buffer (50 mM HEPES, pH 7.5, 150 mM NaCl, 10 mM $CaCl_2$), on ice for 10 mins in 600 µL sortase reaction buffer. The sortase-bound resin was poured in a 2 mL regeneration gravity column and excess sortase and buffer were allowed to flow through. A 3 mL reaction mixture of lectin (final concentration 5 µM) and GGGYK-biotin- peptide/GGYK-SulfoCy5-peptide (21st Century Biochemicals) (final concentration 7 µM) was added to the column and the fractions were collected. The column was washed with an excess of 3 mL of the peptide (7 µM), followed by a 2 mL sortase buffer wash. Finally, the unreacted lectin was eluted from the column using a buffer containing imidazole (50 mM HEPES; pH 7.5, 300 mM NaCl, 400 mM Imidazole, 10% glycerol). The fractions with biotin-conjugated or fluorophore-conjugated lectin were dialyzed against 25 mM HEPES, pH 7.5, 150 mM NaCl, and 10% glycerol, using a 10 kDa MWCO snake-skin dialysis bag. The fractions were saved at -80 °C.

For SML of ZG16P and H-ficolin, 160 µg sortase A protein was used to increase the yield of the SulfoCy5-conjugated lectin.

**Dynamic light scattering (DLS) and Nano differential scanning fluorimetry (nDSF) of mGAPs:**

The cumulant radii of the recombinant lectins resuspended in 25 mM HEPES, pH 7.5 150 mM NaCl, 10% glycerol at 100 µg/mL were determined using a Prometheus NT.48 instrument.

The change in intrinsic tryptophan and tyrosine fluorescent intensity of recombinant lectins as a function of temperature was carried out in a Prometheus NT.48 instrument. Capillaries were filled with 10 µL recombinant lectins at 100 µg/mL concentration in 25 mM HEPES, pH 7.5 150 mM NaCl. The experiment was conducted using two replicates for each sample. The ratio of the emission intensities ($Em_{350nm}/Em_{330nm}$), representing the change in intrinsic fluorescence intensity, was recorded as the temperature was increased from 20 °C to 95 °C, taking one measurement per 0.0288°C. Both the raw fluorescence signal and the first derivative (slope) of the fluorescence signal as a function of the temperature were plotted. The melting temperature was calculated from the inflection point of the slope.

**Mucin binding assay**

Lyophilized MUC2, MUC5AC, and MUC5B were generously provided by Prof. Katharina Ribbeck, whose group purified them from porcine sources as previously described. These mucins were reconstituted to 10 µg/µL overnight at 4 °C in Mucin Assay Buffer (20 mM

HEPES, 150 mM sodium chloride, pH 7.4), then diluted to a concentration of 1 µg/µL in the same buffer. The porcine mucins Type II and type III (Sigma) were diluted at the same concentration and treated with 1mM sodium periodate at 37ºC overnight, protected from light.

*Mucin blot for H-ficolin and ZG16P GAPs:* 1 µg of the mucin samples were dotted on nitrocellulose membranes in triplicate. The membranes were air-dried and were blocked with 3% Bovine serum albumin (Sigma) in Tris buffer saline with 0.1% Tween-20 (TBST). The membranes were incubated with biotinylated GAPs at appropriate concentrations (500 nM, 250 nM and 100 nM) in the same buffer overnight at 4 °C or 2 h at room temperature. The blots were washed thrice with Tris buffer saline with 0.1% Tween-20 (TBST), incubated with alkaline-phosphatase conjugated Streptavidin at 1:10,000 dilution for 1 h, RT, and washed with TBST and TBS before the incubation with alkaline phosphatase substrate (Thermo Scientific).

*Mucin blot for Galectin GAPs:* 1 µg of the mucin solution was then spotted onto activated PVDF membranes and allowed to air dry at room temperature for 1 hour. The membranes were then blocked with 5% bovine serum albumin (BSA) and 0.1% Tween 20 in Mucin Assay Buffer overnight at 4 °C. The membranes were then incubated for 2 hours at room temperature with 250 nM of the galectin GAPs bearing a C-terminal Strep-tag in Mucin Assay Buffer with 0.1% BSA and 0.05% Tween 20. For each variant, one membrane was incubated with the above solution plus 50 mM lactose, while the other was incubated with a solution not containing lactose. After incubation and washing, the blots were incubated for 1 hour at room temperature with 1:5000 anti-His6-HRP antibody (Thermo Fisher Scientific MA121315HRP) and developed for 5 minutes with Clarity ECL reagent (BioRad). Binding was visualized using chemiluminescence on a ChemiDoc imager (BioRad).

The intensity of the chemiluminescence signal for each spot was quantified using the Fiji image suite. The signal from each spot was normalized to the average of the triplicate data from the highest-intensity sample.

**Screening GAPs on mammalian glycan array:**

A glass slide printed with mammalian glycans (Glycan Array 300 from RayBiotech) was taken out of -20$^{0}$C and equilibrated to room temperature inside the sealed plastic bag for 20-30 minutes. The slide was removed from the plastic bag; the cover film was peeled off and air-dried at room temperature for another 1-2 hours. The sub-arrays on the slide were blocked by adding 400µl binding buffer (20 mM HEPES; pH7.5, 150 mM NaCl, 10 mM $CaCl_2$, 0.1% BSA, 0.1% Tween 20) into each well and incubated at room temperature for 30 min. The buffer was decanted from the sub-arrays. Strep-II tagged lectins MBL, SPD and CL-K1, diluted to final concentration 10 µg/mL, 10 µg/mL and 50 µg/mL, respectively in 400 µL binding buffer, were added to three subarrays. The fourth array was treated as control by adding the same volume of buffer. The arrays were incubated with the lectins at 4°C overnight, shaking gently.

The next day, the lectins were decanted from the wells and the arrays were washed twice with 800 µL binding buffer and once with 800 µL wash buffer (20 mM HEPES; pH7.5, 150 mM NaCl, 10 mM $CaCl_2$), 5 minutes each at room temperature. The arrays were incubated with 4 μg/mL StrepMAB Dy549 or Streptavidin conjugated Cy3 in 400 μL binding buffer for 1 h at room temperature with gentle shaking. The sub-arrays were washed twice with 800 μL binding buffer and once with 800 μL wash buffer (20 mM HEPES; pH7.5, 150 mM NaCl, 10 mM $CaCl_2$), 5 minutes each at room temperature. The slide was disassembled from the chamber and washed in double-deionized water for 15 minutes in a slide holder tube. The slide was spin-dried and scanned at 532 nm laser using Genepix 4400A scanner.

The data obtained from the instrument was analyzed by using Microsoft Excel by subtracting the local background and blank control background and then averaging the fluorescence intensities of the three replicates for each glycan. The average intensity was plotted against the glycan ID number.

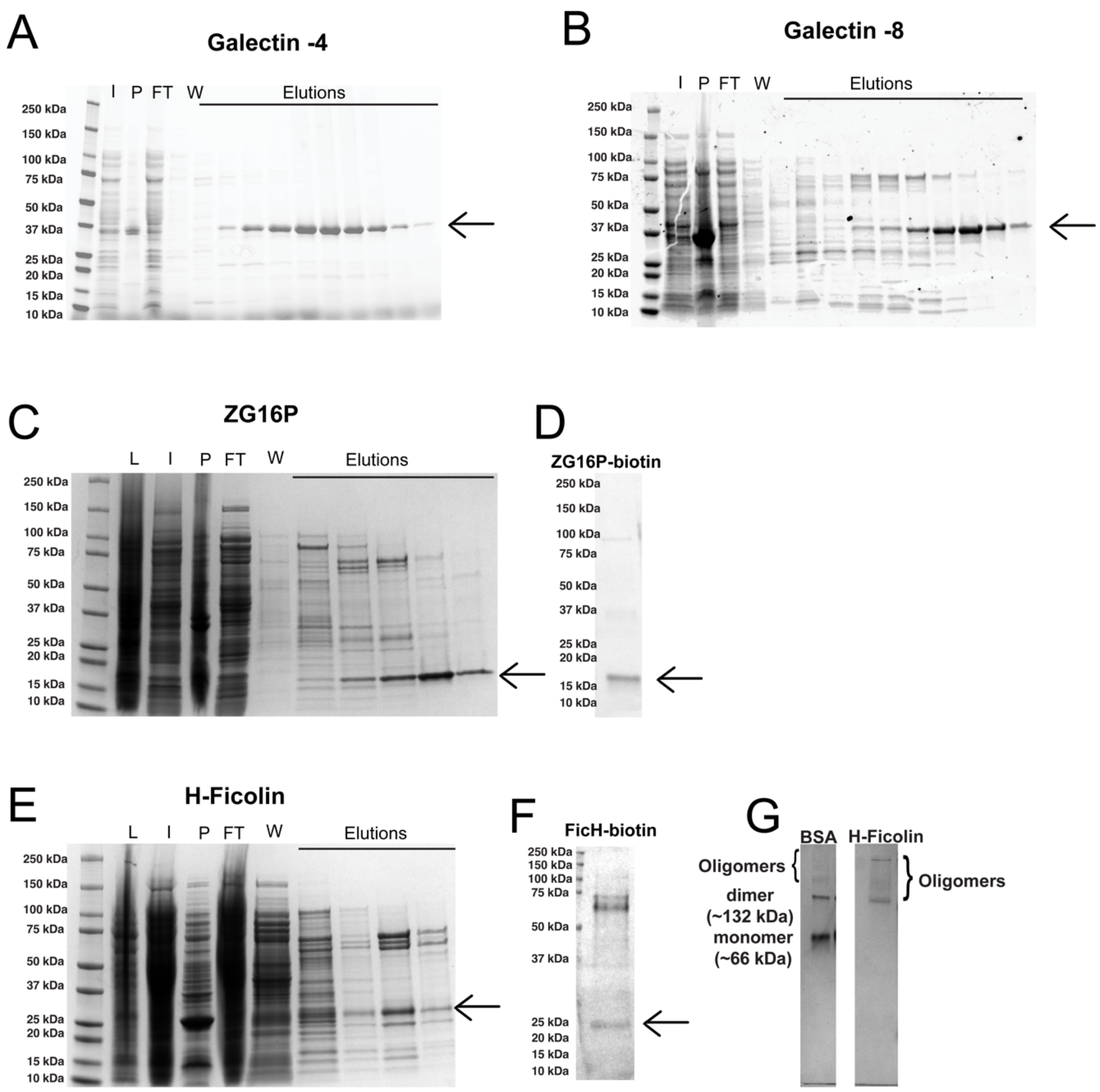


Figure S1: Recombinant production of human soluble lectins in E. coli BL21(DE3) strain. Gel electrophoresis of different fractions in affinity chromatography purification of (A) Strep-II-Galcetin-4-His$_6$, (B) Strep-II Galectin-8-His$_6$ (C) ZG16P-Linker-LPETG- His$_6$. The abbreviations used are: L-crude lysate, I-input or cleared lysate, P-pellet, FT-flow-through, and W-wash fractions. (D) Anti-streptavidin western blot of ZG16P conjugated to biotin produced by sortase-mediated ligation. (E) Gel electrophoresis of different fractions in affinity chromatography purification of neck and carbohydrate recognition domain of H-ficolin with a C-terminal LPETG motif and 6x Histidine tag. (F) Anti-streptavidin western blot of H-ficolin conjugated to biotin produced by sortase-mediated ligation. (G) Native gel electrophoresis of 6 $\mu$g of affinity-purified H-ficolin construct, showing the oligomeric formation in solution. Bovine-serum albumin (Sigma) resuspended in the lectin storage buffer (25 mM HEPES, pH 7.5, 150 mM NaCl, 5 mM CaCl2, 5% glycerol was loaded on the gel at the same amount for molecular weight standard.

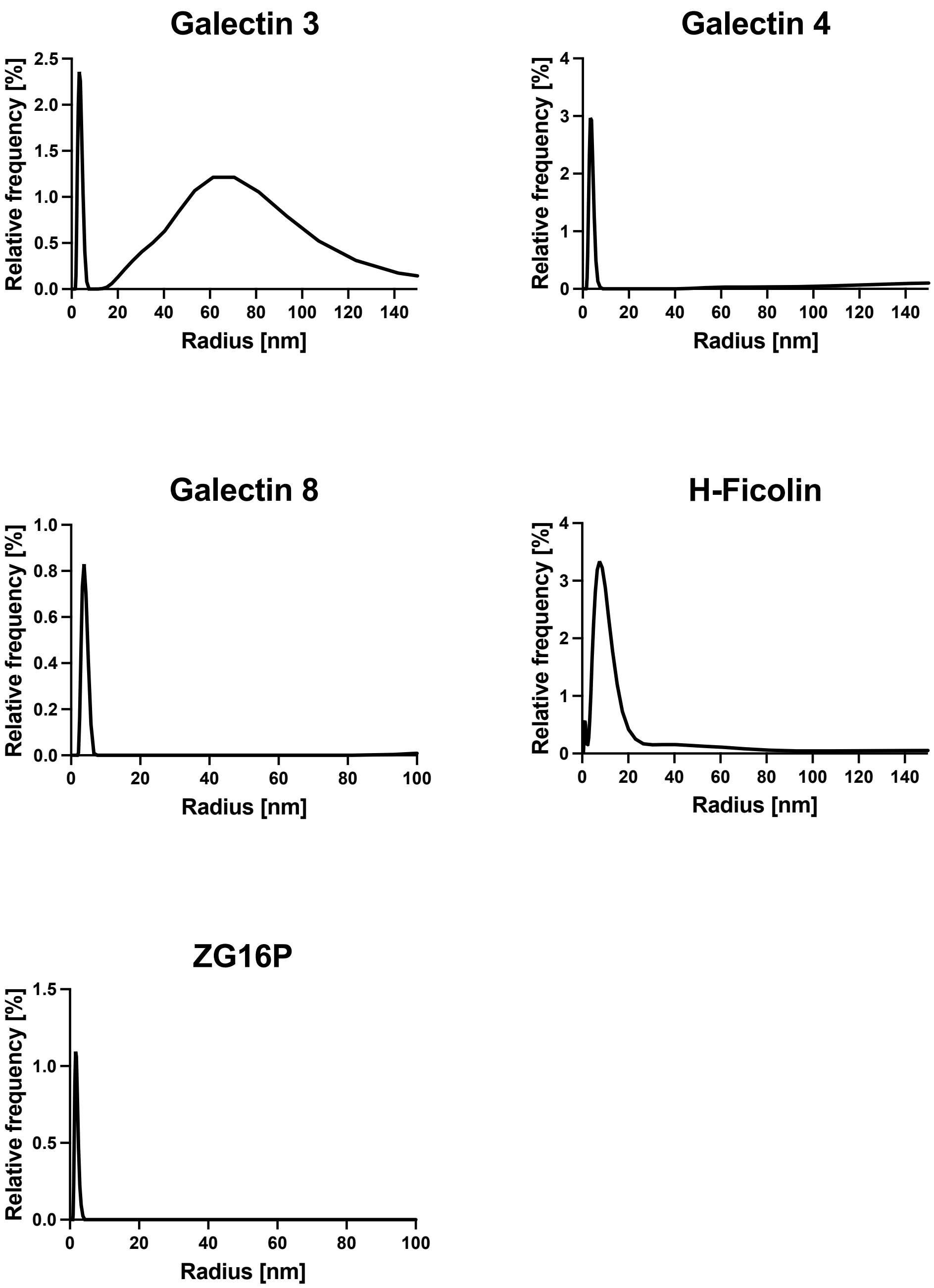


Figure S2: Intensity-weighted particle size distribution of GAPs produced from bacterial cells: Dynamic light scattering data showing the dispersity of recombinant lectins in solution. Traces represent the average of two independent experiments. The cumulant radii were calculated using the PR.Panta Control software.

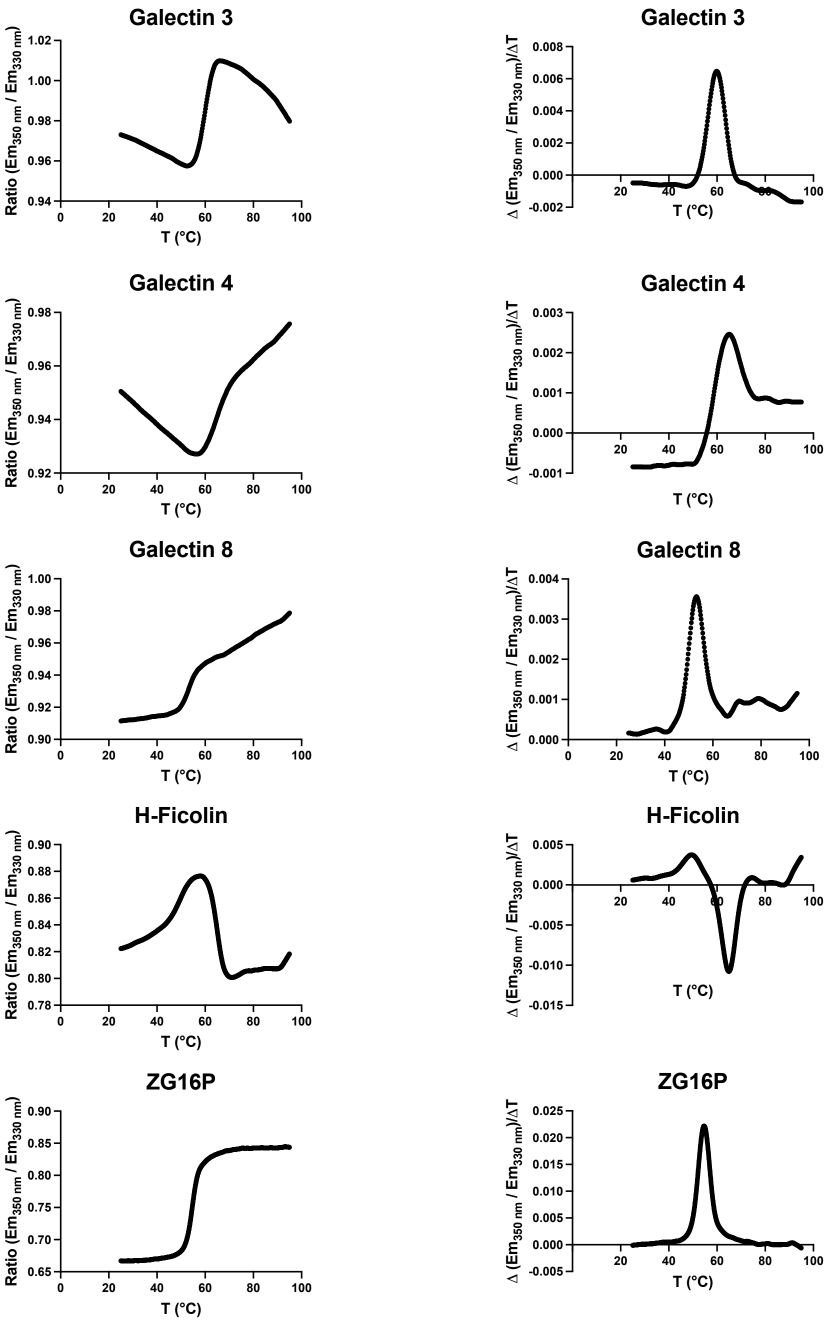


Figure S3: Thermostability of GAPs produced from bacterial cells: The ratio of change in intrinsic fluorescence intensity at 350 nm and 330 nm (left) and its first derivative (right) as a function of temperature for recombinant lectins obtained from nano-differential scanning fluorimetry. Traces represent the average of two independent experiments. The transition temperatures were calculated using the PR.Panta Control software.

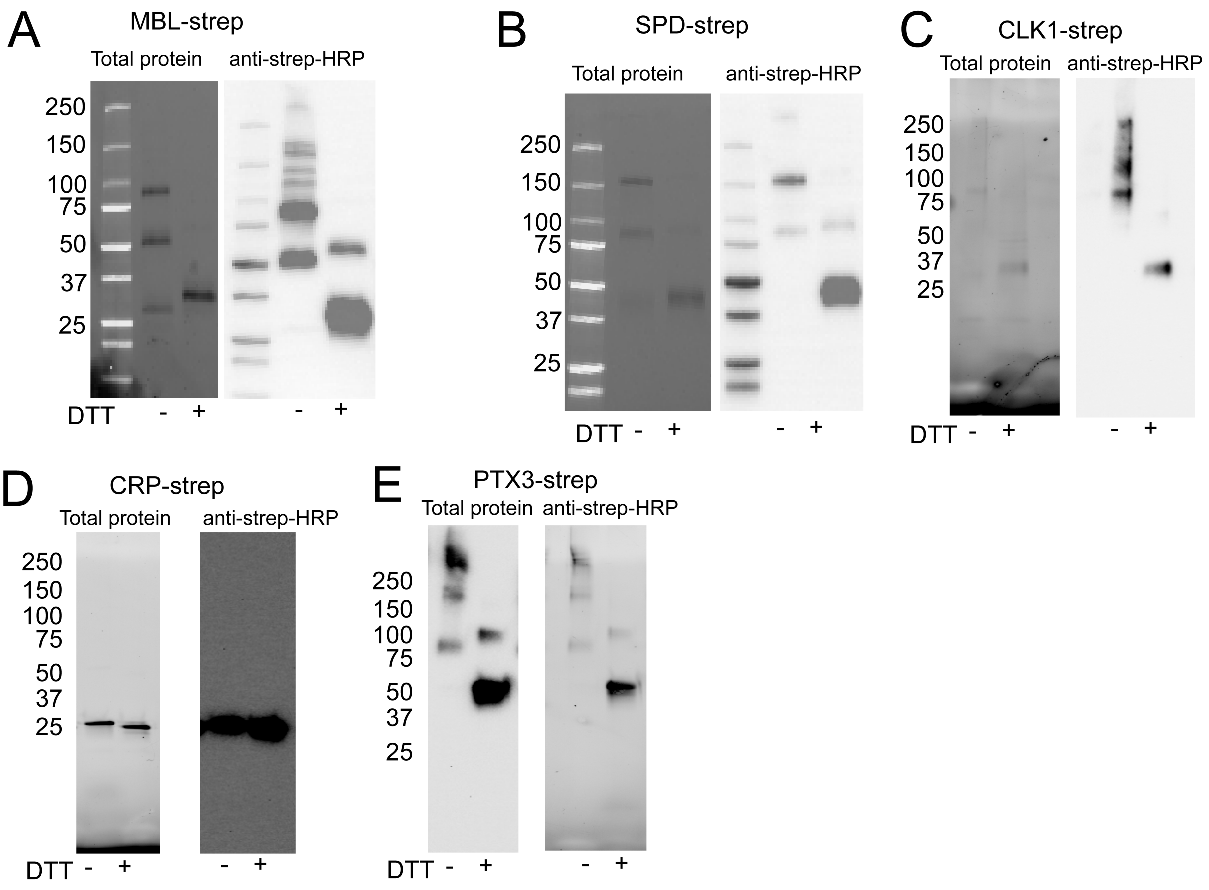


Figure S4: Recombinant production of human soluble lectins in Human embryonic kidney cells. Gel electrophoresis and anti-streptactin western blots of twin strep-tagged lectins (A) mannose-binding lectin 2 (MBL), (B) surfactant protein D (SPD), (C) Collectin -kidney-1, (D) C-reactive protein, and (E) Pentraxin 3 produced in HEK293T cells and purified by affinity chromatography. The purified lectins were treated with the reducing agent dithiothreitol (DTT) to monitor their oligomeric formation in the solution.

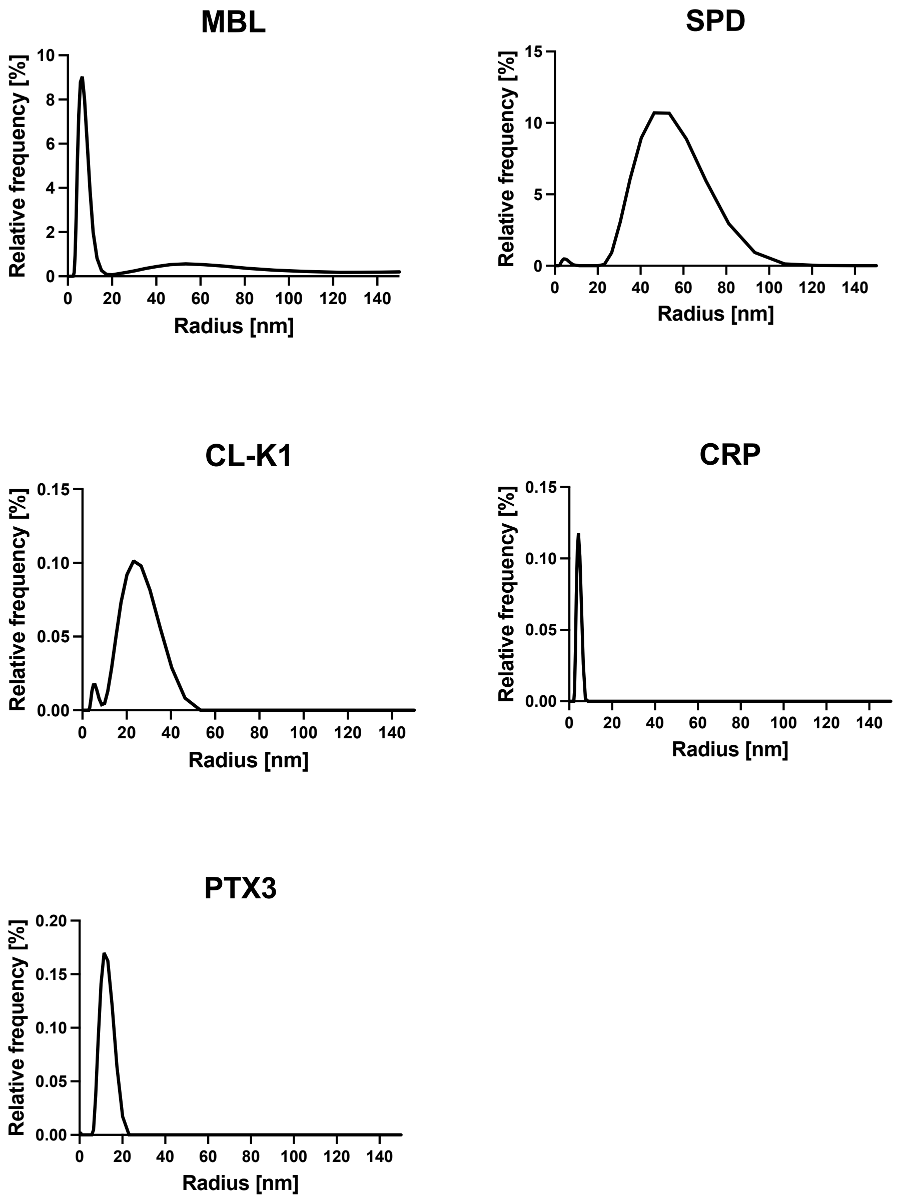


Figure S5: Intensity-weighted particle size distribution of GAPs produced from mammalian cells: Dynamic light scattering data showing the dispersity of recombinant lectins in solution. Traces represent the average of two independent experiments. The cumulant radii were calculated using the PR.Panta Control software.

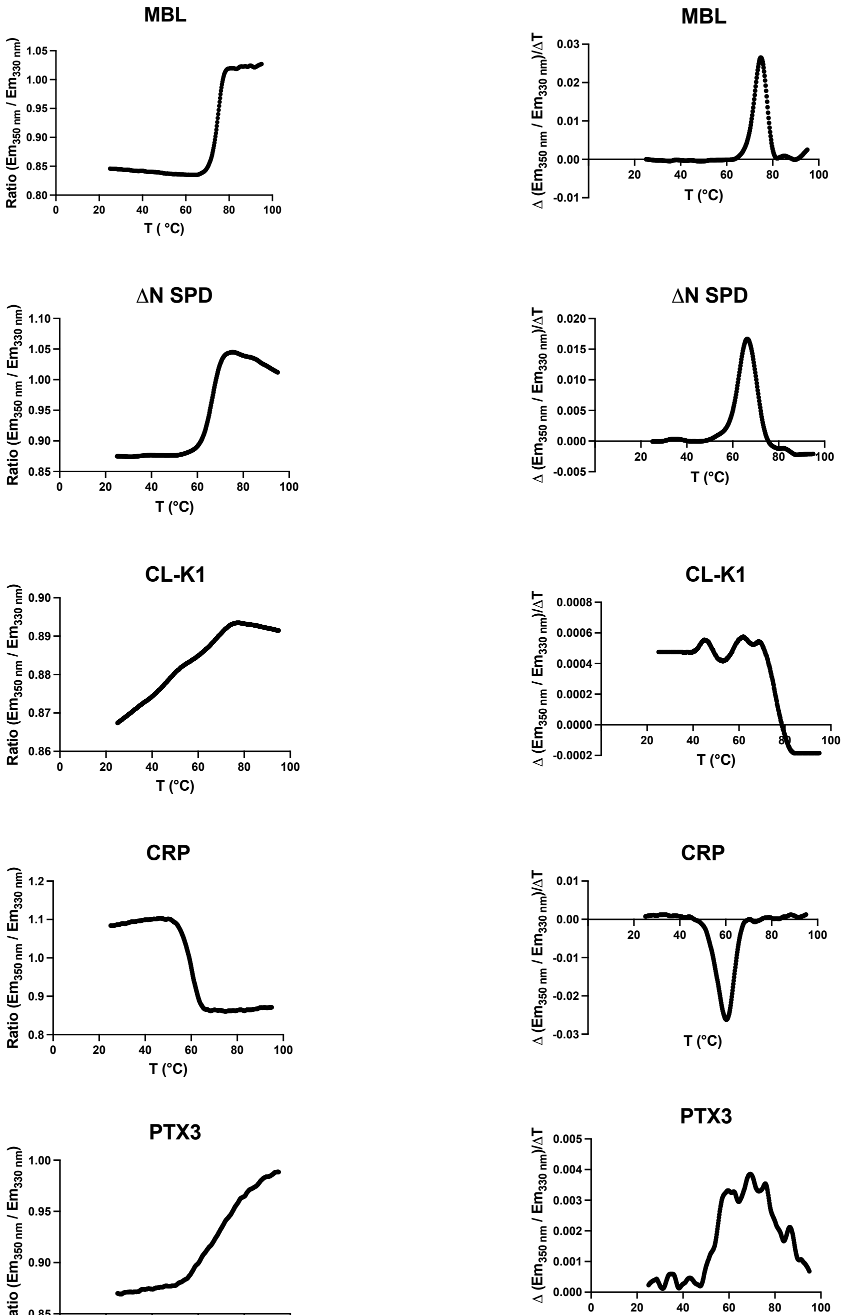


Figure S6: Thermostability of GAPs produced from mammalian cells: The ratio of change in intrinsic fluorescence intensity at 350 nm and 330 nm (left) and its first derivative (right) as a function of temperature for recombinant lectins obtained from nano-differential scanning fluorimetry. Traces represent the average of two independent experiments. The transition temperatures were calculated using the PR.Panta Control software.

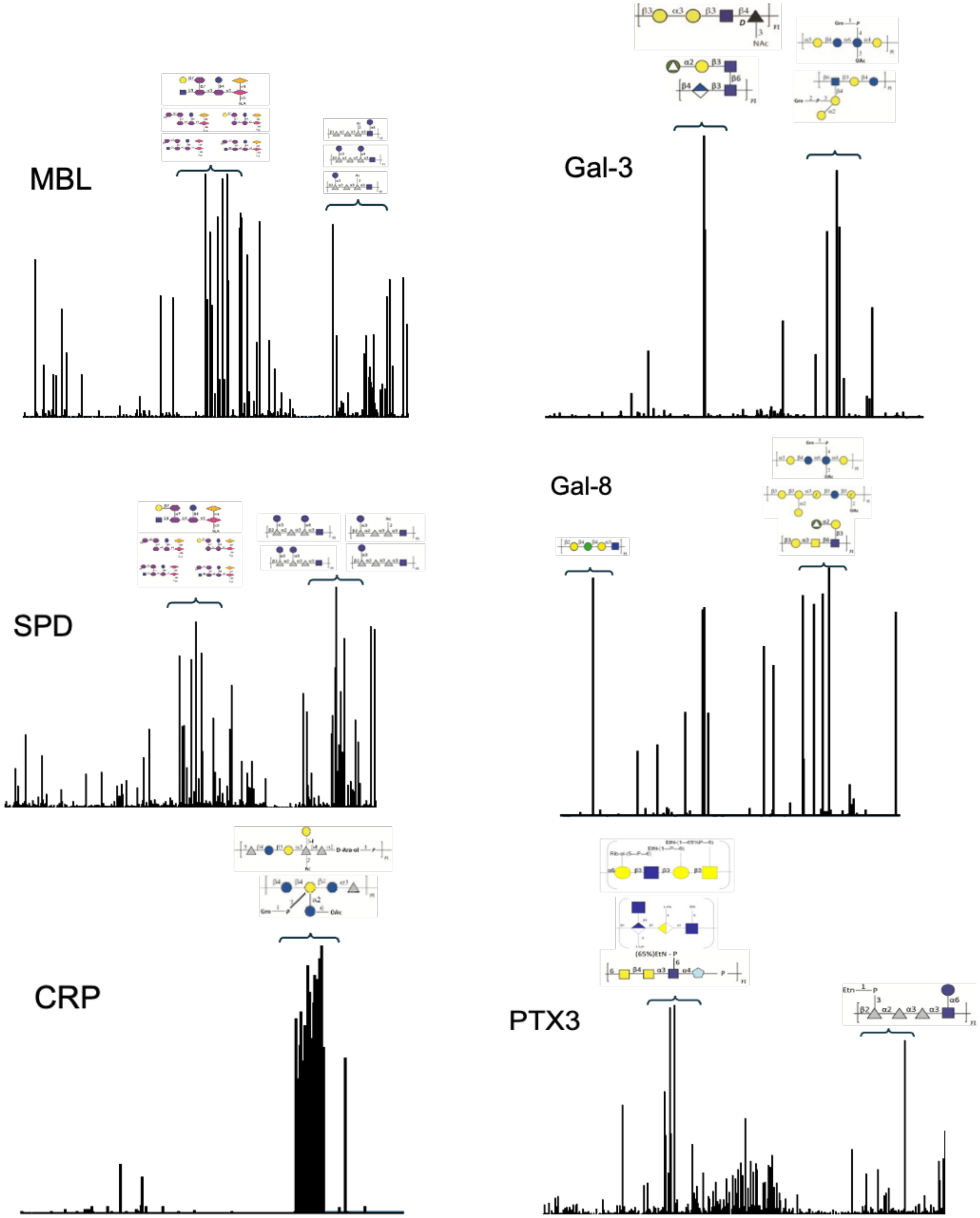
MBL
Gal-3
SPD
Gal-8
CRP
PTX3

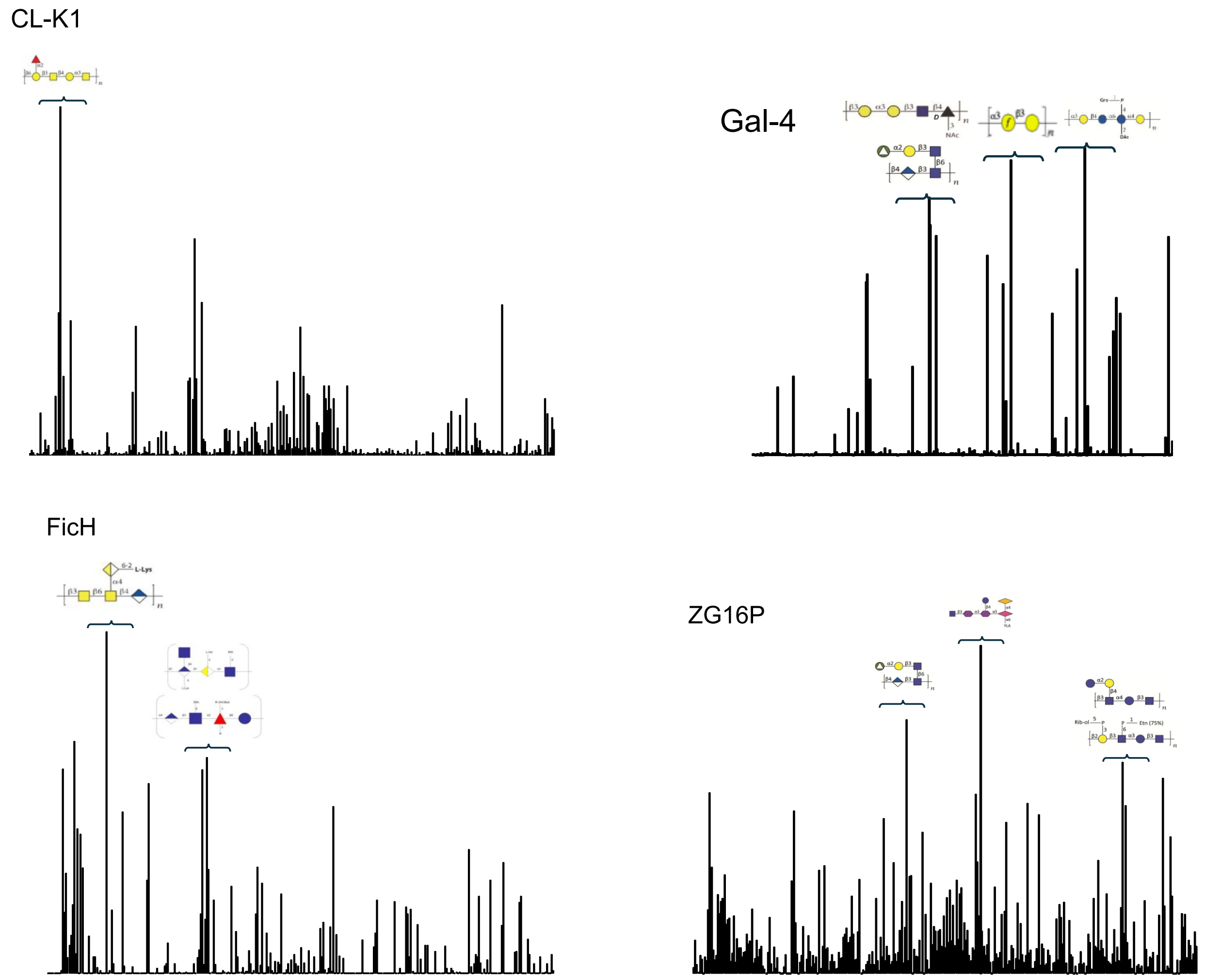


Figure S7: Glycan binding profile of GAPs assessed by microbial glycan array: GAPs (5-10 µg/mL) binding to microbial glycan microarray. The total binding intensity measured as relative fluorescence unit (RFU) is presented as the mean ± s.d. (*n*=6 technical replicates), as listed on supplementary information Spreadsheet 2. The structures of top glycan hits are shown by the data point. Please refer to Supplement 3 for the details of the glycan structures.

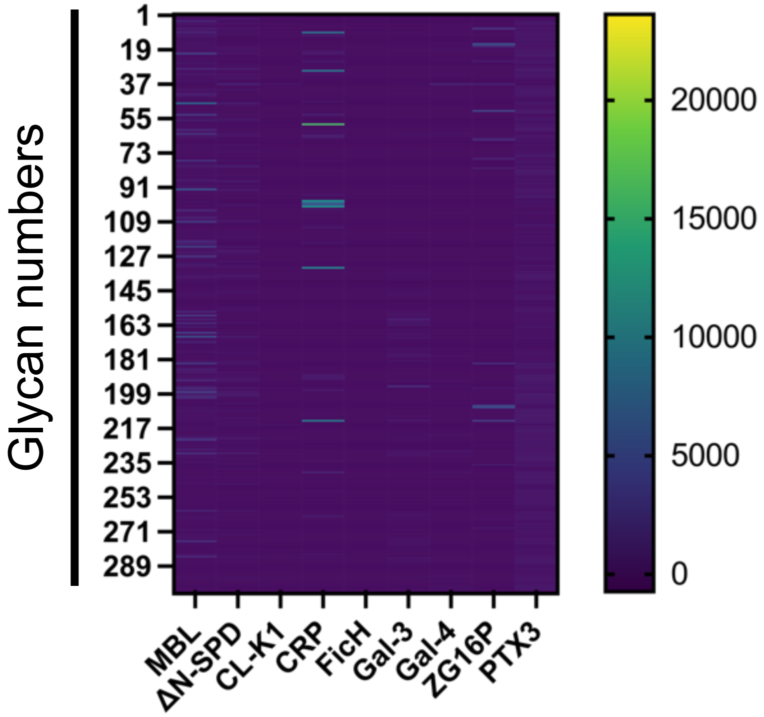


Figure S8: Glycan binding profile of GAPs on mammalian glycan array (RayBiotech) represented as heatmaps. Heatmap generated based on the total binding intensity quantified as relative fluorescence unit (on the scale). Data are presented as the mean ± s.d. (*n*=6 technical replicates).

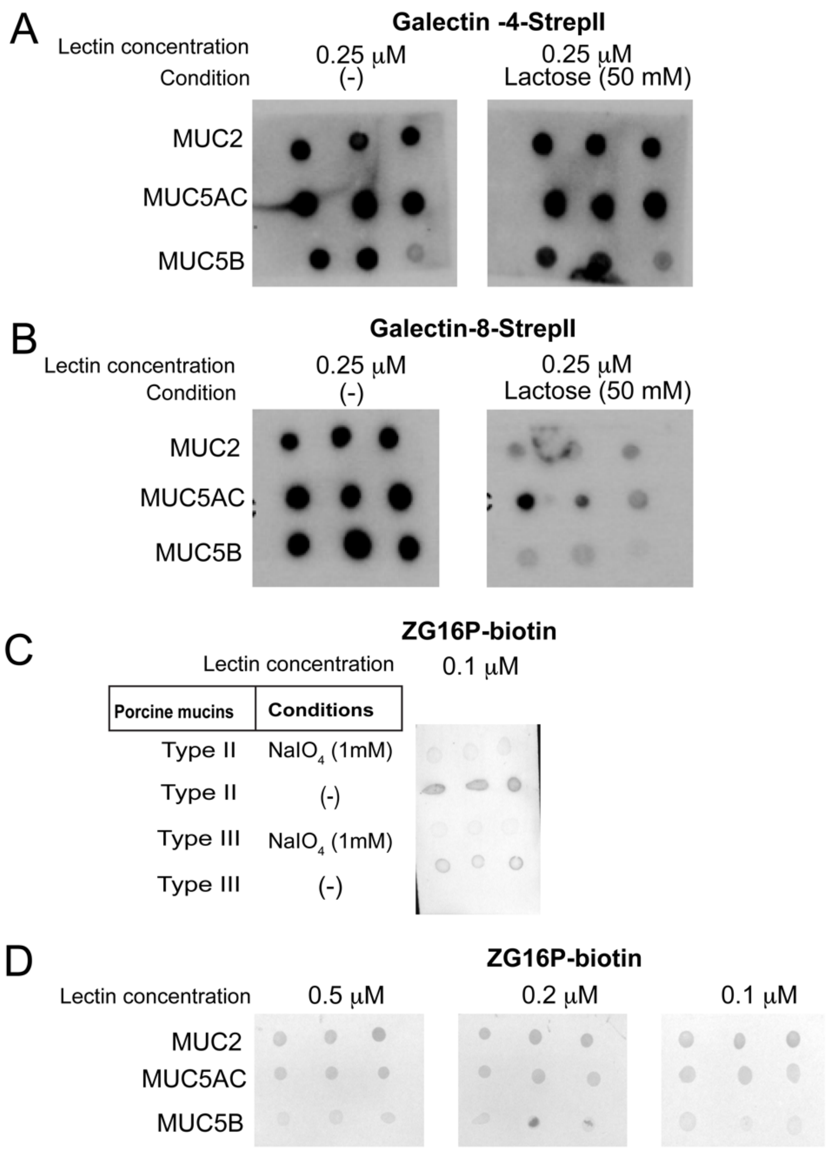


Figure S9: Mucin binding of GAPs: Porcine mucins MUC2, MUC5AC, and MUC5B, dotted on nitrocellulose membrane in triplicate, were probed with (A) The Strep-II tagged Galectin-4 GAP, (B) Strep-II Galectin-8 GAP at a 0.25 μM concentration. The blots were washed with or without 50 mM lactose in the mucin-binding buffer to test the glycan-mediated binding of GAPs to mucins. Porcine mucins MUC2, MUC5AC, and MUC5B, dotted on nitrocellulose membranes in triplicate, were probed with (C) The ZG16P Biotin GAP, at 0.1 μM, 0.25 μM, 0.5 μM concentrations, followed by incubating with alkaline phosphatase-conjugated streptavidin. (D) Crude Type II and Type III porcine mucins, treated with 1.0 mM sodium periodate to oxidize the vicinal diol present in mucin O-glycans, were dotted on nitrocellulose membranes in triplicates and probed with ZG16P-biotin GAP. The binding intensities were quantified using Fiji and represented as bar graphs, as shown in Figure 3 in the main text.

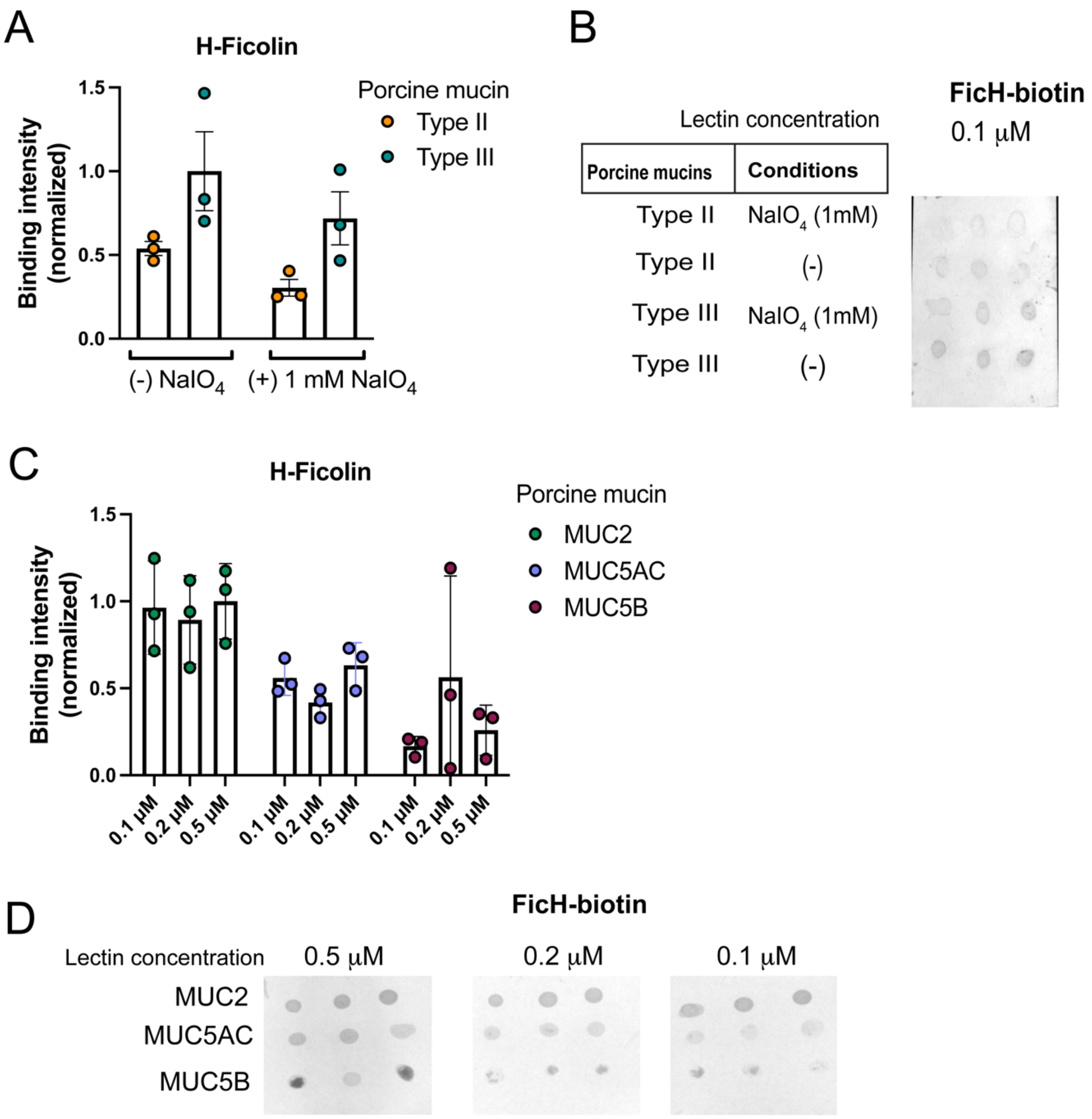


Figure S10: Mucin binding of H-Ficolin: (A) Binding of ZG16P-biotin GAPs at a 0.1 μM concentration to crude porcine mucins, treated with 1 mM sodium periodate to oxidize the vicinal-diols of mucin O-glycans. Data are represented as mean± SEM. (*n*=3 technical replicates). (B) Crude Type II and Type III porcine mucins, treated with 1.0 mM sodium periodate to oxidize the vicinal diol present in mucin O-glycans, were dotted on nitrocellulose membranes in triplicates and probed with ZG16P-biotin GAP. The binding intensities were quantified using Fiji and represented as bar graphs, as shown in Figure S6(A). (C) Binding of H-Ficolin-biotin GAP to purified porcine mucins MUC2, MUC5AC, and MUC5B at different concentrations. Data are represented as mean± SEM. (*n*=3 technical replicates). (D) Porcine mucins MUC2, MUC5AC, and MUC5B, dotted on nitrocellulose membranes in triplicate, were probed with H-Ficolin-biotin GAP, at 0.1 μM, 0.25 μM, 0.5 μM concentrations, followed by incubating with alkaline phosphatase-conjugated streptavidin. The binding intensities were quantified using Fiji and represented as bar graphs, as shown in Figure S6(C).

### SI-Spreadsheet 1

| | | | Lectins | | | | | | | | | |
|---|---|---|---|---|---|---|---|---|---|---|---|---|
| BPS# | BACTERIA / STRAIN | NAME / STRUCTURE /Cat.No. | MBL | ΔN SPD | CL-K1 | CRP | FicH | Gal-3 | Gal-4 | Gal-8 | PTX3 | ZG16P |
| 1 | Providencia stuartii O49 | PO49 Core-linked | 0.10 | 1.56 | 0.57 | 0.00 | -10.52 | 0.05 | 0.01 | 0.01 | 0.21 | 1.72 |
| 2 | Providencia stuartii O52 | PO52 Core-linked | 1.99 | 2.61 | 1.49 | 0.54 | -7.43 | 0.10 | 0.09 | 0.12 | 4.68 | 10.00 |
| 3 | Pseudomonas aerguinosa O4 (Habs serotype 4) | PO4 Core-linked | 0.20 | 0.83 | 0.64 | -0.01 | -11.75 | 0.01 | -0.15 | 0.00 | 0.79 | 2.74 |
| 4 | Pseudomonas aerguinosa O1 (Fisher immunotype 4) | PO1 Core-linked | 0.05 | 0.56 | -0.02 | 0.02 | -10.04 | 0.02 | 0.10 | 0.00 | 0.47 | 0.04 |
| 5 | Pseudomonas aerguinosa O2 (Fisher immunotype 3) | PO2 Core-linked | 0.17 | 1.04 | 0.57 | 0.09 | -5.73 | 0.65 | 0.06 | 0.09 | 5.51 | 5.56 |
| 6 | Pseudomonas aerguinosa O13 (Sandvik serotype II) | PO13 Core-linked | -0.01 | -0.05 | 0.09 | 0.01 | -4.32 | 0.08 | -0.08 | 0.26 | -0.21 | 4.35 |
| 7 | Pseudomonas aerguinosa O9 (9a, 9b, 9d) | PO9 Core-linked | 0.20 | 0.13 | 12.17 | -0.02 | -41.62 | 0.01 | 0.68 | -0.02 | 1.77 | -0.74 |
| 8 | Pseudomonas aerguinosa O6a (Habs serotype6, fraction IIa) | PO6a Core-linked-O-unit | 0.04 | 1.28 | 0.09 | 0.00 | -3.20 | 1.32 | -0.01 | 0.03 | 0.35 | 4.95 |
| 9 | Pseudomonas aerguinosa O6a (Habs serotype6, fraction IIb) | PO6a unsubstituted core | 0.07 | 2.33 | 0.71 | 0.00 | 0.78 | 0.05 | 0.02 | -0.02 | 0.79 | 3.82 |
| 12 | Salmonella typhimurium SL 11881 (Re mut) | LPS-L9516 | 65.06 | 17.89 | 4.25 | -0.01 | 59.83 | 0.07 | 0.57 | 0.01 | 7.66 | 19.46 |
| 13 | Salmonella typhimurium TV 119 (Ra mut) | LPS-L6016 | 2.35 | 2.96 | 1.55 | 0.03 | 17.79 | 0.13 | 0.59 | 0.04 | 3.99 | 55.09 |
| 14 | Salmonella typhimurium SL 684 (Rc mut) | LPS-L5891 | 1.66 | 9.66 | 2.72 | 0.00 | 29.31 | 0.04 | 0.73 | 0.05 | 8.60 | 32.43 |
| 15 | Pseudomonas aerguinosa O10 | L8643 | 0.10 | 1.45 | -0.03 | 0.02 | 4.65 | 1.58 | 0.03 | 0.12 | 0.51 | 7.01 |
| 16 | Salmonella typhimurium dodeca saccharide | 4809 | 0.03 | 0.97 | 0.32 | 0.01 | 8.20 | 0.19 | 0.09 | 0.01 | 0.77 | 3.99 |
| 17 | Salmonella enteritidis dodeca saccharide | 1262 | 0.08 | 1.02 | 0.69 | 0.00 | 11.01 | 0.09 | 0.06 | 0.01 | 0.52 | 5.36 |
| 18 | Salmonella typhimurium LPS | L2262 | 3.27 | 2.08 | 16.86 | 0.04 | 20.17 | 0.59 | 0.75 | 0.49 | 2.30 | 24.03 |
| 20 | Serratia marcescens LPS | L6136 | 21.28 | 2.97 | 1.36 | 0.01 | 68.03 | 0.48 | 0.30 | 0.04 | 2.28 | 18.26 |
| 22 | Escherichia coli K235 LPS | L2143 | 0.85 | 33.20 | 40.84 | 0.04 | 5.62 | 0.35 | 0.70 | 0.11 | 3.01 | 21.88 |
| 23 | Escherichia coli O128-B12 LPS | L2755 | 1.56 | 1.54 | 100.00 | 0.01 | 42.20 | 0.43 | 22.15 | 1.27 | 3.40 | 22.82 |
| 25 | Salmonella enterica abortus equi LPS | L5886 | 0.14 | 1.21 | 1.64 | 0.05 | -5.51 | 0.43 | 0.26 | 0.20 | 1.67 | 30.08 |
| 26 | Salmonella typhosa LPS | L2387 | 2.67 | 2.01 | 22.64 | 0.00 | 40.81 | 0.50 | 0.50 | 0.35 | 3.56 | 15.08 |
| 27 | Salmonella enteritidis LPS | L2012 | 2.29 | 0.65 | 2.10 | -0.02 | 30.80 | 0.38 | 0.12 | 0.07 | 2.06 | 16.44 |
| 28 | Shigella bodyii type2 | | 0.67 | 0.93 | 1.26 | 0.01 | -6.51 | 1.06 | 0.23 | 0.11 | 2.21 | 16.77 |
| 29 | Shigella bodyii type4 | | 3.15 | 3.12 | 4.37 | 0.35 | -3.24 | 0.05 | 0.05 | 0.02 | 4.82 | 3.63 |
| 30 | Shigella bodyii type10 | | 17.43 | 1.34 | 38.47 | -0.01 | -7.33 | 0.31 | 0.02 | 0.03 | 0.88 | 6.19 |
| 31 | Shigella dysennteriae type 3 | | 1.22 | 1.59 | 4.63 | 0.46 | 10.95 | 0.67 | -0.01 | 0.05 | 4.29 | 17.55 |
| 32 | Shigella dysennteriae type 8 (batch 12) | | 17.20 | 1.35 | 1.38 | 0.05 | -3.15 | 0.09 | -0.04 | 0.02 | 5.09 | 3.27 |
| 33 | Shigella dysennteriae type 11 | | 0.04 | 0.62 | -0.63 | 0.09 | -2.80 | 0.29 | 0.06 | 0.02 | 0.67 | 10.46 |
| 34 | Shigella dysennteriae type 13 | | 0.36 | 4.75 | 0.74 | 0.13 | 4.77 | 0.11 | -0.04 | 0.07 | 1.30 | 7.84 |
| 35 | Escherichia coli O29 | | 0.30 | 0.68 | 0.30 | 0.01 | 4.88 | 0.05 | 0.00 | 0.01 | 0.36 | 7.78 |
| 36 | Escherichia coli O40 | | 0.00 | 1.79 | 0.19 | 0.01 | 1.54 | 0.13 | 25.47 | 95.83 | 0.20 | 5.04 |
| 37 | Escherichia coli O106 | | 44.68 | 23.73 | 1.11 | 0.19 | -7.87 | 0.04 | -0.01 | 0.08 | 2.69 | 6.35 |
| 38 | Escherichia coli O130 | | 0.82 | 3.68 | 0.52 | 0.18 | 0.81 | 0.03 | 0.13 | 0.02 | 0.86 | 6.59 |
| 39 | Escherichia coli O148 | | 2.97 | 1.53 | 0.45 | 0.00 | -15.99 | 0.04 | 0.02 | 0.01 | 0.92 | 12.04 |
| 40 | Escherichia coli O150 | | 0.53 | 2.15 | 0.59 | 0.05 | -1.84 | 0.04 | 0.09 | 0.06 | 1.05 | 8.92 |
| 41 | Escherichia coli O180 | | 26.46 | 6.19 | 0.27 | 0.09 | -14.79 | 0.15 | 0.20 | 0.08 | 1.33 | 9.94 |
| 42 | Proteus mirabilis O3a, 3c (G1) | | -0.01 | 0.12 | -0.05 | 1.49 | 100.00 | 1.01 | 0.08 | 2.50 | 0.04 | 19.66 |
| 43 | Proteus mirabilis O8 (TG326) | | 0.20 | 0.75 | 0.60 | 0.00 | -1.18 | 0.04 | 0.18 | 0.03 | 0.73 | 3.12 |
| 44 | Proteus mirabilis O10 (HJ4320) | | 0.00 | -0.01 | 0.13 | 0.29 | 0.80 | 0.07 | -0.06 | 0.02 | -0.01 | 4.51 |
| 45 | Proteus mirabilis O29a, 29b (2002) | | 0.07 | 1.08 | 0.63 | 0.07 | 18.55 | 0.18 | 0.07 | 0.07 | 0.45 | 10.15 |
| 46 | Proteus mirabilis O50 (TG332) | | -0.02 | 0.01 | -0.01 | 0.05 | 2.66 | 0.05 | 0.09 | 0.00 | -0.26 | 4.89 |
| 47 | Proteus mirabilis O54a, 54b (10704) | | 0.23 | 1.25 | 1.40 | 0.08 | -8.24 | 0.02 | 0.29 | 0.22 | 2.47 | 7.40 |
| 48 | Proteus mirabilis O57 (TG319) | | -0.08 | -0.01 | -0.17 | 0.05 | -0.37 | 0.12 | -0.02 | 0.14 | -0.22 | 8.13 |
| 49 | Proteus penneri O8 (106) | | -0.04 | 0.45 | -2.84 | -0.02 | -8.14 | 0.08 | 0.06 | -0.01 | 0.19 | 2.21 |
| 50 | Proteus penneri O64a, 64b, 64d (39) | | 0.09 | 0.96 | 0.30 | 0.02 | -7.49 | 0.04 | 0.06 | 0.01 | 0.50 | 1.36 |
| 51 | Proteus penneri O66 (2) | | 1.39 | 0.62 | 0.86 | 0.00 | -13.74 | 0.14 | -0.01 | 0.03 | 0.25 | -0.41 |
| 52 | Proteus penneri O69 (25) | | 0.67 | 1.77 | 6.43 | 0.47 | 47.35 | 0.35 | 0.73 | 0.51 | 6.95 | 6.44 |
| 53 | Proteus penneri O71 (42) | | 17.48 | 8.37 | 2.21 | 0.06 | 2.10 | 2.04 | 0.87 | 0.33 | 2.22 | 17.00 |
| 54 | Proteus penneri O72a, 72b (4) | | 0.19 | 2.90 | 0.40 | 0.18 | -0.58 | 0.11 | 0.14 | 0.02 | 0.67 | 2.78 |
| 55 | Pseudomonas aerguinosa O2 (2a),2d,2f | IATS 10 , OPS | 0.31 | 0.39 | 0.05 | 0.10 | -14.68 | 0.07 | 0.11 | 0.24 | 0.67 | 0.02 |
| 56 | Pseudomonas aerguinosa O2 2a,2b | IATS 16 OPS | 0.03 | 1.64 | 0.57 | 0.33 | -2.80 | 0.18 | 0.27 | 0.05 | 1.97 | 8.12 |
| 57 | Pseudomonas aerguinosa O2 2a,2b,2e | IATS NO, OPS | 0.17 | 0.89 | 0.60 | 0.06 | -9.40 | 0.04 | 0.32 | 0.05 | 1.19 | 1.47 |
| 58 | Pseudomonas aerguinosa O2 2a,2d | IATS 5 OPS | 0.08 | 0.97 | -0.24 | 1.19 | 1.79 | 0.04 | 0.10 | 0.06 | 0.73 | 4.80 |
| 59 | Pseudomonas aerguinosa O2 Immuno 7 | IATS 18, OPS | 0.10 | 0.44 | 0.40 | 0.15 | -4.62 | 0.07 | 0.04 | 0.75 | 0.73 | 0.31 |
| 60 | Pseudomonas aerguinosa O3 3a,3b | IATS NO, OPS | -0.10 | 0.41 | 0.15 | 1.34 | -19.72 | 0.02 | 0.18 | 0.04 | 0.53 | -1.56 |
| 61 | Pseudomonas aerguinosa O3 3a,3b,3c | IATS 3, OPS | 0.14 | 0.29 | 0.29 | 0.01 | -26.91 | 0.10 | 0.14 | 0.10 | 0.50 | -0.34 |
| 62 | Pseudomonas aerguinosa O3 3a,3d | IATS NO, OPS | 0.00 | 0.91 | 0.51 | 1.33 | -10.88 | 0.06 | 0.15 | 0.06 | 1.16 | 2.00 |
| 63 | Pseudomonas aerguinosa O4 4a,4c | IATS NO, OPS | 0.34 | 2.86 | 0.92 | 0.04 | -2.56 | 0.15 | 0.23 | 0.03 | 4.03 | 5.61 |
| 64 | Pseudomonas aerguinosa O6 6a | IATS 6, OPS | 0.00 | 0.65 | 0.12 | 2.30 | -15.65 | 0.05 | -0.02 | 0.05 | 0.73 | -2.06 |

| | | | | | | | | | | | | |
|---|---|---|---|---|---|---|---|---|---|---|---|---|
| 65 | Pseudomonas aerguinosa O6 6a,6c | IATS NO, OPS | 0.24 | 0.72 | 2.71 | 0.54 | -14.34 | 0.21 | 0.08 | 0.21 | 4.31 | -7.64 |
| 66 | Pseudomonas aerguinosa O6 Immuno 1 | IATS NO, OPS | 0.09 | 0.94 | 1.38 | 0.09 | -15.85 | 0.06 | 0.18 | 0.10 | 2.06 | -3.85 |
| 67 | Pseudomonas aerguinosa O7 7a,7b,7c | IATS 7,LPS | 2.50 | 0.85 | 18.14 | 0.15 | 27.26 | 0.78 | 6.54 | 0.54 | 52.50 | 28.31 |
| 68 | Pseudomonas aerguinosa O7 7a,7b,7d | IATS 8,LPS | 0.18 | 0.40 | 2.97 | 0.04 | 55.67 | 2.29 | 1.03 | 0.13 | 3.44 | 49.59 |
| 69 | Pseudomonas aerguinosa O7 7a,7d | IATS NO, LPS | 0.27 | 0.59 | 36.99 | 0.04 | -32.30 | 0.66 | 0.33 | 0.28 | 4.00 | 15.44 |
| 71 | Pseudomonas aerguinosa O10 10a,10b | IATS 10, OPS | 0.17 | 0.53 | -1.99 | 0.20 | -23.19 | 0.09 | 0.13 | 0.04 | 0.88 | 5.86 |
| 72 | Pseudomonas aerguinosa O10 10a,10c | IATS 19, OPS | -0.12 | 0.29 | 0.60 | 0.10 | -18.52 | 0.01 | 0.16 | 0.04 | 0.88 | -5.99 |
| 73 | Pseudomonas aerguinosa O11 11a,11b | IATS 11, OPS | 0.14 | 0.58 | 0.33 | -0.01 | -0.62 | 0.05 | 0.07 | 0.03 | 0.87 | 3.41 |
| 74 | Pseudomonas aerguinosa O12 12 | IATS 12, OPS Habs 12 | 0.00 | 1.15 | 0.44 | 0.03 | 1.35 | 0.24 | 0.58 | 0.10 | 1.97 | 5.35 |
| 75 | Pseudomonas aerguinosa O13 13a,13c | IATS 14, OPS | 0.24 | 15.16 | 2.31 | 0.04 | -19.38 | 0.27 | 0.11 | 0.05 | 2.74 | 14.18 |
| 76 | Pseudomonas aerguinosa O14 14 | IATS 17,OPS Meitert X | 0.01 | 0.37 | 0.95 | 0.13 | -26.18 | 0.05 | 0.60 | 0.05 | 0.57 | 0.07 |
| 77 | Pseudomonas aerguinosa O15 15 | IATS 15, OPS | 0.16 | 0.65 | 0.68 | 0.01 | -8.90 | 0.02 | 0.27 | 0.08 | 1.91 | 4.30 |
| 78 | Proteus vulgaris O1 (18984)* | LPS | 0.11 | 0.78 | 3.79 | 2.49 | -5.76 | 8.61 | 14.92 | 25.99 | 3.28 | 11.56 |
| 79 | Proteus vulgaris O4 (PrK 9/57) | OPS | 0.06 | 0.67 | 0.14 | 0.00 | 2.10 | 0.04 | 0.02 | 0.01 | 0.71 | 6.65 |
| 80 | Proteus vulgaris O12 (PrK 25/57) | OPS | 0.03 | 0.96 | 0.37 | 0.14 | 2.80 | 0.11 | 0.06 | 0.02 | 1.08 | 1.86 |
| 81 | Proteus vulgaris O13 (8344) | OPS | 0.24 | 0.69 | 0.49 | 0.67 | 8.76 | 0.11 | 0.16 | 0.03 | 1.95 | 5.53 |
| 82 | Proteus vulgaris O15 (PrK 30/57) | OPS | 0.00 | 0.01 | -0.36 | 0.03 | -4.03 | 0.04 | -0.15 | 0.03 | 0.00 | 2.89 |
| 83 | Proteus vulgaris O17 (PrK 33/57) | OPS | 0.67 | 1.05 | 4.98 | 0.07 | -9.21 | 2.00 | 0.95 | 0.22 | 0.38 | 4.93 |
| 84 | Proteus vulgaris O19a (PrK 37/57) | OPS | 0.11 | 1.26 | 0.11 | 0.00 | -11.69 | 0.06 | 0.11 | 0.01 | 0.69 | -0.14 |
| 85 | Proteus vulgaris O21 (PrK 39/57)* | LPS | 4.50 | 1.13 | 6.79 | 0.26 | 2.74 | 1.73 | 13.67 | 3.07 | 13.71 | 31.46 |
| 86 | Proteus vulgaris O22 (PrK 40/57) | OPS | 0.14 | 1.08 | 0.19 | 0.00 | -10.73 | 0.08 | 0.06 | 0.01 | 1.07 | 1.46 |
| 88 | Proteus vulgaris O25 (PrK 48/57) | OPS | 0.28 | 0.54 | 0.26 | 0.04 | -13.59 | 0.14 | 0.03 | 0.02 | 1.92 | 8.34 |
| 89 | Proteus vulgaris O34 (4669)* | LPS | 1.23 | 16.08 | 6.64 | 18.21 | 0.31 | 1.44 | 1.26 | 0.52 | 15.17 | 32.88 |
| 90 | Proteus vulgaris O37a,b (PrK 63/57) | OPS | 0.23 | 2.12 | 1.09 | 0.16 | -3.08 | 0.12 | 0.16 | 0.03 | 2.61 | 7.22 |
| 91 | Proteus vulgaris O37a,c (PrK 72/57) | OPS | 1.19 | 1.99 | -0.70 | 0.03 | -2.26 | 0.09 | 0.10 | 0.00 | 1.75 | 7.73 |
| 92 | Proteus vulgaris O44 (PrK 67/57) | OPS | -0.18 | 0.58 | 0.01 | 0.07 | -4.43 | 0.07 | 56.29 | 0.02 | 0.68 | 0.82 |
| 93 | Proteus vulgaris O45 (4680) | OPS | 0.06 | 0.85 | 0.10 | 0.01 | 0.03 | 23.78 | 58.68 | 1.17 | 1.08 | 3.61 |
| 94 | Proteus vulgaris O53 (TG 276-10) | OPS | 0.68 | 0.85 | 2.63 | 0.42 | -2.53 | 0.26 | 0.35 | 0.05 | 3.77 | 5.73 |
| 95 | Proteus vulgaris O54a,54c (TG 103) | OPS | 0.15 | 0.81 | 1.24 | 0.06 | -8.19 | 0.14 | 24.49 | 0.44 | 2.91 | 3.69 |
| 96 | Proteus vulgaris O55 (TG 155) | OPS | 0.04 | 0.29 | 0.60 | 0.22 | -12.05 | 0.04 | 0.01 | 0.06 | 0.77 | 1.28 |
| 97 | Proteus vulgaris O65 (TG 251) | OPS | 0.02 | 0.88 | -1.11 | -0.04 | -12.39 | 3.28 | 0.99 | 28.51 | 2.99 | 1.45 |
| 98 | Proteus mirabilis O6 (PrK 14/57) | OPS | 0.18 | 1.67 | 0.56 | 0.00 | 2.74 | 0.10 | 0.00 | 0.02 | 1.26 | 6.46 |
| 99 | Proteus mirabilis O11 (PrK 24/57) | OPS | 1.34 | 0.72 | -1.23 | 0.02 | -1.72 | 0.03 | 0.33 | 0.00 | 1.62 | 4.57 |
| 100 | Proteus mirabilis O13 (PrK 26/57) | OPS | 3.28 | 2.74 | 0.87 | 0.04 | 4.59 | 0.10 | 0.10 | 0.03 | 2.33 | 4.84 |
| 101 | Proteus mirabilis O14a,14b (PrK 29/57) | OPS | 0.10 | 0.89 | -0.94 | 0.10 | 6.79 | 0.14 | 0.20 | 0.03 | 3.33 | 11.07 |
| 102 | Proteus mirabilis O16 (4652) | OPS | 8.45 | 10.08 | 21.33 | 1.25 | 17.42 | 0.63 | 0.52 | 0.71 | 58.57 | 11.97 |
| 103 | Proteus mirabilis O17 (PrK 32/57) | OPS | 1.50 | 5.16 | 22.18 | 1.07 | 59.53 | 0.55 | 0.91 | 1.09 | 37.86 | 13.90 |
| 104 | Proteus mirabilis O23a,b,d (PrK 42/57) | OPS | 0.11 | 1.23 | 0.27 | 0.03 | 2.50 | 0.16 | 0.06 | 0.03 | 1.35 | 9.29 |
| 106 | Proteus mirabilis O26 (PrK 49/57) | OPS | 2.83 | 5.33 | 15.88 | 0.66 | 15.69 | 0.54 | 0.29 | 0.50 | 20.01 | 11.01 |
| 107 | Proteus mirabilis O27 (PrK 50/57) | OPS | 2.81 | 10.62 | 62.15 | 2.70 | 63.10 | 2.64 | 2.28 | 3.19 | 98.82 | 23.66 |
| 108 | Proteus mirabilis O28 (PrK 51/57) | OPS | 0.98 | 5.92 | 21.95 | 13.56 | 30.40 | 0.73 | 0.37 | 2.60 | 38.66 | 11.94 |
| 109 | Proteus mirabilis O29a (PrK 52/57) | OPS | 0.10 | 0.51 | 0.71 | 0.02 | -0.37 | 0.10 | 0.13 | 0.06 | 1.21 | 4.13 |
| 110 | Proteus mirabilis O40 (10703) | OPS | 0.07 | 0.67 | 0.38 | 0.03 | 0.36 | 0.09 | 0.09 | 0.03 | 1.20 | 4.21 |
| 111 | Proteus mirabilis O41 (PrK 67/57) | OPS | 2.81 | 12.36 | 43.92 | 2.49 | 21.45 | 0.69 | 1.69 | 1.41 | 100.00 | 12.56 |
| 112 | Proteus mirabilis O51 (19011)* | LPS | 0.15 | 1.28 | 4.49 | 0.24 | 2.63 | 1.38 | 0.37 | 0.19 | 5.35 | 28.76 |
| 113 | Proteus mirabilis O74 (10705, OF) | OPS | 0.90 | 0.54 | 3.79 | 0.04 | -4.17 | 0.17 | 0.08 | 0.18 | 7.21 | -0.58 |
| 114 | Proteus mirabilis O75 (10702, OC) | OPS | 0.06 | 0.69 | 0.92 | 0.00 | -12.05 | 0.15 | 0.04 | 0.00 | 1.18 | 1.02 |
| 115 | Proteus mirabilis O77 (3 B-m) | OPS | -1.88 | 0.35 | -3.07 | 0.02 | -6.23 | 0.06 | 0.18 | 0.00 | 1.16 | 0.51 |
| 116 | Proteus penneri O31a (26) | OPS | 0.72 | 1.29 | 0.69 | 0.00 | -7.92 | 0.08 | 0.03 | 0.00 | 1.51 | 2.61 |
| 117 | Proteus penneri O52 (15) | OPS | -0.17 | 0.61 | -0.50 | 0.02 | 0.67 | 0.08 | -0.03 | 0.01 | 1.56 | 3.68 |
| 118 | Proteus penneri O58 (12) | OPS | 0.04 | 0.78 | 0.14 | 0.01 | 0.24 | 0.12 | 0.04 | 0.02 | 1.13 | 5.70 |
| 119 | Proteus penneri O59 (9) | OPS | 0.33 | 0.61 | 1.65 | 0.02 | -2.59 | 0.04 | 0.17 | 0.02 | 0.92 | 2.73 |
| 120 | Proteus penneri O61 (21) | OPS | 49.83 | 9.18 | 0.76 | 0.03 | 0.07 | 0.13 | 0.13 | 0.04 | 3.49 | 6.67 |
| 121 | Proteus penneri O62 (41) | OPS | 0.19 | 0.61 | 0.62 | 0.06 | -7.73 | 0.17 | 0.17 | 0.09 | 5.08 | 2.76 |
| 122 | Proteus penneri O63 (22) | OPS | 1.36 | 1.55 | 1.91 | 0.03 | 25.36 | 0.50 | 0.17 | 0.04 | 1.66 | 22.79 |
| 123 | Proteus penneri O64a,b,c (27) | OPS | 0.21 | 0.50 | 0.70 | 0.00 | -4.28 | 0.21 | 0.08 | 0.04 | 0.69 | 3.77 |
| 124 | Proteus penneri O65 (34) | OPS | 0.93 | 1.63 | 0.68 | 0.01 | -9.35 | 1.10 | 0.34 | 41.78 | 2.16 | 0.80 |
| 125 | Proteus penneri O67 (8) | OPS | 2.02 | 3.32 | 7.40 | 2.34 | 12.15 | 0.67 | 1.12 | 1.17 | 33.62 | 11.04 |
| 126 | Proteus penneri O68 (63) | OPS | 0.60 | 22.69 | 7.65 | 0.31 | 4.84 | 0.34 | 0.12 | 0.34 | 10.38 | 5.02 |
| 127 | Proteus penneri O70 (60) | OPS | 1.81 | 0.86 | 3.80 | 0.13 | 0.16 | 1.10 | 0.75 | 0.33 | 7.24 | 47.36 |
| 128 | Proteus penneri O73a,b (103) | OPS | 1.14 | 2.31 | 7.00 | 0.35 | 0.34 | 0.22 | 28.73 | 1.95 | 13.54 | 3.81 |

| | | | | | | | | | | | | |
|---|---|---|---|---|---|---|---|---|---|---|---|---|
| 129 | Proteus myxofaciens O60 | OPS | 0.28 | 0.87 | 0.15 | 0.08 | 1.60 | 0.50 | 0.54 | 0.14 | 5.39 | 8.20 |
| 130 | Proteus O56 (genomospecies 4) | OPS | 49.22 | 35.47 | 1.12 | 0.02 | -0.36 | 0.08 | 0.05 | 0.02 | 2.90 | 5.45 |
| 131 | Providencia stuartii O4 | OPS | 1.12 | 0.76 | -2.69 | 0.01 | 0.43 | 0.43 | 1.16 | 0.04 | 1.97 | 2.70 |
| 132 | Providencia stuartii O18 | OPS | 0.30 | 3.03 | 0.28 | 0.00 | 0.46 | 0.21 | 0.03 | 0.03 | 0.63 | 3.52 |
| 133 | Providencia stuartii O20* | LPS | 1.80 | 1.92 | 6.94 | 0.23 | 6.56 | 0.53 | 0.91 | 0.31 | 19.96 | 33.89 |
| 134 | Providencia stuartii O43 | OPS | 0.32 | 1.63 | 2.23 | 0.01 | 2.77 | 0.18 | 0.07 | 0.01 | 3.21 | 5.93 |
| 135 | Providencia stuartii O44 | OPS | 1.31 | 3.74 | 0.92 | 0.02 | -2.15 | 0.09 | 0.04 | 0.08 | 1.95 | 3.55 |
| 136 | Providencia stuartii O47 | OPS | 0.08 | 2.00 | 0.68 | 0.03 | 0.81 | 0.10 | 0.01 | 0.04 | 1.15 | 6.35 |
| 137 | Providencia stuartii O47, Core 9 | OPS | 0.28 | 1.52 | 3.91 | 0.20 | 17.33 | 0.26 | 0.24 | 0.10 | 8.59 | 13.21 |
| 138 | Providencia stuartii O49, Core 1 | OPS | 0.42 | 1.58 | 4.77 | 0.45 | 31.17 | 0.40 | 0.50 | 0.11 | 7.57 | 21.93 |
| 139 | Providencia stuartii O57 | OPS | 0.16 | 0.77 | 0.83 | 0.03 | -0.78 | 0.05 | 0.15 | 0.03 | 1.40 | 3.04 |
| 140 | Providencia alcalifaciens O5 | OPS | 0.20 | 0.81 | 0.93 | 0.07 | 2.22 | 100.00 | 83.57 | 83.20 | 3.37 | 13.97 |
| 141 | Providencia alcalifaciens O6* | LPS | 0.25 | 1.14 | 8.04 | 0.12 | 26.33 | 66.68 | 74.72 | 84.15 | 11.14 | 77.48 |
| 142 | Providencia alcalifaciens O19 | OPS | 0.03 | 0.80 | -1.27 | 0.02 | -5.52 | 0.11 | 0.33 | 0.05 | 3.72 | 3.94 |
| 143 | Providencia alcalifaciens O19 | LPS | 0.30 | 1.92 | 9.38 | 0.15 | -4.90 | 0.55 | 0.58 | 0.41 | 15.53 | 29.71 |
| 144 | Providencia alcalifaciens O19 | LPS/NaOH | 0.14 | 2.74 | 6.69 | 0.17 | 11.66 | 1.30 | 1.17 | 0.15 | 6.12 | 29.79 |
| 145 | Providencia alcalifaciens O21 | OPS | 0.46 | 1.09 | 3.50 | 0.15 | 1.76 | 0.57 | 71.25 | 41.43 | 4.83 | -8.24 |
| 146 | Providencia alcalifaciens O23 | OPS | 0.15 | 1.01 | 2.71 | 0.13 | 5.82 | 0.76 | 1.78 | 0.02 | 7.03 | 7.33 |
| 147 | Providencia alcalifaciens O27 | OPS | -0.52 | 0.98 | 1.57 | 0.04 | 0.75 | 0.09 | 0.67 | 0.02 | 2.33 | 5.19 |
| 148 | Providencia alcalifaciens O29 | OPS | 0.13 | 4.28 | 0.80 | 0.03 | -7.37 | 0.10 | 0.08 | 0.00 | 3.18 | 4.90 |
| 149 | Providencia alcalifaciens O30 | OPS | 2.20 | 1.80 | 4.06 | 0.16 | 3.48 | 0.20 | 0.53 | 0.10 | 6.99 | 10.00 |
| 150 | Providencia alcalifaciens O32 | OPS | 0.06 | 3.19 | 0.04 | 0.05 | -2.84 | 0.14 | 0.16 | 0.00 | 3.80 | 0.20 |
| 151 | Providencia alcalifaciens O36* | LPS-NH4OH | 0.60 | 1.31 | 8.09 | 1.03 | 3.18 | 0.62 | 0.71 | 0.52 | 17.05 | 43.03 |
| 152 | Providencia alcalifaciens O39 | OPS | 0.16 | 0.93 | 1.42 | 0.07 | 2.56 | 0.08 | 0.12 | 0.00 | 5.33 | -0.52 |
| 153 | Providencia rustigianii O14 | OPS | 0.49 | 1.85 | 9.23 | 0.18 | 23.12 | 0.97 | 0.32 | 0.06 | 10.68 | 17.91 |
| 154 | Providencia rustigianii O16 | OPS | 0.06 | 0.17 | 2.21 | 0.07 | -8.90 | 0.25 | 0.16 | 0.03 | -0.32 | 4.13 |
| 155 | Providencia rustigianii O34 | OPS | 0.72 | 1.10 | 0.28 | 0.02 | -4.32 | 0.60 | 0.16 | 0.01 | 2.64 | 1.26 |
| 156 | Yersinia pestis, KM260(11)-Δ0187 | LPS | 99.96 | 68.80 | 21.30 | 0.21 | 1.04 | 0.58 | 0.57 | 0.12 | 18.64 | 27.58 |
| 157 | Yersinia pestis, KM260(11)-Δ0187 | Core oligo saccharide | 0.47 | 2.31 | 1.46 | 0.01 | 2.69 | 0.08 | 0.05 | 0.00 | 4.05 | 8.61 |
| 158 | Yersinia pestis, KM260(11)-Δrfe | LPS | 48.53 | 36.74 | 12.62 | 0.14 | -17.02 | 0.55 | 0.64 | 0.11 | 17.20 | 14.23 |
| 159 | Yersinia pestis, KM260(11)-Δrfe | Core oligo saccharide | 0.50 | 2.63 | 2.66 | 0.00 | 3.79 | 0.09 | 0.15 | 0.00 | 10.07 | 12.86 |
| 160 | Yersinia pestis, 1146-25 | LPS | 76.42 | 37.25 | 14.07 | 0.17 | -14.86 | 0.32 | 0.71 | 0.17 | 21.59 | 6.95 |
| 161 | Yersinia pestis 1146-25 | Core oligo saccharide | -0.12 | 1.76 | 1.41 | 0.02 | 5.55 | 0.17 | 0.00 | 0.02 | 3.07 | 12.97 |
| 162 | Yersinia pestis, 1146-37 | LPS | 46.09 | 18.02 | 11.67 | 0.17 | 2.72 | 0.53 | 1.77 | 0.22 | 24.63 | 13.46 |
| 163 | Yersinia pestis, 1146-37 | Core oligo saccharide | -0.26 | 1.48 | 3.28 | 0.03 | 2.15 | 0.13 | 0.01 | 0.01 | 11.03 | 8.15 |
| 164 | Yersinia pestis, 0KM218-37 | LPS | 9.81 | 4.32 | 5.08 | 0.10 | -0.90 | 1.46 | 0.63 | 0.08 | 18.46 | 17.55 |
| 165 | Yersinia pestis, KM218-37 | Core oligo saccharide | -0.11 | 1.62 | 1.91 | 0.00 | 7.45 | 0.09 | -0.04 | 0.01 | 3.63 | 7.78 |
| 166 | Yersinia pestis, KM218-25 | LPS | 82.58 | 67.27 | 23.85 | 0.18 | -4.16 | 0.34 | 0.99 | 0.20 | 46.06 | 23.03 |
| 167 | Yersinia pestis, KM218-25 | Core oligo saccharide | -0.23 | 1.63 | 2.19 | 0.01 | 3.05 | 0.12 | 0.02 | 0.01 | 1.25 | 9.43 |
| 168 | Yersinia pestis, KM260(11)-ΔpmrF | LPS | 15.40 | 7.33 | 4.80 | 0.07 | -15.91 | 1.02 | 0.40 | 0.03 | 12.47 | 25.35 |
| 169 | Yersinia pestis, KM260(11)-ΔpmrF | Core oligo saccharide | 0.09 | 2.52 | 1.80 | 0.01 | -1.32 | 0.17 | 0.00 | 0.01 | 3.37 | 9.87 |
| 170 | Yersinia pestis, KM260(11)-Δ0186 | LPS | 98.11 | 84.27 | 36.84 | 0.07 | 6.59 | 0.62 | 2.14 | 0.15 | 25.66 | 20.84 |
| 171 | Yersinia pestis, KM260(11)-Δ0186 | Core oligo saccharide | 0.12 | 2.44 | 1.46 | 0.00 | 0.16 | 0.15 | 0.04 | 0.00 | 4.73 | 8.51 |
| 172 | Yersinia pestis, KM260(11)-ΔwaaQ | LPS | 15.43 | 13.20 | 22.51 | 0.11 | -18.24 | 2.83 | 1.13 | 0.10 | 20.09 | 28.37 |
| 173 | Yersinia pestis, KM260(11)-ΔwaaQ | Core oligo saccharide | 0.08 | 2.36 | 2.03 | 0.03 | 4.83 | 0.19 | 0.06 | 0.00 | 4.28 | 10.37 |
| 174 | Yersinia pestis, KM260(11)-ΔwaaL | LPS | 100.00 | 70.11 | 17.67 | 0.09 | -11.04 | 0.78 | 1.57 | 0.08 | 24.41 | 24.19 |
| 175 | Yersinia pestis, KM260(11)-25 | LPS | 56.11 | 25.85 | 17.16 | 0.11 | -11.81 | 0.78 | 0.42 | 0.17 | 21.47 | 26.15 |
| 176 | Yersinia pestis, KM260(11)-25 | Core oligo saccharide | 0.43 | 1.46 | 0.59 | 0.01 | 0.47 | 0.11 | 0.13 | 0.01 | 4.40 | 6.85 |
| 177 | Yersinia pestis, KM260(11)-37 | Core oligo saccharide | -0.17 | 1.78 | 2.03 | 0.03 | 13.13 | 0.46 | 0.10 | 0.01 | 4.26 | 15.52 |
| 178 | Yersinia pestis, KIMD1-37 | Core oligo saccharide | 0.56 | 3.14 | 1.42 | 0.00 | 4.88 | 0.40 | -0.12 | 0.00 | 2.09 | 10.14 |
| 179 | Yersinia pestis, KIMD1-25 | Core oligo saccharide | 3.06 | 1.76 | 0.96 | 0.02 | 14.92 | 0.61 | 0.20 | 0.03 | 1.09 | 13.22 |
| 180 | Yersinia pestis, 11M-25 | LPS | 0.68 | 1.76 | 9.45 | 0.10 | 1.01 | 0.25 | -0.20 | 0.13 | 15.01 | 5.53 |
| 181 | Yersinia pestis, 11M-37 | LPS | 0.33 | 2.69 | 8.35 | 0.45 | -2.86 | 0.18 | 1.21 | 0.19 | 15.56 | 3.75 |
| 182 | Proteus mirabilis O23a, 23b, 23c (CCUG 10701) | OPS | 0.10 | 1.89 | 1.54 | 0.03 | 1.51 | 0.15 | 0.09 | 0.00 | 2.87 | 7.68 |
| 183 | Proteus vulgaris O24 (PrK 47/57) | LPSOH | -0.14 | 1.29 | 6.49 | 0.21 | 13.54 | 0.72 | 64.90 | 2.90 | 7.53 | 7.12 |
| 184 | Yersinia pestis KM260(11)-6C | LPS | 77.68 | 40.80 | 19.78 | 0.09 | 0.62 | 2.53 | 0.54 | 0.19 | 23.64 | 54.68 |
| 185 | Yersinia pestis 260(11)-37C-186 | LPS | 84.03 | 12.12 | 16.20 | 0.12 | 48.82 | 1.48 | 0.59 | 0.12 | 22.63 | 34.09 |
| 186 | Yersinia pestis 260(11)-37C-187 | LPS | 82.28 | 16.02 | 12.76 | 0.06 | -9.18 | 0.88 | 1.16 | 0.08 | 13.95 | 17.62 |
| 187 | Yersinia pestis 260(11)-37C-416 | LPS | 1.35 | 2.89 | 19.92 | 0.17 | 11.25 | 2.86 | 0.36 | 0.18 | 28.41 | 100.00 |
| 188 | Yersinia pestis 260(11)-37C-417 | LPS | 5.83 | 5.35 | 13.23 | 0.16 | -12.20 | 2.22 | 0.73 | 0.13 | 12.95 | 12.81 |
| 189 | Yersinia pestis P-1680-25C | OS | 0.38 | 1.61 | 1.94 | 0.03 | 5.00 | 0.17 | 0.21 | 0.01 | 2.91 | 8.18 |

| | | | | | | | | | | | | |
|---|---|---|---|---|---|---|---|---|---|---|---|---|
| 190 | Yersinia pestis P-1680-37C | LPS | 66.81 | 13.18 | 16.11 | 0.12 | -14.92 | 0.78 | -0.54 | 0.16 | 14.94 | 5.43 |
| 191 | Yersinia pestis I-2377-25C | OS | 0.21 | 2.27 | 1.40 | 0.01 | 4.31 | 0.10 | 0.07 | 0.01 | 3.01 | 8.73 |
| 192 | Yersinia pestis I-2377-37C | LPS | 10.59 | 6.72 | 7.84 | 0.08 | -6.79 | 1.23 | 1.26 | 0.10 | 10.11 | 13.56 |
| 193 | Francisella novicida OPS | OPS | 0.20 | 2.08 | 0.81 | 0.01 | -4.60 | 0.03 | -0.30 | 0.01 | 6.54 | 3.38 |
| 194 | Francisella tularensis OPS | OPS | 0.11 | 2.07 | 0.35 | -0.01 | -13.51 | 0.00 | -0.07 | 0.00 | 0.81 | -1.04 |
| 195 | Klebsiella O1 OPS | OPS | 0.05 | 0.92 | 0.54 | 0.02 | -7.90 | 2.55 | 55.70 | 0.26 | 1.50 | 3.54 |
| 196 | Klebsiella O2a OPS | OPS | 6.59 | 9.87 | 2.43 | 0.02 | -9.25 | 3.84 | -5.92 | 68.24 | 3.21 | 26.42 |
| 197 | Klebsiella O2ac OPS | OPS | 1.01 | 3.24 | 1.09 | 0.00 | -26.05 | 0.05 | 17.41 | 0.00 | 7.62 | 5.88 |
| 198 | Klebsiella O3 OPS | OPS | 30.88 | 35.48 | 19.87 | 0.02 | -4.71 | 2.47 | -0.13 | 0.52 | 1.29 | 4.73 |
| 199 | Klebsiella O4 OPS | OPS | 0.70 | 1.02 | 0.71 | 0.02 | -22.94 | 2.36 | 0.14 | 0.51 | 0.89 | 6.39 |
| 200 | Klebsiella O5 OPS | OPS | 80.71 | 55.45 | 0.76 | 0.01 | -7.61 | 0.07 | 1.32 | 0.03 | 2.57 | 17.93 |
| 201 | Klebsiella O8 OPS | OPS | 1.12 | 1.29 | 1.25 | 0.12 | -16.09 | 1.40 | 95.92 | 0.58 | 1.04 | 1.53 |
| 202 | Klebsiella O12 OPS | OPS | 0.63 | 1.05 | 0.73 | 0.00 | -10.33 | 0.30 | 1.46 | -0.01 | 1.75 | 18.82 |
| 203 | Shigella boydii type 1 | LPSOH | 1.43 | 2.34 | 1.65 | 0.03 | 9.72 | 1.89 | 0.15 | 0.59 | 2.87 | 46.16 |
| 204 | Shigella boydii type 3 | OPS | 0.81 | 1.10 | 0.26 | 0.14 | -2.71 | 2.91 | -0.24 | 2.45 | 0.30 | 2.76 |
| 205 | Shigella boydii type 5 | OPS | 0.13 | 0.94 | 0.37 | 0.02 | -6.51 | 34.42 | 0.06 | 60.58 | 0.29 | 3.75 |
| 206 | Shigella boydii type 9 | OPS | 0.68 | 0.56 | 1.53 | 0.01 | -10.73 | 1.39 | 3.68 | 0.03 | 9.32 | 8.44 |
| 207 | Shigella boydii type 11 | OPS | 2.64 | 3.64 | 0.28 | 0.00 | -15.44 | 0.20 | 0.01 | 0.03 | 0.67 | 7.43 |
| 208 | Shigella boydii type 12 | OPS | 31.56 | 21.01 | 0.57 | -0.01 | -8.20 | 0.46 | 0.22 | 0.01 | 2.44 | 1.13 |
| 209 | Shigella boydii type 15 | OPS | 2.37 | 2.92 | 0.21 | 0.00 | -6.56 | 0.08 | 0.16 | 0.02 | 0.72 | 4.59 |
| 210 | Shigella boydii type 16 | OPS | 1.56 | 1.76 | 0.62 | 0.03 | 7.73 | 1.53 | 0.05 | 0.01 | 1.28 | 21.49 |
| 211 | Shigella boydii type 17 | OPS | 0.29 | 1.06 | 0.49 | 0.02 | 11.68 | 0.17 | 1.55 | 0.25 | 0.74 | 4.02 |
| 212 | Shigella boydii type 18 | OPS | 1.75 | 1.79 | 0.51 | 0.00 | 21.40 | 0.64 | 0.05 | 0.01 | 0.64 | 15.12 |
| 213 | Escherichia coli O49 | OPS | 19.95 | 15.76 | 0.38 | 0.00 | -9.72 | 0.43 | 0.36 | 0.02 | 1.14 | 5.61 |
| 214 | Escherichia coli O52 | OPS | 0.53 | 9.55 | 0.76 | 0.00 | -1.76 | 0.27 | 0.35 | 0.17 | 0.83 | 3.37 |
| 215 | Escherichia coli O58 | OPS | 2.15 | 9.09 | 0.40 | 0.01 | -14.80 | 0.18 | 0.18 | -0.03 | 2.49 | 5.56 |
| 216 | Escherichia coli O61 | LPSOH | 0.57 | 1.41 | 1.77 | 0.03 | -2.22 | 1.08 | 0.30 | 0.04 | 2.55 | 51.85 |
| 217 | Escherichia coli O73 | OPS | 3.40 | 21.29 | 0.42 | 0.05 | -4.38 | 0.05 | 0.01 | 0.01 | 0.30 | 6.19 |
| 218 | Escherichia coli O112ab | OPS | 10.01 | 8.60 | 1.00 | 0.01 | -9.34 | 0.96 | 0.19 | 0.14 | 1.39 | 7.14 |
| 219 | Escherichia coli O118 | OPS | 0.15 | 0.94 | 0.49 | 0.00 | -11.77 | 0.17 | 0.13 | 0.02 | 0.63 | 1.40 |
| 220 | Escherichia coli O125 | OPS | 0.60 | 1.37 | 0.44 | -0.01 | -8.19 | 0.21 | 1.96 | 5.21 | 0.59 | 1.45 |
| 221 | Escherichia coli O151 | OPS | 1.98 | 1.76 | 1.23 | 0.03 | -4.31 | 0.58 | 0.19 | 0.08 | 1.14 | 3.56 |
| 222 | Escherichia coli O168 | OPS | 1.64 | 1.32 | 0.80 | 0.01 | -1.01 | 0.84 | 0.01 | 0.08 | 1.26 | 7.64 |
| 223 | Shigella dysenteriae type 2 | LPSOH | 0.74 | 1.27 | 1.75 | 0.03 | 20.93 | 1.06 | 0.17 | 0.12 | 1.92 | 48.32 |
| 224 | Shigella dysenteriae type 4 | OPS | 1.45 | 2.11 | 0.15 | 0.00 | -5.61 | 0.09 | 0.04 | 0.06 | 0.71 | 2.41 |
| 225 | Shigella dysenteriae type 5 | OPS | 7.52 | 1.49 | 1.47 | 0.21 | -10.89 | 0.27 | 0.01 | 0.02 | 1.45 | 8.90 |
| 226 | Shigella dysenteriae type 6 SR-strain | SR-strain | -0.03 | 1.86 | 0.51 | 0.06 | 1.36 | 0.08 | 0.01 | 0.03 | 0.72 | 7.56 |
| 227 | Shigella dysenteriae type 7 | OPS | 3.30 | 2.28 | 0.09 | 0.18 | -15.19 | 0.13 | 0.06 | 0.05 | 0.97 | 2.40 |
| 228 | Shigella dysenteriae type 8 (Russian) | OPS | 0.10 | 13.31 | 1.25 | 0.10 | -3.52 | 0.13 | 0.02 | 0.06 | 1.15 | 2.85 |
| 229 | Shigella dysenteriae type 9 | OPS | 1.12 | 0.92 | 0.39 | 0.25 | -10.07 | 0.05 | -0.05 | 0.04 | 0.73 | 7.12 |
| 231 | Escherichia coli O111:B4 LPS- solution at 1 mg/mL | L5293-2ML (LPS) (Sigma) | -0.02 | 0.07 | 0.40 | 0.18 | 19.40 | 0.28 | 0.01 | 0.12 | 0.52 | 5.52 |
| 232 | Escherichia coli O26:B6 LPS- solution at 1 mg/mL | L5543-2ML (LPS) (Sigma) | -0.02 | 0.08 | 0.29 | 0.05 | 17.41 | 0.17 | 0.01 | 0.15 | 0.50 | 0.63 |
| 233 | Escherichia coli O55:B5 LPS- solution at 1 mg/mL | L5418-2ML (LPS) (Sigma) | 0.12 | 0.09 | 0.78 | 0.13 | 4.75 | 22.44 | 45.86 | 88.97 | 0.89 | 3.05 |
| 234 | Escherichia coli O127:B8 LPS- solution at 1 mg/mL | L5668-2ML (LPS) (Sigma) | -0.11 | -0.01 | -0.29 | 0.69 | 18.78 | 0.25 | 3.20 | 22.92 | 0.00 | 3.44 |
| 235 | Streptococcus pneumoniae type 1 (Danish type 1) | 161-X // Capsular PS | 0.12 | 0.18 | 1.60 | 72.87 | 0.56 | 0.47 | 5.31 | 0.24 | 2.18 | 16.68 |
| 236 | Streptococcus pneumoniae type 2 (Danish type 2) | 165-X// Capsular PS | 0.05 | 0.01 | 0.03 | 50.18 | -18.25 | 0.05 | -0.09 | 0.03 | -0.24 | 4.15 |
| 237 | Streptococcus pneumoniae type 3 (Danish type 3) | 169-X// Capsular PS | 0.17 | 1.31 | 0.94 | 0.52 | -15.73 | 0.10 | 0.93 | 0.05 | 1.38 | 21.06 |
| 238 | Streptococcus pneumoniae type 4 (Danish type 4) | 173-X// Capsular PS | -0.07 | -0.01 | -0.09 | 62.72 | 14.28 | 0.26 | -0.02 | 0.04 | 0.36 | 17.13 |
| 239 | Streptococcus pneumoniae type 5 (Danish type 5) | 177-X// Capsular PS | 0.03 | 0.61 | 0.35 | 75.24 | -0.03 | 0.17 | 0.24 | 0.31 | 0.76 | 8.86 |
| 240 | Streptococcus pneumoniae type 8 (Danish type 8) | 185-X// Capsular PS | -0.02 | 0.02 | 4.19 | 43.13 | 4.11 | 0.66 | -0.19 | 0.03 | -0.35 | 21.66 |
| 241 | Streptococcus pneumoniae type 9 (Danish type 9N) | 189-X// Capsular PS | -0.03 | 0.21 | -0.37 | 56.89 | -9.38 | 0.06 | -0.02 | 0.03 | 1.12 | 3.13 |
| 242 | Streptococcus pneumoniae type 12 (Danish type 12F) | 193-X// Capsular PS | -0.08 | 0.28 | 0.70 | 69.96 | -8.51 | 0.07 | 0.03 | 0.05 | 1.20 | -3.67 |
| 243 | Streptococcus pneumoniae type 14 (Danish type 14) | 197-X// Capsular PS | 0.16 | 0.40 | -0.40 | 12.63 | 8.10 | 66.31 | 12.03 | 85.30 | 1.16 | 16.03 |
| 244 | Streptococcus pneumoniae type 17 (Danish type 17F) | 201-X// Capsular PS | 0.01 | 0.19 | 0.29 | 92.51 | 8.10 | 0.19 | -0.03 | 0.07 | 0.46 | 5.69 |
| 245 | Streptococcus pneumoniae type 19 (Danish type 19F) | 205-X// Capsular PS | -0.04 | 0.29 | 0.15 | 68.89 | -4.77 | 0.07 | 0.23 | 0.44 | 1.04 | 4.92 |
| 246 | Streptococcus pneumoniae type 20 (Danish type 20) | 209-X// Capsular PS | 0.02 | 0.45 | 0.82 | 82.61 | -4.60 | 0.09 | 0.88 | 0.21 | 0.72 | -0.25 |
| 247 | Streptococcus pneumoniae type 22 (Danish type 22F) | 213-X// Capsular PS | -0.04 | 0.19 | 0.87 | 65.43 | -15.82 | 0.07 | 0.25 | 0.16 | 1.26 | 2.27 |
| 248 | Streptococcus pneumoniae type 23 (Danish type 23F) | 217-X// Capsular PS | 0.03 | 0.21 | 0.47 | 8.64 | -17.29 | 0.03 | 0.06 | 0.01 | 0.46 | -5.36 |
| 249 | Streptococcus pneumoniae type 26 (Danish type 6B) | 225-X// Capsular PS | 0.05 | 0.27 | 0.44 | 78.63 | 6.45 | 0.32 | 0.32 | 0.08 | 0.40 | 2.92 |
| 250 | Streptococcus pneumoniae type 34 (Danish type 10A) | 229-X// Capsular PS | 0.59 | 1.80 | 6.39 | 89.04 | -4.48 | 0.10 | 1.89 | 2.56 | 31.28 | 6.57 |
| 251 | Streptococcus pneumoniae type 43 (Danish type 11A) | 233-X// Capsular PS | 0.05 | 0.31 | 0.69 | 76.48 | -5.76 | 87.95 | 60.26 | 89.37 | 0.73 | 2.61 |

| | | | | | | | | | | | |
|---|---|---|---|---|---|---|---|---|---|---|---|
| 252 | Streptococcus pneumoniae type 51 (Danish type 7F) | 237-X// Capsular PS | 0.23 | 0.38 | 0.60 | 87.57 | -9.25 | 0.09 | -0.02 | 0.27 | 1.29 | 3.15 |
| 253 | Streptococcus pneumoniae type 54 (Danish type 15B) | 241-X// Capsular PS | 0.04 | -0.02 | 0.42 | 71.32 | -13.78 | 67.88 | 0.76 | 1.62 | 0.88 | -1.74 |
| 254 | Streptococcus pneumoniae type 56 (Danish type 18C) | 245-X// Capsular PS | -0.01 | 0.19 | 0.02 | 95.27 | -20.01 | 0.04 | -0.10 | 0.02 | 1.21 | 2.16 |
| 255 | Streptococcus pneumoniae type 57 (Danish type 19A) | 249-X// Capsular PS | 0.07 | 1.52 | 0.63 | 88.67 | 7.95 | 0.19 | 0.56 | 1.59 | 1.12 | 11.88 |
| 256 | Streptococcus pneumoniae type 68 (Danish type 9V) | 253-X// Capsular PS | 0.04 | 0.39 | 0.13 | 100.00 | 1.33 | 0.24 | 0.14 | 0.26 | 0.76 | 9.41 |
| 257 | Streptococcus pneumoniae type 70 (Danish type 33F) | 257-X// Capsular PS | 0.91 | 1.18 | -0.71 | 62.14 | -11.09 | 14.05 | 100.00 | 100.00 | 0.81 | 1.35 |
| 258 | Yersinia pestis KM218-6C | OS | 0.25 | 2.39 | 1.85 | 0.03 | -4.16 | 0.51 | -0.07 | 0.04 | 3.82 | 22.76 |
| 259 | Yersinia pestis KM260(11)-yjhW-6C | OS | 0.31 | 0.92 | 9.25 | 0.06 | 1.44 | 0.20 | 16.09 | 2.23 | 1.70 | 8.13 |
| 260 | Yersinia pestis KM260(11)-wabD/waaL | OS | 0.11 | 1.61 | 0.66 | 0.02 | 0.83 | 0.09 | 0.11 | 0.03 | 4.35 | 7.86 |
| 261 | Yersinia pestis KM260(11)-wabC/waaL | OS | 79.47 | 52.04 | 12.55 | 0.08 | -27.90 | 1.34 | 2.47 | 0.39 | 19.45 | 34.50 |
| 262 | Yersinia pseudotuberculosis 85pCad-37C | OS | -0.21 | 1.36 | 0.66 | 0.03 | 0.21 | 0.17 | 0.09 | 0.03 | 1.35 | 6.85 |
| 263 | Yersinia pseudotuberculosis 85pCad-20C | OS | 0.16 | 0.64 | 1.55 | 0.03 | -4.14 | 0.22 | 0.38 | 0.07 | 0.51 | 10.16 |
| 264 | Yersinia pseudotuberculosis O:2a | PS | 33.49 | 43.51 | 2.28 | 0.02 | 0.18 | 0.21 | 0.10 | 0.04 | 3.51 | 7.48 |
| 265 | Yersinia pseudotuberculosis O:2a-dhmA | PS | 2.48 | 10.48 | 1.06 | -0.02 | -4.34 | 0.10 | 0.09 | 0.01 | 1.55 | 4.90 |
| 266 | Yersinia pseudotuberculosis O:2c | PS | 3.95 | 4.89 | 11.46 | 0.06 | -5.31 | 0.97 | 1.24 | 0.08 | 2.79 | 24.14 |
| 267 | Yersinia pseudotuberculosis O:3 | PS | 3.38 | 3.78 | 0.39 | 0.00 | -10.38 | 0.12 | 0.03 | 0.00 | 0.51 | 3.54 |
| 268 | Yersinia pseudotuberculosis O:4b | PS | 6.75 | 15.75 | 0.79 | 0.01 | -7.70 | 0.16 | 0.20 | 0.04 | 1.03 | 3.10 |
| 269 | Proteus vulgaris O2 (OX2) | PS | 5.88 | 0.61 | 0.50 | 0.01 | -4.63 | 0.77 | 0.33 | 0.18 | 0.77 | 2.74 |
| 270 | Proteus mirabilis O3ab (S1959) | PS | 0.89 | 4.41 | 16.29 | 1.98 | 36.13 | 0.47 | 0.48 | 2.04 | 21.60 | 5.81 |
| 271 | Proteus mirabilis O5 (PrK 12/57) | PS | 1.35 | 1.74 | 1.22 | 0.05 | 1.79 | 0.52 | 1.12 | 0.12 | 1.83 | 13.25 |
| 272 | Proteus mirabilis O9 (PrK 18/57) | PS | 0.29 | 0.92 | 0.06 | 0.26 | -2.77 | 0.18 | 0.06 | 0.03 | 2.53 | 3.32 |
| 273 | Proteus mirabilis O11 (9B-m) | PS | 10.26 | 0.78 | 2.66 | 0.15 | 2.80 | 0.15 | 0.14 | 0.05 | 2.59 | 8.02 |
| 274 | Proteus penneri O17 (16) | PS | 0.32 | 5.23 | 0.67 | 0.03 | 5.57 | 0.14 | 0.03 | 0.01 | 1.45 | 3.85 |
| 275 | Proteus mirabilis O18 (PrK 34/57) | LPSOH | 0.75 | 1.14 | 9.14 | 58.04 | 4.68 | 0.59 | 31.79 | 0.62 | 9.06 | 19.97 |
| 276 | Proteus mirabilis O20 (PrK 38/57) | LPSOH | 0.67 | 1.48 | 4.88 | 0.18 | 22.45 | 7.66 | 1.78 | 12.71 | 10.36 | 64.43 |
| 277 | Proteus penneri O31ab (28) | PS | 1.35 | 0.87 | 2.08 | 0.45 | 2.93 | 0.36 | 0.06 | 0.07 | 0.82 | 12.21 |
| 278 | Proteus mirabilis O33 (D52) | PS | 0.55 | 2.29 | 4.19 | 0.32 | -1.87 | 6.74 | 40.28 | 4.54 | 12.80 | 51.17 |
| 279 | Proteus mirabilis O43 (PrK 69/57) | PS | 1.82 | 1.75 | 1.28 | 0.06 | 1.27 | 0.12 | 0.01 | 0.03 | 1.95 | 9.47 |
| 280 | Proteus vulgaris O47 (PrK 73/57) | Not stated | 0.48 | 1.18 | 0.38 | 0.00 | -6.56 | 39.21 | 51.05 | 6.75 | 2.49 | 4.01 |
| 281 | Proteus mirabilis O49 (PrK 75/57) | PS | 1.89 | 1.87 | 0.74 | 0.07 | 3.89 | 0.54 | 0.24 | 0.04 | 2.43 | 4.31 |
| 282 | Proteus mirabilis O54ab (OE) | PS | 0.21 | 0.68 | 1.61 | 0.23 | -5.62 | 0.15 | 0.32 | 0.02 | 2.96 | 1.58 |
| 283 | Proteus penneri O73ac (75) | PS | 0.59 | 1.34 | 1.23 | 0.02 | 27.22 | 0.32 | 46.02 | 1.19 | 1.69 | 15.61 |
| 284 | Proteus vulgaris O76 (HSC438) | PS | -0.02 | 0.39 | 0.56 | 0.18 | -2.00 | 0.05 | -0.05 | 0.01 | 1.88 | -3.09 |
| 285 | Shigella flexneri type 1a | PS | 2.93 | 33.43 | 0.54 | 0.01 | -11.77 | 0.06 | 0.04 | 0.01 | 0.89 | 2.23 |
| 286 | Shigella flexneri type 1b | PS | 26.09 | 41.82 | 1.28 | 0.06 | -8.14 | 0.06 | 0.16 | 0.05 | 2.39 | 2.59 |
| 287 | Shigella flexneri type 2a | PS | 0.16 | 1.89 | 0.51 | 0.00 | -11.75 | 0.25 | -0.01 | 0.04 | 0.99 | 1.40 |
| 288 | Shigella flexneri type 2b | PS | 33.46 | 63.55 | 0.54 | 0.04 | -14.71 | 0.01 | 0.18 | 0.01 | 1.66 | 2.20 |
| 289 | Shigella flexneri type 3a | PS | 0.63 | 100.00 | 1.14 | 0.03 | -14.74 | 0.12 | 0.13 | 0.02 | 1.77 | -3.89 |
| 290 | Shigella flexneri type 3b | PS | 16.29 | 16.37 | 0.86 | 0.02 | 13.91 | 0.31 | 0.08 | 0.09 | 2.57 | 22.18 |
| 291 | Shigella flexneri type 4a | PS | 20.72 | 28.62 | 43.07 | 2.54 | 32.49 | 1.60 | 2.05 | 2.36 | 83.06 | 7.26 |
| 292 | Shigella flexneri type 4b | PS | 14.26 | 15.73 | 0.15 | 0.00 | -11.28 | 0.03 | 0.05 | 0.06 | 0.84 | 1.95 |
| 293 | Shigella flexneri type 5b | PS | 11.96 | 52.64 | -2.22 | 0.01 | -9.04 | 0.03 | 0.20 | 0.04 | 1.34 | 4.87 |
| 294 | Shigella flexneri type 6a | PS | 33.97 | 25.27 | 0.73 | 0.02 | -5.71 | 0.22 | 0.14 | 0.03 | 1.60 | 4.41 |
| 295 | Shigella flexneri type 6 | PS | 5.80 | 5.19 | -1.57 | 0.03 | -4.38 | 0.16 | 0.40 | 0.02 | 1.45 | 6.40 |
| 296 | Shigella flexneri type X | PS | 3.10 | 76.99 | 0.02 | 0.03 | -17.55 | 0.02 | 0.11 | 0.05 | 1.43 | 0.40 |
| 297 | Shigella dysenteriae type 1 | PS | 1.88 | 0.88 | 0.30 | 0.00 | -12.64 | 0.07 | 0.13 | 0.02 | 1.88 | 0.13 |
| 298 | Shigella boydii type 6 | PS | 0.33 | 0.70 | 0.66 | 0.00 | 6.19 | 0.21 | 0.05 | 0.01 | 1.16 | 5.42 |
| 299 | Shigella boydii type 7 | PS | 11.19 | 10.37 | 2.59 | 0.07 | 6.39 | 0.90 | 0.15 | 0.09 | 3.84 | 17.69 |
| 300 | Shigella boydii type 8 | PS | 1.15 | 1.31 | 0.46 | 0.05 | -10.64 | 0.11 | -0.07 | 0.06 | 0.82 | 0.44 |
| 301 | Shigella boydii type 13 | LPSOH | 13.74 | 11.91 | 3.08 | 0.08 | 20.80 | 1.59 | 0.43 | 0.17 | 4.51 | 59.51 |
| 302 | Shigella boydii type 14 | LPSOH | 0.42 | 5.86 | 4.48 | 0.08 | 22.57 | 0.90 | 0.24 | 0.14 | 5.01 | 20.49 |
| 303 | Escherichia coli O71 | PS | 11.73 | 8.30 | 0.44 | 0.01 | -12.63 | 0.02 | 0.07 | 0.02 | 0.58 | 0.76 |
| 304 | Escherichia coli O85 | PS | 8.87 | 7.52 | 0.40 | 0.06 | -12.43 | 0.16 | 0.02 | 0.07 | 0.89 | 2.30 |
| 305 | Escherichia coli O99 | PS | 49.45 | 39.68 | 1.26 | 0.00 | -3.26 | 0.09 | 0.01 | 0.00 | 1.61 | 1.58 |
| 306 | Escherichia coli O145 | LPSOH | 0.95 | 0.94 | 4.05 | 0.08 | 9.87 | 1.89 | 0.34 | 0.18 | 8.93 | 41.67 |
| 307 | Escherichia coli O107 | PS | 56.84 | 29.82 | -0.87 | 0.02 | 5.54 | 0.30 | 0.19 | 0.11 | 3.22 | 23.47 |
| 308 | Salmonella enterica O17 | PS | 1.88 | 1.35 | 0.43 | 0.03 | -8.82 | 0.05 | 0.03 | 0.03 | 1.96 | -0.42 |
| 309 | Salmonella enterica O28 | PS | 21.01 | 10.94 | 1.09 | 0.02 | -11.15 | 0.00 | 0.17 | 0.03 | 1.04 | -2.34 |
| 310 | Salmonella enterica O47 | PS | 0.20 | 0.65 | -0.09 | 0.03 | 2.66 | 0.47 | 0.05 | 0.02 | 1.08 | 4.00 |
| 311 | Salmonella enterica O55 | PS | 1.17 | 0.74 | 1.44 | 0.02 | -9.59 | 0.08 | 0.06 | 0.00 | 4.57 | -1.55 |
| 312 | Escherichia coli K92 | CPS | 0.02 | 0.98 | 0.70 | 0.00 | -3.95 | 0.05 | 0.02 | 0.01 | 0.68 | 3.81 |

| | | | | | | | | | | | | |
|---|---|---|---|---|---|---|---|---|---|---|---|---|
| 313 | Escherichia coli K5 | CPS | 0.28 | 1.53 | 2.73 | 0.02 | -7.45 | 0.09 | 0.01 | 0.02 | 3.48 | 18.28 |
| 314 | Escherichia coli K13 | CPS | 0.03 | 3.39 | 1.61 | 0.05 | 4.77 | 0.26 | 0.05 | 0.06 | 1.98 | 14.13 |
| 315 | Neisseria meningitidis Group C | CPS | 0.06 | 1.13 | -0.19 | 0.01 | -4.83 | 0.12 | 0.23 | 0.02 | 1.44 | 18.07 |
| 316 | Davanat | | 0.16 | 1.18 | -0.01 | 0.03 | -13.16 | 0.06 | -0.07 | 0.02 | 1.69 | 4.68 |
| 317 | Laminarin | | 1.30 | 1.54 | 16.20 | 0.03 | 0.36 | 0.05 | 5.74 | 0.18 | 25.30 | 10.07 |
| 318 | Yeast Mannan | | 57.26 | 82.30 | 11.81 | 0.33 | 2.68 | 0.11 | 0.26 | 0.05 | 16.29 | 8.56 |
| 319 | Escherichia coli O86 | | 0.22 | 0.67 | 0.80 | 0.02 | -7.34 | 0.74 | 70.83 | 82.11 | 2.94 | 4.03 |
| 320 | Galactomannan DAVANT (160102) Pro-Pharmacenti | | 0.20 | 0.89 | 2.12 | 0.04 | -7.76 | 0.03 | 0.00 | 0.02 | 1.92 | 5.87 |
| 321 | Yeast Mannan Sigma M-3640 | | 38.29 | 80.96 | 10.85 | 0.29 | 5.43 | 0.17 | 0.03 | 0.02 | 26.80 | 12.57 |
| 322 | 1-2 Mannan Acetobacter methanolieus MB135 | | 1.59 | 3.22 | 7.21 | 0.03 | 0.75 | 0.11 | 4.55 | 0.15 | 39.81 | 8.23 |

### SI-Spreadsheet 2

| | | | Lectins | | | | | | | | | |
|---|---|---|---|---|---|---|---|---|---|---|---|---|
| BPS# | BACTERIA / STRAIN | NAME / STRUCTURE /Cat.No. | MBL | ΔN SPD | CL-K1 | CRP | FicH | Gal-3 | Gal-4 | Gal-8 | PTX3 | ZG16P |
| 1 | Providencia stuartii O49 | PO49 Core-linked | 56.25 | 270.25 | 13.25 | 3 | -177.75 | 27 | 5.25 | 3.5 | 4.25 | 93.5 |
| 2 | Providencia stuartii O52 | PO52 Core-linked | 1105.25 | 453.5 | 34.5 | 335.5 | -125.5 | 62.25 | 46.75 | 70 | 94.5 | 544.5 |
| 3 | Pseudomonas aerguinosa O4 (Habs serotype 4) | PO4 Core-linked | 110.25 | 143.75 | 14.75 | -3.5 | -198.5 | 4.75 | -77 | 1.5 | 16 | 149 |
| 4 | Pseudomonas aerguinosa O1 (Fisher immunotype 4) | PO1 Core-linked | 25.25 | 97.25 | -0.5 | 13.25 | -169.5 | 9.5 | 51 | -2.25 | 9.5 | 2 |
| 5 | Pseudomonas aerguinosa O2 (Fisher immunotype 3) | PO2 Core-linked | 94.25 | 180.75 | 13.25 | 54 | -96.75 | 389.75 | 29 | 53.25 | 111.25 | 302.75 |
| 6 | Pseudomonas aerguinosa O13 (Sandvik serotype II) | PO13 Core-linked | -7 | -8 | 2 | 6.25 | -73 | 47 | -41 | 151.25 | -4.25 | 236.75 |
| 7 | Pseudomonas aerguinosa O9 (9a, 9b, 9d) | PO9 Core-linked | 110 | 21.75 | 282 | -12.25 | -703 | 5.75 | 340 | -11.25 | 35.75 | -40.25 |
| 8 | Pseudomonas aerguinosa O6a (Habs serotype6, fraction IIa) | PO6a Core-linked-O-unit | 19.5 | 223.25 | 2 | 1.75 | -54 | 788.5 | -4 | 19.5 | 7 | 269.5 |
| 9 | Pseudomonas aerguinosa O6a (Habs serotype6, fraction IIb) | PO6a unsubstituted core | 37.5 | 405.5 | 16.5 | 0 | 13.25 | 27.5 | 12.25 | -9.25 | 16 | 207.75 |
| 12 | Salmonella typhimurium SL 11881 (Re mut) | LPS-L9516 | 36058.75 | 3109 | 98.5 | -7.25 | 1010.5 | 40.75 | 287.25 | 5 | 154.5 | 1059 |
| 13 | Salmonella typhimurium TV 119 (Ra mut) | LPS-L6016 | 1302.25 | 514.5 | 36 | 16.75 | 300.5 | 77 | 298.75 | 22.25 | 80.5 | 2998.25 |
| 14 | Salmonella typhimurium SL 684 (Rc mut) | LPS-L5891 | 921.5 | 1678.75 | 63 | 0.25 | 495 | 21 | 366.75 | 28 | 173.5 | 1765 |
| 15 | Pseudomonas aerguinosa O10 | L8643 | 57.5 | 251.25 | -0.75 | 9.5 | 78.5 | 939.5 | 14 | 72.5 | 10.25 | 381.75 |
| 16 | Salmonella typhimurium dodeca saccharide | 4809 | 14 | 168 | 7.5 | 4.25 | 138.5 | 113.25 | 46 | 3 | 15.5 | 217.25 |
| 17 | Salmonella enteritidis dodeca saccharide | 1262 | 46.5 | 177 | 16 | -1 | 186 | 54.75 | 32 | 4.25 | 10.5 | 291.5 |
| 18 | Salmonella typhimurium LPS | L2262 | 1813.5 | 361 | 390.5 | 27 | 340.75 | 351.5 | 376.5 | 286.5 | 46.5 | 1308 |
| 20 | Serratia marcescens LPS | L6136 | 11792 | 516.75 | 31.5 | 6.25 | 1149 | 287.25 | 151.25 | 24.5 | 46 | 994 |
| 22 | Escherichia coli K235 LPS | L2143 | 471.25 | 5767.75 | 946 | 24.75 | 95 | 207.75 | 351.5 | 65.5 | 60.75 | 1190.75 |
| 23 | Escherichia coli O128-B12 LPS | L2755 | 862.25 | 267 | 2316.25 | 4.25 | 712.75 | 256 | 11121.75 | 740.25 | 68.5 | 1242.25 |
| 25 | Salmonella enterica abortus equi LPS | L5886 | 79.25 | 209.5 | 38 | 28.5 | -93 | 258.25 | 128.25 | 114 | 33.75 | 1637 |
| 26 | Salmonella typhosa LPS | L2387 | 1481.25 | 349 | 524.5 | 2.5 | 689.25 | 299 | 250.75 | 206.25 | 71.75 | 820.75 |
| 27 | Salmonella enteritidis LPS | L2012 | 1269.75 | 112.5 | 48.75 | -12.5 | 520.25 | 226.25 | 62.5 | 39.25 | 41.5 | 894.75 |
| 28 | Shigella bodyii type2 | | 373.5 | 160.75 | 29.25 | 6 | -110 | 630.75 | 118 | 64 | 44.5 | 912.5 |
| 29 | Shigella bodyii type4 | | 1744 | 541.75 | 101.25 | 216 | -54.75 | 29 | 26 | 13 | 97.25 | 197.5 |
| 30 | Shigella bodyii type10 | | 9662.75 | 232.5 | 891 | -5.25 | -123.75 | 185.25 | 12.5 | 19.25 | 17.75 | 336.75 |
| 31 | Shigella dysennteriae type 3 | | 676.5 | 275.5 | 107.25 | 286.25 | 185 | 401.25 | -5.5 | 26.75 | 86.5 | 955.25 |
| 32 | Shigella dysennteriae type 8 (batch 12) | | 9535.25 | 234 | 32 | 29.25 | -53.25 | 55.5 | -21.5 | 14.5 | 102.75 | 177.75 |
| 33 | Shigella dysennteriae type 11 | | 23 | 107.75 | -14.5 | 56.5 | -47.25 | 171.75 | 29.25 | 13.25 | 13.5 | 569.5 |
| 34 | Shigella dysennteriae type 13 | | 200 | 825.25 | 17.25 | 77.75 | 80.5 | 65.25 | -18.5 | 38.25 | 26.25 | 426.5 |
| 35 | Escherichia coli O29 | | 164 | 118.75 | 7 | 8.5 | 82.5 | 27.75 | 1 | 7.75 | 7.25 | 423.5 |
| 36 | Escherichia coli O40 | | 0.5 | 310.5 | 4.5 | 5.5 | 26 | 79.5 | 12793 | 55977.75 | 4 | 274.25 |
| 37 | Escherichia coli O106 | | 24766.25 | 4123.5 | 25.75 | 117.25 | -133 | 22.25 | -4.75 | 44.5 | 54.25 | 345.5 |
| 38 | Escherichia coli O130 | | 453.5 | 638.5 | 12 | 109.25 | 13.75 | 19.5 | 66.25 | 12 | 17.25 | 358.5 |
| 39 | Escherichia coli O148 | | 1643.75 | 265.75 | 10.5 | 2.75 | -270 | 24.25 | 9.5 | 6.75 | 18.5 | 655.25 |
| 40 | Escherichia coli O150 | | 294 | 374.25 | 13.75 | 31.75 | -31 | 23.25 | 42.75 | 37.75 | 21.25 | 485.25 |
| 41 | Escherichia coli O180 | | 14664.5 | 1075 | 6.25 | 54.75 | -249.75 | 91.25 | 101 | 47.75 | 26.75 | 541.25 |
| 42 | Proteus mirabilis O3a, 3c (G1) | | -7 | 20.25 | -1.25 | 919 | 1689 | 600.25 | 37.75 | 1458.75 | 0.75 | 1070 |
| 43 | Proteus mirabilis O8 (TG326) | | 112.25 | 129.75 | 14 | -2.75 | -20 | 24.5 | 88 | 19.5 | 14.75 | 169.75 |
| 44 | Proteus mirabilis O10 (HJ4320) | | -1.75 | -2.25 | 3 | 180.25 | 13.5 | 44 | -30.25 | 14 | -0.25 | 245.5 |
| 45 | Proteus mirabilis O29a, 29b (2002) | | 38.75 | 187.25 | 14.5 | 42.5 | 313.25 | 108 | 36 | 41.5 | 9 | 552.5 |
| 46 | Proteus mirabilis O50 (TG332) | | -9.5 | 1.75 | -0.25 | 30 | 45 | 30.5 | 43.25 | 1.5 | -5.25 | 266 |
| 47 | Proteus mirabilis O54a, 54b (10704) | | 129.5 | 216.75 | 32.5 | 51.5 | -139.25 | 9.5 | 147 | 126.25 | 49.75 | 402.75 |
| 48 | Proteus mirabilis O57 (TG319) | | -44 | -1.25 | -4 | 31 | -6.25 | 68.75 | -11.75 | 80.5 | -4.5 | 442.25 |
| 49 | Proteus penneri O8 (106) | | -23.5 | 78.25 | -65.75 | -11.25 | -137.5 | 45.25 | 32.5 | -4.75 | 3.75 | 120.5 |
| 50 | Proteus penneri O64a, 64b, 64d (39) | | 48 | 166.75 | 7 | 13.25 | -126.5 | 26.25 | 32 | 8 | 10 | 73.75 |
| 51 | Proteus penneri O66 (2) | | 769 | 107.75 | 20 | 1.5 | -232 | 81.75 | -4.75 | 18.75 | 5 | -22.5 |
| 52 | Proteus penneri O69 (25) | | 371.25 | 307 | 149 | 287.5 | 799.75 | 211.25 | 366.25 | 298.25 | 140.25 | 350.25 |
| 53 | Proteus penneri O71 (42) | | 9691.25 | 1454 | 51.25 | 34 | 35.5 | 1215 | 436.5 | 194.5 | 44.75 | 925.25 |
| 54 | Proteus penneri O72a, 72b (4) | | 102.75 | 503.75 | 9.25 | 111.5 | -9.75 | 65.5 | 68 | 9.5 | 13.5 | 151.25 |
| 55 | Pseudomonas aerguinosa O2 (2a),2d,2f | IATS 10 , OPS | 169.25 | 67.25 | 1.25 | 59.75 | -248 | 43.75 | 54.75 | 139.5 | 13.5 | 1 |
| 56 | Pseudomonas aerguinosa O2 2a,2b | IATS 16 OPS | 15.25 | 285.25 | 13.25 | 203.5 | -47.25 | 106.75 | 136.25 | 30 | 39.75 | 441.75 |
| 57 | Pseudomonas aerguinosa O2 2a,2b,2e | IATS NO, OPS | 93 | 154.25 | 14 | 36.5 | -158.75 | 22 | 162.25 | 29.25 | 24 | 80.25 |
| 58 | Pseudomonas aerguinosa O2 2a,2d | IATS 5 OPS | 42.75 | 168.5 | -5.5 | 734.25 | 30.25 | 24 | 51 | 36 | 14.75 | 261 |
| 59 | Pseudomonas aerguinosa O2 Immuno 7 | IATS 18, OPS | 53.5 | 76 | 9.25 | 90 | -78 | 42.5 | 22.5 | 435.25 | 14.75 | 16.75 |
| 60 | Pseudomonas aerguinosa O3 3a,3b | IATS NO, OPS | -56.75 | 71 | 3.5 | 831 | -333 | 10.5 | 89.5 | 23.75 | 10.75 | -84.75 |
| 61 | Pseudomonas aerguinosa O3 3a,3b,3c | IATS 3, OPS | 77.5 | 50.75 | 6.75 | 8 | -454.5 | 60.25 | 70.5 | 59.75 | 10 | -18.5 |
| 62 | Pseudomonas aerguinosa O3 3a,3d | IATS NO, OPS | 0 | 157.5 | 11.75 | 823.25 | -183.75 | 37.5 | 75 | 34.25 | 23.5 | 108.75 |
| 63 | Pseudomonas aerguinosa O4 4a,4c | IATS NO, OPS | 186.75 | 496.5 | 21.25 | 22.75 | -43.25 | 87.5 | 118 | 19.75 | 81.25 | 305.5 |
| 64 | Pseudomonas aerguinosa O6 6a | IATS 6, OPS | 1.5 | 113.75 | 2.75 | 1424 | -264.25 | 29.25 | -7.75 | 32 | 14.75 | -112 |

| | | | | | | | | | | | | |
|---|---|---|---|---|---|---|---|---|---|---|---|---|
| 65 | Pseudomonas aerguinosa O6 6a,6c | IATS NO, OPS | 133.75 | 125.25 | 62.75 | 330.75 | -242.25 | 123 | 39.75 | 125 | 87 | -416 |
| 66 | Pseudomonas aerguinosa O6 Immuno 1 | IATS NO, OPS | 48 | 164 | 32 | 54.75 | -267.75 | 36.5 | 92 | 55.75 | 41.5 | -209.5 |
| 67 | Pseudomonas aerguinosa O7 7a,7b,7c | IATS 7,LPS | 1385.75 | 147 | 420.25 | 90.25 | 460.5 | 463.5 | 3286 | 316 | 1059.25 | 1540.75 |
| 68 | Pseudomonas aerguinosa O7 7a,7b,7d | IATS 8,LPS | 100.25 | 69.75 | 68.75 | 22.25 | 940.25 | 1362.75 | 517.25 | 73.75 | 69.5 | 2699.25 |
| 69 | Pseudomonas aerguinosa O7 7a,7d | IATS NO, LPS | 152.25 | 101.75 | 856.75 | 26.75 | -545.5 | 393.25 | 166.75 | 163.5 | 80.75 | 840.25 |
| 71 | Pseudomonas aerguinosa O10 10a,10b | IATS 10, OPS | 91.5 | 92.25 | -46 | 124.5 | -391.75 | 51.75 | 65.75 | 24.75 | 17.75 | 318.75 |
| 72 | Pseudomonas aerguinosa O10 10a,10c | IATS 19, OPS | -67 | 51 | 14 | 59 | -312.75 | 6.5 | 78.5 | 24.5 | 17.75 | -325.75 |
| 73 | Pseudomonas aerguinosa O11 11a,11b | IATS 11, OPS | 80.25 | 100.25 | 7.75 | -3.25 | -10.5 | 28.75 | 35 | 17.5 | 17.5 | 185.5 |
| 74 | Pseudomonas aerguinosa O12 12 | IATS 12, OPS Habs 12 | -1.25 | 199.75 | 10.25 | 17 | 22.75 | 140.5 | 290.25 | 57.75 | 39.75 | 291.25 |
| 75 | Pseudomonas aerguinosa O13 13a,13c | IATS 14, OPS | 132.25 | 2633.25 | 53.5 | 26.25 | -327.25 | 160.25 | 55.75 | 31.25 | 55.25 | 772 |
| 76 | Pseudomonas aerguinosa O14 14 | IATS 17,OPS Meitert X | 7.75 | 63.5 | 22 | 83 | -442.25 | 32.25 | 301.25 | 29.5 | 11.5 | 4 |
| 77 | Pseudomonas aerguinosa O15 15 | IATS 15, OPS | 90 | 112.25 | 15.75 | 4 | -150.25 | 11.25 | 133.75 | 44.5 | 38.5 | 234.25 |
| 78 | Proteus vulgaris O1 (18984)* | LPS | 59.75 | 135.25 | 87.75 | 1540 | -97.25 | 5124.5 | 7492.75 | 15184 | 66.25 | 629.25 |
| 79 | Proteus vulgaris O4 (PrK 9/57) | OPS | 35 | 116.25 | 3.25 | -2.5 | 35.5 | 23 | 11.25 | 6.5 | 14.25 | 362 |
| 80 | Proteus vulgaris O12 (PrK 25/57) | OPS | 14.75 | 166.5 | 8.5 | 87.75 | 47.25 | 67.5 | 29.75 | 11 | 21.75 | 101 |
| 81 | Proteus vulgaris O13 (8344) | OPS | 132.5 | 119.25 | 11.25 | 411.25 | 148 | 67.25 | 80.5 | 18.75 | 39.25 | 301.25 |
| 82 | Proteus vulgaris O15 (PrK 30/57) | OPS | -0.75 | 2 | -8.25 | 16.5 | -68 | 24 | -76.25 | 16.75 | 0 | 157.5 |
| 83 | Proteus vulgaris O17 (PrK 33/57) | OPS | 368.75 | 182.5 | 115.25 | 42.75 | -155.5 | 1190 | 475.25 | 129.5 | 7.75 | 268.5 |
| 84 | Proteus vulgaris O19a (PrK 37/57) | OPS | 61 | 219.5 | 2.5 | 0 | -197.5 | 36.25 | 53.5 | 4 | 14 | -7.5 |
| 85 | Proteus vulgaris O21 (PrK 39/57)* | LPS | 2492.75 | 196.5 | 157.25 | 158.5 | 46.25 | 1032.25 | 6863.5 | 1791.25 | 276.5 | 1712.5 |
| 86 | Proteus vulgaris O22 (PrK 40/57) | OPS | 80.25 | 188.5 | 4.5 | 0 | -181.25 | 45.25 | 29.5 | 5.75 | 21.5 | 79.25 |
| 88 | Proteus vulgaris O25 (PrK 48/57) | OPS | 154 | 94.25 | 6 | 22.25 | -229.5 | 86 | 14.75 | 11.5 | 38.75 | 453.75 |
| 89 | Proteus vulgaris O34 (4669)* | LPS | 683.25 | 2794 | 153.75 | 11254.5 | 5.25 | 854.75 | 630.5 | 303 | 306 | 1789.5 |
| 90 | Proteus vulgaris O37a,b (PrK 63/57) | OPS | 127 | 368.25 | 25.25 | 96.5 | -52 | 70.75 | 81.75 | 16 | 52.75 | 392.75 |
| 91 | Proteus vulgaris O37a,c (PrK 72/57) | OPS | 661.25 | 345 | -16.25 | 21.5 | -38.25 | 52.5 | 49.5 | 1.75 | 35.25 | 420.75 |
| 92 | Proteus vulgaris O44 (PrK 67/57) | OPS | -102.25 | 101 | 0.25 | 44.75 | -74.75 | 40.75 | 28271 | 9 | 13.75 | 44.75 |
| 93 | Proteus vulgaris O45 (4680) | OPS | 33 | 148.25 | 2.25 | 7.75 | 0.5 | 14160.5 | 29468.75 | 685.5 | 21.75 | 196.5 |
| 94 | Proteus vulgaris O53 (TG 276-10) | OPS | 379 | 148.5 | 61 | 259.75 | -42.75 | 153.5 | 177.75 | 26.75 | 76 | 312 |
| 95 | Proteus vulgaris O54a,54c (TG 103) | OPS | 81.75 | 140 | 28.75 | 37.25 | -138.25 | 83 | 12297.75 | 256.75 | 58.75 | 201 |
| 96 | Proteus vulgaris O55 (TG 155) | OPS | 22.25 | 50.75 | 14 | 138.75 | -203.5 | 25 | 4.25 | 34.25 | 15.5 | 69.75 |
| 97 | Proteus vulgaris O65 (TG 251) | OPS | 9.25 | 152.75 | -25.75 | -25.75 | -209.25 | 1953.75 | 496.75 | 16655.5 | 60.25 | 79 |
| 98 | Proteus mirabilis O6 (PrK 14/57) | OPS | 98.75 | 289.5 | 13 | 2.75 | 46.25 | 61.25 | 1.75 | 11 | 25.5 | 351.75 |
| 99 | Proteus mirabilis O11 (PrK 24/57) | OPS | 741.5 | 125 | -28.5 | 10 | -29 | 16.5 | 163.5 | 1.5 | 32.75 | 249 |
| 100 | Proteus mirabilis O13 (PrK 26/57) | OPS | 1816.25 | 476.25 | 20.25 | 27.5 | 77.5 | 59.5 | 49.75 | 19.5 | 47 | 263.5 |
| 101 | Proteus mirabilis O14a,14b (PrK 29/57) | OPS | 54.25 | 154.75 | -21.75 | 63.25 | 114.75 | 85.25 | 101 | 16.75 | 67.25 | 602.75 |
| 102 | Proteus mirabilis O16 (4652) | OPS | 4685.25 | 1750.75 | 494 | 775.5 | 294.25 | 374.75 | 263 | 412.75 | 1181.75 | 651.5 |
| 103 | Proteus mirabilis O17 (PrK 32/57) | OPS | 833.75 | 896.75 | 513.75 | 663.25 | 1005.5 | 327.75 | 454.75 | 635.5 | 763.75 | 756.75 |
| 104 | Proteus mirabilis O23a,b,d (PrK 42/57) | OPS | 59.5 | 214.5 | 6.25 | 15.5 | 42.25 | 96.5 | 29.25 | 18.75 | 27.25 | 505.75 |
| 106 | Proteus mirabilis O26 (PrK 49/57) | OPS | 1566.75 | 925.25 | 367.75 | 410.75 | 265 | 323.75 | 147 | 290.25 | 403.75 | 599 |
| 107 | Proteus mirabilis O27 (PrK 50/57) | OPS | 1558.25 | 1844.5 | 1439.5 | 1666 | 1065.75 | 1571 | 1147 | 1864.5 | 1993.75 | 1288 |
| 108 | Proteus mirabilis O28 (PrK 51/57) | OPS | 544.5 | 1029 | 508.5 | 8380.75 | 513.5 | 435.75 | 183.75 | 1520.75 | 780 | 650 |
| 109 | Proteus mirabilis O29a (PrK 52/57) | OPS | 52.75 | 88.5 | 16.5 | 10.5 | -6.25 | 59 | 66.25 | 37.25 | 24.5 | 225 |
| 110 | Proteus mirabilis O40 (10703) | OPS | 41.5 | 117 | 8.75 | 18.25 | 6 | 56 | 46.5 | 17.25 | 24.25 | 229 |
| 111 | Proteus mirabilis O41 (PrK 67/57) | OPS | 1558.75 | 2146.75 | 1017.25 | 1539.5 | 362.25 | 409.5 | 851.25 | 822.75 | 2017.5 | 683.5 |
| 112 | Proteus mirabilis O51 (19011)* | LPS | 85.5 | 223.25 | 104 | 146.25 | 44.5 | 820.75 | 188.25 | 113.25 | 108 | 1565.25 |
| 113 | Proteus mirabilis O74 (10705, OF) | OPS | 498 | 94.25 | 87.75 | 25 | -70.5 | 101.75 | 38.25 | 106 | 145.5 | -31.75 |
| 114 | Proteus mirabilis O75 (10702, OC) | OPS | 31.75 | 120.5 | 21.25 | -1 | -203.5 | 86.5 | 18 | -1 | 23.75 | 55.75 |
| 115 | Proteus mirabilis O77 (3 B-m) | OPS | -1044 | 61.5 | -71 | 11.75 | -105.25 | 35.25 | 88.5 | -1.75 | 23.5 | 27.5 |
| 116 | Proteus penneri O31a (26) | OPS | 400.5 | 223.5 | 16 | 2 | -133.75 | 45.75 | 14.5 | 2.25 | 30.5 | 142.25 |
| 117 | Proteus penneri O52 (15) | OPS | -96.75 | 106.5 | -11.5 | 10.5 | 11.25 | 46.75 | -15.75 | 3.25 | 31.5 | 200.25 |
| 118 | Proteus penneri O58 (12) | OPS | 22.25 | 135 | 3.25 | 6 | 4 | 74.25 | 21.5 | 10.5 | 22.75 | 310.5 |
| 119 | Proteus penneri O59 (9) | OPS | 180.5 | 106.75 | 38.25 | 10.25 | -43.75 | 24 | 86.25 | 14.5 | 18.5 | 148.75 |
| 120 | Proteus penneri O61 (21) | OPS | 27618.75 | 1594.75 | 17.5 | 19.75 | 1.25 | 79 | 64 | 21.75 | 70.5 | 363 |
| 121 | Proteus penneri O62 (41) | OPS | 103.25 | 106.5 | 14.25 | 35.25 | -130.5 | 101 | 86.75 | 51.5 | 102.5 | 150.25 |
| 122 | Proteus penneri O63 (22) | OPS | 755.75 | 270 | 44.25 | 21.25 | 428.25 | 299.75 | 87.25 | 23.25 | 33.5 | 1240.5 |
| 123 | Proteus penneri O64a,b,c (27) | OPS | 118.75 | 86.5 | 16.25 | -2 | -72.25 | 126.75 | 39.75 | 22.5 | 14 | 205 |
| 124 | Proteus penneri O65 (34) | OPS | 513.5 | 283 | 15.75 | 8.75 | -158 | 652.25 | 171.75 | 24408.25 | 43.5 | 43.5 |
| 125 | Proteus penneri O67 (8) | OPS | 1121.75 | 577.25 | 171.5 | 1444.25 | 205.25 | 400.25 | 562.75 | 684 | 678.25 | 600.75 |
| 126 | Proteus penneri O68 (63) | OPS | 332 | 3942.75 | 177.25 | 193.75 | 81.75 | 199.5 | 59.75 | 198.25 | 209.5 | 273 |
| 127 | Proteus penneri O70 (60) | OPS | 1001.5 | 149 | 88 | 82.75 | 2.75 | 653.5 | 378.75 | 193.5 | 146 | 2577.75 |
| 128 | Proteus penneri O73a,b (103) | OPS | 633.25 | 401 | 162.25 | 215.75 | 5.75 | 129.25 | 14429.25 | 1140.5 | 273.25 | 207.5 |

| | | | | | | | | | | | | |
|---|---|---|---|---|---|---|---|---|---|---|---|---|
| 129 | Proteus myxofaciens O60 | OPS | 157 | 152 | 3.5 | 49.75 | 27 | 294.75 | 273.25 | 80.5 | 108.75 | 446.5 |
| 130 | Proteus O56 (genomospecies 4) | OPS | 27280.5 | 6162.25 | 26 | 11.5 | -6 | 47.75 | 25.75 | 14.5 | 58.5 | 296.75 |
| 131 | Providencia stuartii O4 | OPS | 623.25 | 131.75 | -62.25 | 8.75 | 7.25 | 254.75 | 584.75 | 20.5 | 39.75 | 146.75 |
| 132 | Providencia stuartii O18 | OPS | 167.75 | 525.75 | 6.5 | 0.5 | 7.75 | 125.5 | 16.25 | 18.75 | 12.75 | 191.75 |
| 133 | Providencia stuartii O20* | LPS | 996.75 | 333.25 | 160.75 | 142.25 | 110.75 | 316 | 455 | 179.5 | 402.75 | 1844.75 |
| 134 | Providencia stuartii O43 | OPS | 175.5 | 283.5 | 51.75 | 5.5 | 46.75 | 108.25 | 35.5 | 5.75 | 64.75 | 322.75 |
| 135 | Providencia stuartii O44 | OPS | 728.5 | 649.5 | 21.25 | 14.75 | -36.25 | 55 | 18.5 | 45.5 | 39.25 | 193 |
| 136 | Providencia stuartii O47 | OPS | 43.25 | 347.25 | 15.75 | 21.25 | 13.75 | 59 | 7.5 | 22.75 | 23.25 | 345.5 |
| 137 | Providencia stuartii O47, Core 9 | OPS | 153.25 | 264 | 90.5 | 120.75 | 292.75 | 153 | 120 | 55.5 | 173.25 | 718.75 |
| 138 | Providencia stuartii O49, Core 1 | OPS | 235.5 | 274.5 | 110.5 | 277.25 | 526.5 | 239.25 | 253 | 65.25 | 152.75 | 1193.75 |
| 139 | Providencia stuartii O57 | OPS | 86 | 133.75 | 19.25 | 17.5 | -13.25 | 29.25 | 76 | 17.25 | 28.25 | 165.25 |
| 140 | Providencia alcalifaciens O5 | OPS | 109.5 | 140.5 | 21.5 | 41.5 | 37.5 | 59539 | 41969.75 | 48604 | 68 | 760.5 |
| 141 | Providencia alcalifaciens O6* | LPS | 140.75 | 197.25 | 186.25 | 75 | 444.75 | 39700.5 | 37527.75 | 49159.25 | 224.75 | 4217 |
| 142 | Providencia alcalifaciens O19 | OPS | 14.25 | 138.75 | -29.5 | 11.5 | -93.25 | 68 | 167 | 29.25 | 75 | 214.25 |
| 143 | Providencia alcalifaciens O19 | LPS | 165.25 | 333.25 | 217.25 | 95.25 | -82.75 | 327 | 289 | 237.25 | 313.25 | 1617.25 |
| 144 | Providencia alcalifaciens O19 | LPS/NaOH | 78.5 | 475.5 | 155 | 107 | 197 | 776.25 | 585.75 | 86.25 | 123.5 | 1621.5 |
| 145 | Providencia alcalifaciens O21 | OPS | 252.75 | 189.75 | 81 | 95 | 29.75 | 336.5 | 35782.25 | 24204 | 97.5 | -448.5 |
| 146 | Providencia alcalifaciens O23 | OPS | 83.75 | 175.5 | 62.75 | 81.5 | 98.25 | 452.5 | 896 | 10 | 141.75 | 398.75 |
| 147 | Providencia alcalifaciens O27 | OPS | -289.75 | 170.75 | 36.25 | 23 | 12.75 | 55.25 | 334.75 | 11 | 47 | 282.25 |
| 148 | Providencia alcalifaciens O29 | OPS | 73.75 | 742.75 | 18.5 | 18.5 | -124.5 | 59.75 | 40.5 | 2 | 64.25 | 266.75 |
| 149 | Providencia alcalifaciens O30 | OPS | 1220.75 | 313 | 94 | 96.75 | 58.75 | 119 | 264.25 | 55.75 | 141 | 544.5 |
| 150 | Providencia alcalifaciens O32 | OPS | 36 | 555 | 1 | 33.25 | -48 | 81.5 | 82.5 | -2.25 | 76.75 | 11 |
| 151 | Providencia alcalifaciens O36* | LPS-NH4OH | 334.5 | 227 | 187.5 | 633.5 | 53.75 | 372 | 356.25 | 305.5 | 344 | 2341.75 |
| 152 | Providencia alcalifaciens O39 | OPS | 87.75 | 160.75 | 33 | 44.25 | 43.25 | 49.5 | 61 | -1.25 | 107.5 | -28.5 |
| 153 | Providencia rustigianii O14 | OPS | 270.75 | 321 | 213.75 | 111 | 390.5 | 578.75 | 158.25 | 34.5 | 215.5 | 974.75 |
| 154 | Providencia rustigianii O16 | OPS | 35.25 | 29.25 | 51.25 | 42.5 | -150.25 | 146 | 79.75 | 16.5 | -6.5 | 225 |
| 155 | Providencia rustigianii O34 | OPS | 397.5 | 190.5 | 6.5 | 12.25 | -73 | 355 | 80.25 | 8.5 | 53.25 | 68.75 |
| 156 | Yersinia pestis, KM260(11)-Δ0187 | LPS | 55405.25 | 11953.25 | 493.25 | 132 | 17.5 | 345.25 | 286.25 | 72 | 376 | 1501.25 |
| 157 | Yersinia pestis, KM260(11)-Δ0187 | Core oligo saccharide | 261 | 400.5 | 33.75 | 4.25 | 45.5 | 50.5 | 24.25 | -2.75 | 81.75 | 468.5 |
| 158 | Yersinia pestis, KM260(11)-Δrfe | LPS | 26897.75 | 6382.5 | 292.25 | 84.75 | -287.5 | 327.75 | 319 | 63 | 347 | 774.5 |
| 159 | Yersinia pestis, KM260(11)-Δrfe | Core oligo saccharide | 277.75 | 456.25 | 61.5 | -0.25 | 64 | 54.75 | 74 | -1.5 | 203.25 | 700 |
| 160 | Yersinia pestis, 1146-25 | LPS | 42354.25 | 6472.5 | 326 | 107 | -251 | 189.25 | 354.5 | 97.75 | 435.5 | 378.5 |
| 161 | Yersinia pestis 1146-25 | Core oligo saccharide | -64.75 | 305.25 | 32.75 | 14.25 | 93.75 | 99 | 1 | 10.75 | 62 | 705.75 |
| 162 | Yersinia pestis, 1146-37 | LPS | 25548.5 | 3130.5 | 270.25 | 102.25 | 46 | 317.75 | 887 | 131.25 | 497 | 732.5 |
| 163 | Yersinia pestis, 1146-37 | Core oligo saccharide | -145 | 257.5 | 76 | 17.5 | 36.25 | 77.75 | 7.25 | 4.25 | 222.5 | 443.75 |
| 164 | Yersinia pestis, 0KM218-37 | LPS | 5436.5 | 749.75 | 117.75 | 61.5 | -15.25 | 870 | 316.25 | 46.75 | 372.5 | 955 |
| 165 | Yersinia pestis, KM218-37 | Core oligo saccharide | -58.5 | 281.5 | 44.25 | -1 | 125.75 | 52.75 | -18.75 | 4.75 | 73.25 | 423.5 |
| 166 | Yersinia pestis, KM218-25 | LPS | 45772.25 | 11687 | 552.5 | 111 | -70.25 | 204.25 | 497.25 | 114.75 | 929.25 | 1253.5 |
| 167 | Yersinia pestis, KM218-25 | Core oligo saccharide | -129.5 | 283.25 | 50.75 | 5.25 | 51.5 | 68.5 | 9 | 8.5 | 25.25 | 513.25 |
| 168 | Yersinia pestis, KM260(11)-ΔpmrF | LPS | 8537.25 | 1273.25 | 111.25 | 46 | -268.75 | 604.5 | 200.75 | 18 | 251.5 | 1380 |
| 169 | Yersinia pestis, KM260(11)-ΔpmrF | Core oligo saccharide | 52.25 | 437 | 41.75 | 6.5 | -22.25 | 99.75 | 2.5 | 5.25 | 68 | 537.25 |
| 170 | Yersinia pestis, KM260(11)-Δ0186 | LPS | 54380.25 | 14641 | 853.25 | 42.75 | 111.25 | 366.5 | 1076.5 | 88.25 | 517.75 | 1134.5 |
| 171 | Yersinia pestis, KM260(11)-Δ0186 | Core oligo saccharide | 65.75 | 424.75 | 33.75 | -1 | 2.75 | 90 | 22.5 | 0.25 | 95.5 | 463 |
| 172 | Yersinia pestis, KM260(11)-ΔwaaQ | LPS | 8553.25 | 2293.5 | 521.5 | 65.5 | -308 | 1687 | 570 | 57 | 405.25 | 1544 |
| 173 | Yersinia pestis, KM260(11)-ΔwaaQ | Core oligo saccharide | 43.75 | 409.75 | 47 | 18.5 | 81.5 | 114.5 | 30.75 | 1.5 | 86.25 | 564.25 |
| 174 | Yersinia pestis, KM260(11)-ΔwaaL | LPS | 55426.25 | 12181.25 | 409.25 | 53.5 | -186.5 | 465.5 | 790.25 | 44.75 | 492.5 | 1316.75 |
| 175 | Yersinia pestis, KM260(11)-25 | LPS | 31099.25 | 4491.5 | 397.5 | 66.25 | -199.5 | 462.5 | 213 | 98.5 | 433.25 | 1423.25 |
| 176 | Yersinia pestis, KM260(11)-25 | Core oligo saccharide | 238.25 | 254 | 13.75 | 9.25 | 8 | 65.75 | 63 | 8.5 | 88.75 | 373 |
| 177 | Yersinia pestis, KM260(11)-37 | Core oligo saccharide | -95.5 | 308.5 | 47 | 20.75 | 221.75 | 272.25 | 47.75 | 3.75 | 86 | 844.75 |
| 178 | Yersinia pestis, KIMD1-37 | Core oligo saccharide | 311 | 545.75 | 33 | 1.5 | 82.5 | 235.5 | -58.5 | 2.5 | 42.25 | 552 |
| 179 | Yersinia pestis, KIMD1-25 | Core oligo saccharide | 1695 | 305.75 | 22.25 | 12.25 | 252 | 363.5 | 100 | 15.75 | 22 | 719.5 |
| 180 | Yersinia pestis, 11M-25 | LPS | 378.5 | 306.5 | 219 | 64 | 17 | 151 | -98.5 | 77.75 | 302.75 | 301 |
| 181 | Yersinia pestis, 11M-37 | LPS | 184.75 | 467.5 | 193.5 | 278.75 | -48.25 | 107 | 607.25 | 108.25 | 314 | 204 |
| 182 | Proteus mirabilis O23a, 23b, 23c (CCUG 10701) | OPS | 53 | 328.25 | 35.75 | 19.75 | 25.5 | 92.25 | 44.5 | 0.5 | 58 | 418.25 |
| 183 | Proteus vulgaris O24 (PrK 47/57) | LPSOH | -75.75 | 223.5 | 150.25 | 132.75 | 228.75 | 427 | 32595.25 | 1692 | 152 | 387.75 |
| 184 | Yersinia pestis KM260(11)-6C | LPS | 43055.5 | 7088.75 | 458.25 | 55 | 10.5 | 1506 | 269.75 | 110.25 | 477 | 2976.25 |
| 185 | Yersinia pestis 260(11)-37C-186 | LPS | 46574.25 | 2105.5 | 375.25 | 73.5 | 824.5 | 879.75 | 295 | 73 | 456.5 | 1855.5 |
| 186 | Yersinia pestis 260(11)-37C-187 | LPS | 45605.5 | 2782.5 | 295.5 | 38.5 | -155 | 526.5 | 583.5 | 49 | 281.5 | 958.75 |
| 187 | Yersinia pestis 260(11)-37C-416 | LPS | 748.25 | 501.25 | 461.5 | 105.75 | 190 | 1701.75 | 180.5 | 105.75 | 573.25 | 5442.75 |
| 188 | Yersinia pestis 260(11)-37C-417 | LPS | 3229 | 929.5 | 306.5 | 100.25 | -206 | 1322.5 | 368.75 | 75.25 | 261.25 | 697.25 |
| 189 | Yersinia pestis P-1680-25C | OS | 211.5 | 280 | 45 | 19.5 | 84.5 | 98.5 | 103.75 | 7.25 | 58.75 | 445 |

| | | | | | | | | | | | | |
|---|---|---|---|---|---|---|---|---|---|---|---|---|
| 190 | Yersinia pestis P-1680-37C | LPS | 37028.5 | 2290.25 | 373.25 | 73.75 | -252 | 464.25 | -272.5 | 94.75 | 301.5 | 295.5 |
| 191 | Yersinia pestis I-2377-25C | OS | 113.75 | 394.25 | 32.5 | 6.25 | 72.75 | 59.75 | 36.75 | 5 | 60.75 | 475 |
| 192 | Yersinia pestis I-2377-37C | LPS | 5870.75 | 1167.75 | 181.5 | 49.25 | -114.75 | 734.75 | 634 | 59.75 | 204 | 738 |
| 193 | Francisella novicida OPS | OPS | 111.25 | 362 | 18.75 | 5 | -77.75 | 19.75 | -151 | 4.75 | 132 | 184 |
| 194 | Francisella tularensis OPS | OPS | 62.25 | 360 | 8 | -7.75 | -228.25 | -2 | -36.5 | -1 | 16.25 | -56.75 |
| 195 | Klebsiella O1 OPS | OPS | 26.25 | 160.5 | 12.5 | 14.25 | -133.5 | 1516.5 | 27971.75 | 151.75 | 30.25 | 192.5 |
| 196 | Klebsiella O2a OPS | OPS | 3651 | 1715.5 | 56.25 | 13 | -156.25 | 2285.75 | -2972 | 39863.5 | 64.75 | 1438 |
| 197 | Klebsiella O2ac OPS | OPS | 558.75 | 562.5 | 25.25 | -2.25 | -440 | 31.75 | 8746 | 1 | 153.75 | 320.25 |
| 198 | Klebsiella O3 OPS | OPS | 17116.25 | 6164.5 | 460.25 | 9.5 | -79.5 | 1468.5 | -67.25 | 305.5 | 26 | 257.25 |
| 199 | Klebsiella O4 OPS | OPS | 387.5 | 178 | 16.5 | 13.5 | -387.5 | 1402.25 | 68.25 | 300.25 | 18 | 348 |
| 200 | Klebsiella O5 OPS | OPS | 44733 | 9633.75 | 17.5 | 4.25 | -128.5 | 38.75 | 660.5 | 15.75 | 51.75 | 976 |
| 201 | Klebsiella O8 OPS | OPS | 623.5 | 224.5 | 29 | 76 | -271.75 | 834.5 | 48170.75 | 340.5 | 21 | 83.25 |
| 202 | Klebsiella O12 OPS | OPS | 350 | 182.5 | 17 | 1 | -174.5 | 175.75 | 732 | -3.5 | 35.25 | 1024.25 |
| 203 | Shigella boydii type 1 | LPSOH | 792 | 407 | 38.25 | 15.75 | 164.25 | 1124.75 | 74.25 | 344.25 | 58 | 2512.5 |
| 204 | Shigella boydii type 3 | OPS | 448.25 | 191 | 6 | 84.75 | -45.75 | 1730.75 | -121.25 | 1433.75 | 6 | 150.25 |
| 205 | Shigella boydii type 5 | OPS | 73.25 | 162.75 | 8.5 | 11.75 | -110 | 20493.75 | 31 | 35386.5 | 5.75 | 204.25 |
| 206 | Shigella boydii type 9 | OPS | 378.25 | 97 | 35.5 | 8.5 | -181.25 | 829.5 | 1848 | 17.75 | 188 | 459.5 |
| 207 | Shigella boydii type 11 | OPS | 1461.25 | 632.75 | 6.5 | -3 | -260.75 | 118.5 | 3.75 | 16.75 | 13.5 | 404.5 |
| 208 | Shigella boydii type 12 | OPS | 17492.25 | 3651 | 13.25 | -7.5 | -138.5 | 271.25 | 109 | 5.5 | 49.25 | 61.75 |
| 209 | Shigella boydii type 15 | OPS | 1312.25 | 506.5 | 4.75 | 2.5 | -110.75 | 45.75 | 78.25 | 9.75 | 14.5 | 250 |
| 210 | Shigella boydii type 16 | OPS | 863 | 306 | 14.25 | 16 | 130.5 | 913 | 25 | 6.75 | 25.75 | 1169.75 |
| 211 | Shigella boydii type 17 | OPS | 160.25 | 183.75 | 11.25 | 13.25 | 197.25 | 100.5 | 779.75 | 146.75 | 15 | 219 |
| 212 | Shigella boydii type 18 | OPS | 970 | 310.75 | 11.75 | -1.75 | 361.5 | 380.5 | 22.75 | 3.25 | 13 | 822.75 |
| 213 | Escherichia coli O49 | OPS | 11058.75 | 2738.5 | 8.75 | 2.25 | -164.25 | 257 | 179 | 14.25 | 23 | 305.5 |
| 214 | Escherichia coli O52 | OPS | 293.25 | 1658.75 | 17.5 | -2 | -29.75 | 160.5 | 177 | 96.75 | 16.75 | 183.25 |
| 215 | Escherichia coli O58 | OPS | 1189.25 | 1579.25 | 9.25 | 7.5 | -250 | 109.25 | 88.25 | -17.25 | 50.25 | 302.5 |
| 216 | Escherichia coli O61 | LPSOH | 315 | 245.25 | 41 | 16.25 | -37.5 | 641.75 | 150.75 | 22.5 | 51.5 | 2822 |
| 217 | Escherichia coli O73 | OPS | 1883.75 | 3699 | 9.75 | 32.25 | -74 | 27 | 6.75 | 8.75 | 6 | 337 |
| 218 | Escherichia coli O112ab | OPS | 5549 | 1494 | 23.25 | 7.25 | -157.75 | 570.75 | 93.5 | 80.75 | 28 | 388.5 |
| 219 | Escherichia coli O118 | OPS | 84.5 | 163 | 11.25 | 2.25 | -198.75 | 101.25 | 67.5 | 11.5 | 12.75 | 76 |
| 220 | Escherichia coli O125 | OPS | 330.5 | 237.5 | 10.25 | -4.5 | -138.25 | 127.25 | 983.5 | 3046.25 | 12 | 79 |
| 221 | Escherichia coli O151 | OPS | 1095 | 305 | 28.5 | 16.25 | -72.75 | 346.75 | 93 | 47.25 | 23 | 193.5 |
| 222 | Escherichia coli O168 | OPS | 906.75 | 229.75 | 18.5 | 6.75 | -17 | 500.75 | 6.5 | 46.5 | 25.5 | 416 |
| 223 | Shigella dysenteriae type 2 | LPSOH | 409 | 220.75 | 40.5 | 19.75 | 353.5 | 630.75 | 83.5 | 71.5 | 38.75 | 2630 |
| 224 | Shigella dysenteriae type 4 | OPS | 803.5 | 367 | 3.5 | -1 | -94.75 | 51.75 | 19.75 | 33 | 14.25 | 131 |
| 225 | Shigella dysenteriae type 5 | OPS | 4165.75 | 259.5 | 34 | 128.75 | -184 | 158.75 | 6.25 | 14.25 | 29.25 | 484.5 |
| 226 | Shigella dysenteriae type 6 SR-strain | SR-strain | -18 | 322.75 | 11.75 | 36 | 23 | 45.25 | 4.25 | 20 | 14.5 | 411.25 |
| 227 | Shigella dysenteriae type 7 | OPS | 1829.25 | 396 | 2 | 108.75 | -256.5 | 76.5 | 31.75 | 29.75 | 19.5 | 130.75 |
| 228 | Shigella dysenteriae type 8 (Russian) | OPS | 53.25 | 2312 | 29 | 59 | -59.5 | 76.75 | 10.75 | 34.5 | 23.25 | 155 |
| 229 | Shigella dysenteriae type 9 | OPS | 622 | 160.5 | 9 | 155 | -170 | 29.5 | -22.75 | 24.5 | 14.75 | 387.75 |
| 231 | Escherichia coli O111:B4 LPS- solution at 1 mg/mL | L5293-2ML (LPS) (Sigma) | -12 | 12.5 | 9.25 | 110 | 327.75 | 166.75 | 3.25 | 71 | 10.5 | 300.25 |
| 232 | Escherichia coli O26:B6 LPS- solution at 1 mg/mL | L5543-2ML (LPS) (Sigma) | -13 | 13.75 | 6.75 | 29.25 | 294 | 99.75 | 6.25 | 88 | 10 | 34.5 |
| 233 | Escherichia coli O55:B5 LPS- solution at 1 mg/mL | L5418-2ML (LPS) (Sigma) | 69 | 16.5 | 18 | 79 | 80.25 | 13362 | 23033.5 | 51975 | 18 | 165.75 |
| 234 | Escherichia coli O127:B8 LPS- solution at 1 mg/mL | L5668-2ML (LPS) (Sigma) | -63 | -2.5 | -6.75 | 423.75 | 317.25 | 146.5 | 1605.25 | 13389.25 | 0 | 187.5 |
| 235 | Streptococcus pneumoniae type 1 (Danish type 1) | 161-X // Capsular PS | 64 | 31 | 37 | 45035 | 9.5 | 279 | 2668 | 140 | 44 | 907.75 |
| 236 | Streptococcus pneumoniae type 2 (Danish type 2) | 165-X// Capsular PS | 25.75 | 2 | 0.75 | 31008.5 | -308.25 | 32.5 | -43.75 | 20.25 | -4.75 | 226 |
| 237 | Streptococcus pneumoniae type 3 (Danish type 3) | 169-X// Capsular PS | 93.25 | 228 | 21.75 | 319.25 | -265.75 | 59 | 468.25 | 27 | 27.75 | 1146 |
| 238 | Streptococcus pneumoniae type 4 (Danish type 4) | 173-X// Capsular PS | -41.25 | -1.25 | -2 | 38763.25 | 241.25 | 154.75 | -9.75 | 20.5 | 7.25 | 932.5 |
| 239 | Streptococcus pneumoniae type 5 (Danish type 5) | 177-X// Capsular PS | 16.5 | 106.5 | 8 | 46500.75 | -0.5 | 103.75 | 120.25 | 179.25 | 15.25 | 482 |
| 240 | Streptococcus pneumoniae type 8 (Danish type 8) | 185-X// Capsular PS | -11.25 | 3.75 | 97 | 26651.5 | 69.5 | 391.25 | -95.25 | 17.25 | -7 | 1179 |
| 241 | Streptococcus pneumoniae type 9 (Danish type 9N) | 189-X// Capsular PS | -17 | 37 | -8.5 | 35161 | -158.5 | 36.75 | -10.75 | 17.25 | 22.5 | 170.25 |
| 242 | Streptococcus pneumoniae type 12 (Danish type 12F) | 193-X// Capsular PS | -44.25 | 48 | 16.25 | 43233.75 | -143.75 | 42.5 | 14.25 | 28.5 | 24.25 | -200 |
| 243 | Streptococcus pneumoniae type 14 (Danish type 14) | 197-X// Capsular PS | 87.75 | 70.25 | -9.25 | 7803.25 | 136.75 | 39481 | 6039.75 | 49829.5 | 23.5 | 872.5 |
| 244 | Streptococcus pneumoniae type 17 (Danish type 17F) | 201-X// Capsular PS | 6.75 | 32.75 | 6.75 | 57173.5 | 136.75 | 114.75 | -17 | 38 | 9.25 | 309.75 |
| 245 | Streptococcus pneumoniae type 19 (Danish type 19F) | 205-X// Capsular PS | -22.75 | 51 | 3.5 | 42576.75 | -80.5 | 41.25 | 118 | 258.75 | 21 | 268 |
| 246 | Streptococcus pneumoniae type 20 (Danish type 20) | 209-X// Capsular PS | 9.25 | 77.5 | 19 | 51050.5 | -77.75 | 52.25 | 440.75 | 122 | 14.5 | -13.5 |
| 247 | Streptococcus pneumoniae type 22 (Danish type 22F) | 213-X// Capsular PS | -19.5 | 33.75 | 20.25 | 40434.75 | -267.25 | 44.5 | 124.5 | 94.25 | 25.5 | 123.5 |
| 248 | Streptococcus pneumoniae type 23 (Danish type 23F) | 217-X// Capsular PS | 14.75 | 36.5 | 11 | 5341 | -292 | 20.25 | 32.25 | 5.5 | 9.25 | -292 |
| 249 | Streptococcus pneumoniae type 26 (Danish type 6B) | 225-X// Capsular PS | 25.5 | 46.75 | 10.25 | 48596.25 | 109 | 188 | 162.25 | 48.25 | 8 | 158.75 |
| 250 | Streptococcus pneumoniae type 34 (Danish type 10A) | 229-X// Capsular PS | 327 | 312.25 | 148 | 55028.25 | -75.75 | 61 | 950.5 | 1492.75 | 631 | 357.5 |
| 251 | Streptococcus pneumoniae type 43 (Danish type 11A) | 233-X// Capsular PS | 26 | 54.25 | 16 | 47263 | -97.25 | 52363.75 | 30264.25 | 52209.5 | 14.75 | 142.25 |

| | | | | | | | | | | | | |
|---|---|---|---|---|---|---|---|---|---|---|---|---|
| 252 | Streptococcus pneumoniae type 51 (Danish type 7F) | 237-X// Capsular PS | 127 | 66.25 | 14 | 54120.75 | -156.25 | 56 | -7.75 | 155.5 | 26 | 171.5 |
| 253 | Streptococcus pneumoniae type 54 (Danish type 15B) | 241-X// Capsular PS | 23.5 | -2.75 | 9.75 | 44079 | -232.75 | 40412.25 | 380.25 | 946 | 17.75 | -94.75 |
| 254 | Streptococcus pneumoniae type 56 (Danish type 18C) | 245-X// Capsular PS | -7.75 | 32.75 | 0.5 | 58876 | -338 | 21.25 | -49.25 | 11.5 | 24.5 | 117.5 |
| 255 | Streptococcus pneumoniae type 57 (Danish type 19A) | 249-X// Capsular PS | 38.25 | 263.25 | 14.5 | 54796.25 | 134.25 | 112.25 | 280.25 | 928.25 | 22.5 | 646.75 |
| 256 | Streptococcus pneumoniae type 68 (Danish type 9V) | 253-X// Capsular PS | 23.75 | 68.5 | 3 | 61800.25 | 22.5 | 141.5 | 69.5 | 152.75 | 15.25 | 512 |
| 257 | Streptococcus pneumoniae type 70 (Danish type 33F) | 257-X// Capsular PS | 505.25 | 204.5 | -16.5 | 38404.75 | -187.25 | 8366.75 | 50221.75 | 58416.25 | 16.25 | 73.25 |
| 258 | Yersinia pestis KM218-6C | OS | 138.5 | 414.5 | 42.75 | 18.5 | -70.25 | 304 | -36 | 26.25 | 77 | 1239 |
| 259 | Yersinia pestis KM260(11)-yjhW-6C | OS | 172 | 160.25 | 214.25 | 36.5 | 24.25 | 117.5 | 8081.75 | 1305.5 | 34.25 | 442.25 |
| 260 | Yersinia pestis KM260(11)-wabD/waaL | OS | 58.75 | 280.25 | 15.25 | 12.25 | 14 | 53.75 | 54.25 | 18.25 | 87.75 | 427.75 |
| 261 | Yersinia pestis KM260(11)-wabC/waaL | OS | 44049.75 | 9042 | 290.75 | 47 | -471.25 | 796 | 1238.5 | 226 | 392.5 | 1877.75 |
| 262 | Yersinia pseudotuberculosis 85pCad-37C | OS | -113.75 | 236.75 | 15.25 | 15.5 | 3.5 | 101.25 | 44.75 | 19.5 | 27.25 | 372.75 |
| 263 | Yersinia pseudotuberculosis 85pCad-20C | OS | 86.75 | 111.5 | 36 | 18.5 | -70 | 132 | 191 | 40.5 | 10.25 | 553.25 |
| 264 | Yersinia pseudotuberculosis O:2a | PS | 18565 | 7559.5 | 52.75 | 13 | 3 | 123.75 | 50.5 | 25.75 | 70.75 | 407 |
| 265 | Yersinia pseudotuberculosis O:2a-dhmA | PS | 1372.75 | 1821 | 24.5 | -10.5 | -73.25 | 59.5 | 45 | 5.75 | 31.25 | 266.5 |
| 266 | Yersinia pseudotuberculosis O:2c | PS | 2191.5 | 849.5 | 265.5 | 37.25 | -89.75 | 576 | 621.25 | 44.75 | 56.25 | 1314 |
| 267 | Yersinia pseudotuberculosis O:3 | PS | 1872.25 | 656 | 9 | -1 | -175.25 | 69.25 | 17.5 | 2.25 | 10.25 | 192.75 |
| 268 | Yersinia pseudotuberculosis O:4b | PS | 3743 | 2736 | 18.25 | 4 | -130 | 98 | 102.75 | 23 | 20.75 | 168.75 |
| 269 | Proteus vulgaris O2 (OX2) | PS | 3261 | 106.25 | 11.5 | 4.25 | -78.25 | 460.25 | 163.75 | 104.25 | 15.5 | 149 |
| 270 | Proteus mirabilis O3ab (S1959) | PS | 491.75 | 766 | 377.25 | 1222.25 | 610.25 | 279.5 | 239 | 1190 | 435.75 | 316 |
| 271 | Proteus mirabilis O5 (PrK 12/57) | PS | 746.5 | 302.5 | 28.25 | 31.25 | 30.25 | 309.5 | 562 | 71.5 | 37 | 721.25 |
| 272 | Proteus mirabilis O9 (PrK 18/57) | PS | 163.5 | 160.5 | 1.5 | 161.5 | -46.75 | 104.25 | 30.25 | 18.25 | 51 | 180.5 |
| 273 | Proteus mirabilis O11 (9B-m) | PS | 5685.25 | 136.25 | 61.5 | 91 | 47.25 | 88.75 | 69 | 29.75 | 52.25 | 436.5 |
| 274 | Proteus penneri O17 (16) | PS | 180 | 908.5 | 15.5 | 18 | 94 | 83.25 | 16 | 6.75 | 29.25 | 209.5 |
| 275 | Proteus mirabilis O18 (PrK 34/57) | LPSOH | 413 | 197.25 | 211.75 | 35866.75 | 79 | 349.25 | 15965 | 363.75 | 182.75 | 1087 |
| 276 | Proteus mirabilis O20 (PrK 38/57) | LPSOH | 372.75 | 256.5 | 113 | 113.25 | 379.25 | 4560.5 | 893.5 | 7422 | 209 | 3506.5 |
| 277 | Proteus penneri O31ab (28) | PS | 748.5 | 151.25 | 48.25 | 276.25 | 49.5 | 212.25 | 32.5 | 40 | 16.5 | 664.75 |
| 278 | Proteus mirabilis O33 (D52) | PS | 305.25 | 397 | 97 | 197.25 | -31.5 | 4013 | 20231.75 | 2654.25 | 258.25 | 2785 |
| 279 | Proteus mirabilis O43 (PrK 69/57) | PS | 1008.75 | 303.75 | 29.75 | 39.5 | 21.5 | 68.5 | 3.5 | 18.5 | 39.25 | 515.25 |
| 280 | Proteus vulgaris O47 (PrK 73/57) | Not stated | 265.25 | 205 | 8.75 | -2.5 | -110.75 | 23347.25 | 25640.25 | 3945.75 | 50.25 | 218.25 |
| 281 | Proteus mirabilis O49 (PrK 75/57) | PS | 1048 | 325.75 | 17.25 | 44 | 65.75 | 320.25 | 120 | 23.25 | 49 | 234.75 |
| 282 | Proteus mirabilis O54ab (OE) | PS | 115.25 | 118 | 37.25 | 140.25 | -95 | 89.25 | 161.5 | 12.75 | 59.75 | 86.25 |
| 283 | Proteus penneri O73ac (75) | PS | 326.25 | 232.5 | 28.5 | 11.75 | 459.75 | 190.25 | 23112.75 | 697.5 | 34 | 849.75 |
| 284 | Proteus vulgaris O76 (HSC438) | PS | -8.75 | 67 | 13 | 112.75 | -33.75 | 30.5 | -25.25 | 4.5 | 38 | -168.25 |
| 285 | Shigella flexneri type 1a | PS | 1625.25 | 5807.75 | 12.5 | 3.25 | -198.75 | 36 | 20.25 | 3.5 | 18 | 121.25 |
| 286 | Shigella flexneri type 1b | PS | 14461.25 | 7266.5 | 29.75 | 38.5 | -137.5 | 34 | 78 | 27.75 | 48.25 | 140.75 |
| 287 | Shigella flexneri type 2a | PS | 90.5 | 328.5 | 11.75 | -3 | -198.5 | 149.75 | -4 | 23 | 20 | 76.25 |
| 288 | Shigella flexneri type 2b | PS | 18544 | 11040.5 | 12.5 | 27.5 | -248.5 | 3 | 90.75 | 6.5 | 33.5 | 119.75 |
| 289 | Shigella flexneri type 3a | PS | 348.75 | 17373.75 | 26.5 | 20.25 | -249 | 69.75 | 66 | 9.25 | 35.75 | -211.75 |
| 290 | Shigella flexneri type 3b | PS | 9027.25 | 2844.5 | 20 | 14 | 235 | 186.5 | 39.75 | 49.75 | 51.75 | 1207 |
| 291 | Shigella flexneri type 4a | PS | 11483.5 | 4973 | 997.5 | 1571 | 548.75 | 952.25 | 1028.75 | 1377.5 | 1675.75 | 395 |
| 292 | Shigella flexneri type 4b | PS | 7905.5 | 2733.25 | 3.5 | 0.5 | -190.5 | 17.5 | 25.75 | 32.5 | 17 | 106.25 |
| 293 | Shigella flexneri type 5b | PS | 6629.75 | 9145.5 | -51.5 | 6.5 | -152.75 | 15.75 | 100.25 | 22.5 | 27 | 265.25 |
| 294 | Shigella flexneri type 6a | PS | 18830.5 | 4389.5 | 17 | 9.75 | -96.5 | 131.75 | 72.25 | 14.75 | 32.25 | 240 |
| 295 | Shigella flexneri type 6 | PS | 3216.25 | 902.5 | -36.25 | 16.75 | -74 | 96.5 | 202.75 | 10.25 | 29.25 | 348.25 |
| 296 | Shigella flexneri type X | PS | 1716 | 13375.25 | 0.5 | 15.5 | -296.5 | 12 | 53.5 | 28.25 | 28.75 | 21.75 |
| 297 | Shigella dysenteriae type 1 | PS | 1044.5 | 153.25 | 7 | 0.75 | -213.5 | 43.5 | 67.5 | 11.5 | 38 | 7.25 |
| 298 | Shigella boydii type 6 | PS | 185.5 | 121.25 | 15.25 | 1.25 | 104.5 | 126.75 | 27.5 | 8.25 | 23.5 | 294.75 |
| 299 | Shigella boydii type 7 | PS | 6202.75 | 1801.25 | 60 | 46 | 108 | 534.5 | 76 | 51 | 77.5 | 962.75 |
| 300 | Shigella boydii type 8 | PS | 635.25 | 227.25 | 10.75 | 31.25 | -179.75 | 65.5 | -37 | 33.5 | 16.5 | 24 |
| 301 | Shigella boydii type 13 | LPSOH | 7615.25 | 2069.75 | 71.25 | 51.75 | 351.25 | 944.25 | 216.5 | 100.75 | 91 | 3239.25 |
| 302 | Shigella boydii type 14 | LPSOH | 233 | 1018 | 103.75 | 48.75 | 381.25 | 537.75 | 121.25 | 82.75 | 101 | 1115 |
| 303 | Escherichia coli O71 | PS | 6499 | 1441.5 | 10.25 | 5 | -213.25 | 12.25 | 33 | 12 | 11.75 | 41.25 |
| 304 | Escherichia coli O85 | PS | 4916.25 | 1307.25 | 9.25 | 37.25 | -210 | 96 | 12.25 | 38.75 | 18 | 125 |
| 305 | Escherichia coli O99 | PS | 27410.75 | 6894.5 | 29.25 | 2.5 | -55 | 55 | 6.75 | 0.75 | 32.5 | 85.75 |
| 306 | Escherichia coli O145 | LPSOH | 527.25 | 163.25 | 93.75 | 50.25 | 166.75 | 1125.5 | 172 | 105 | 180.25 | 2268.25 |
| 307 | Escherichia coli O107 | PS | 31505.5 | 5181 | -20.25 | 12.25 | 93.5 | 180.75 | 93.25 | 64.75 | 65 | 1277.25 |
| 308 | Salmonella enterica O17 | PS | 1040.5 | 233.75 | 10 | 21 | -149 | 29.75 | 13.5 | 15 | 39.5 | -22.75 |
| 309 | Salmonella enterica O28 | PS | 11644.25 | 1900.5 | 25.25 | 11.25 | -188.25 | -2.5 | 83.25 | 17.25 | 21 | -127.25 |
| 310 | Salmonella enterica O47 | PS | 109.75 | 113.75 | -2 | 16.5 | 45 | 280.25 | 23.75 | 11.75 | 21.75 | 217.75 |
| 311 | Salmonella enterica O55 | PS | 650.5 | 128.5 | 33.25 | 10.75 | -162 | 49.25 | 28.75 | 2.25 | 92.25 | -84.25 |
| 312 | Escherichia coli K92 | CPS | 12.25 | 170.5 | 16.25 | 2 | -66.75 | 32.5 | 8.5 | 5.75 | 13.75 | 207.5 |

| | | | | | | | | | | | | |
|---|---|---|---|---|---|---|---|---|---|---|---|---|
| 313 | Escherichia coli K5 | CPS | 156.25 | 266.25 | 63.25 | 11.5 | -125.75 | 51 | 6.5 | 14.5 | 70.25 | 994.75 |
| 314 | Escherichia coli K13 | CPS | 15.5 | 589.5 | 37.25 | 32.75 | 80.5 | 153.5 | 27.25 | 36.75 | 40 | 769.25 |
| 315 | Neisseria meningitidis Group C | CPS | 35.25 | 196.25 | -4.5 | 9.25 | -81.5 | 71.5 | 114 | 13.75 | 29 | 983.75 |
| 316 | Davanat | | 90 | 205.25 | -0.25 | 15.5 | -222.25 | 35.5 | -36.25 | 12.25 | 34 | 254.5 |
| 317 | Laminarin | | 721 | 266.75 | 375.25 | 17 | 6 | 30 | 2884 | 104.75 | 510.5 | 548.25 |
| 318 | Yeast Mannan | | 31739.25 | 14298.75 | 273.5 | 204.5 | 45.25 | 68.25 | 131.5 | 30.25 | 328.75 | 465.75 |
| 319 | Escherichia coli O86 | | 122.75 | 115.75 | 18.5 | 14.25 | -124 | 442.75 | 35571 | 47963.5 | 59.25 | 219.5 |
| 320 | Galactomannan DAVANT (160102) Pro-Pharmacenti | | 111.75 | 154 | 49 | 22 | -131 | 20.75 | -2.25 | 11 | 38.75 | 319.75 |
| 321 | Yeast Mannan Sigma M-3640 | | 21222 | 14066.5 | 251.25 | 180 | 91.75 | 101.25 | 14.75 | 11.25 | 540.75 | 684 |
| 322 | 1-2 Mannan Acetobacter methanolieus MB135 | | 880.75 | 559.75 | 167 | 20.75 | 12.75 | 67 | 2283.5 | 85.25 | 803.25 | 447.75 |

### SI-Spreadsheet 3

| Chart# | BPS# | BACTERIA / STRAIN | NAME / STRUCTURE /Cat.No. | STRUCTURE* |
|---|---|---|---|---|
| 1 | 1 | Providencia stuartii O49 | PO49 Core-linked | 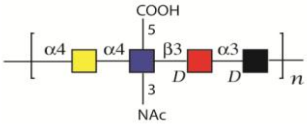 |
| 2 | 2 | Providencia stuartii O52 | PO52 Core-linked | 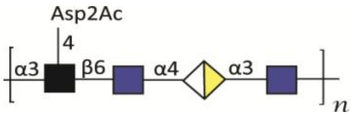 |
| 3 | 3 | Pseudomonas aerguinosa O4 (Habs serotype 4) | PO4 Core-linked | 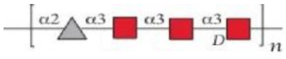 |
| 4 | 4 | Pseudomonas aerguinosa O1 (Fisher immunotype 4) | PO1 Core-linked | 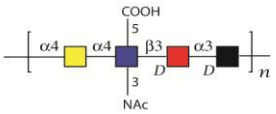 |
| 5 | 5 | Pseudomonas aerguinosa O2 (Fisher immunotype 3) | PO2 Core-linked | 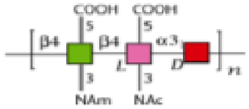 |
| 6 | 6 | Pseudomonas aerguinosa O13 (Sandvik serotype II) | PO13 Core-linked | 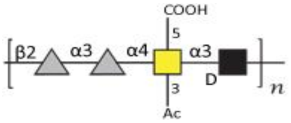 |
| 7 | 7 | Pseudomonas aerguinosa O9 (9a, 9b, 9d) | PO9 Core-linked | 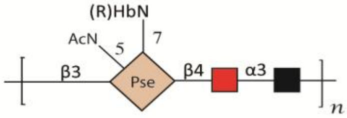 |
| 8 | 8 | Pseudomonas aerguinosa O6a (Habs serotype6, fraction IIa) | PO6a Core-linked-O-unit | 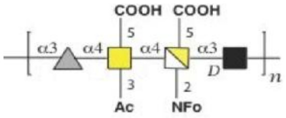 |
| 9 | 9 | Pseudomonas aerguinosa O6a (Habs serotype6, fraction IIb) | PO6a unsubstituted core | 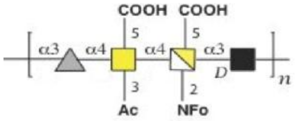 |
| 10 | 12 | Salmonella typhimurium SL 11881 (Re mut) | LPS-L9516 | 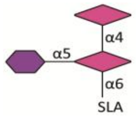 |
| 11 | 13 | Salmonella typhimurium TV 119 (Ra mut) | LPS-L6016 | 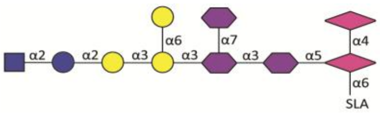 |
| 12 | 14 | Salmonella typhimurium SL 684 (Rc mut) | LPS-L5891 | 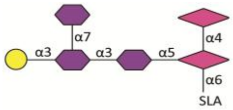 |

| | | | | |
|---|---|---|---|---|
| 13 | 15 | Pseudomonas aerguinosa O10 | L8643 | 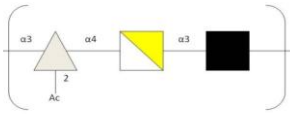 |
| 14 | 16 | Salmonella typhimurium dodeca saccharide | 4809 | 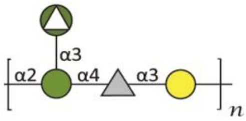 |
| 15 | 17 | Salmonella enteritidis dodeca saccharide | 1262 | 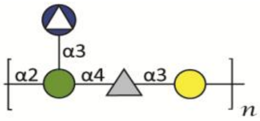 |
| 16 | 18 | Salmonella typhimurium LPS | L2262 | 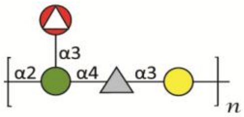 |
| 17 | 20 | Serratia marcescens LPS | L6136 | 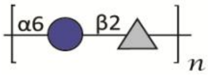 |
| 18 | 22 | Escherichia coli K235 LPS | L2143 | 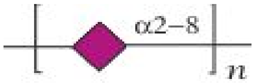 |
| 19 | 23 | Escherichia coli O128-B12 LPS | L2755 | 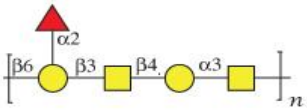 |
| 20 | 25 | Salmonella enterica abortus equi LPS | L5886 | 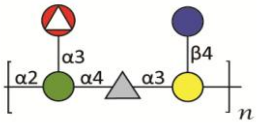 |
| 21 | 26 | Salmonella typhosa LPS | L2387 | 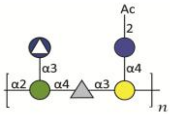 |
| 22 | 27 | Salmonella enteritidis LPS | L2012 | 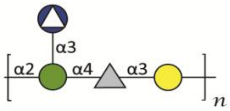 |
| 23 | 28 | Shigella bodyii type2 | | 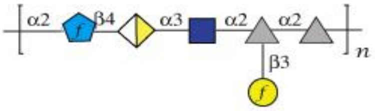 |
| 24 | 29 | Shigella bodyii type4 | | 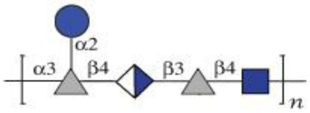 |
| 25 | 30 | Shigella bodyii type10 | | 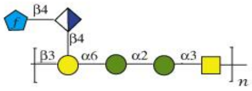 |

| | | | | |
|---|---|---|---|---|
| 26 | 31 | Shigella dysennteriae type 3 | | 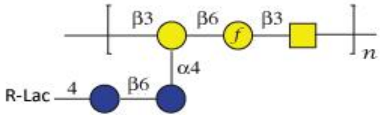 |
| 27 | 32 | Shigella dysennteriae type 8 (batch 12) | | 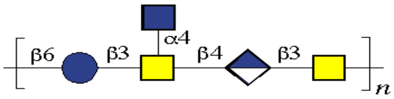 |
| 28 | 33 | Shigella dysennteriae type 11 | | 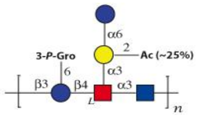 |
| 29 | 34 | Shigella dysennteriae type 13 | | 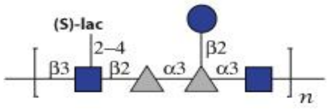 |
| 30 | 35 | Escherichia coli O29 | | 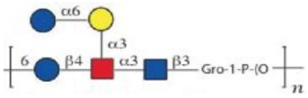 |
| 31 | 36 | Escherichia coli O40 | | 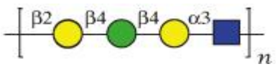 |
| 32 | 37 | Escherichia coli O106 | | 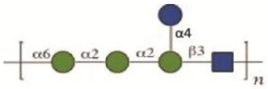 |
| 33 | 38 | Escherichia coli O130 | | 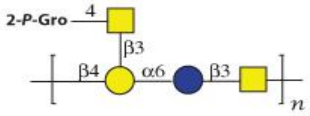 |
| 34 | 39 | Escherichia coli O148 | | 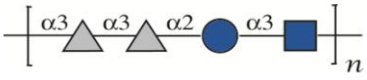 |
| 35 | 40 | Escherichia coli O150 | | 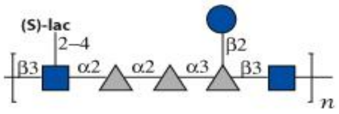 |
| 36 | 41 | Escherichia coli O180 | | 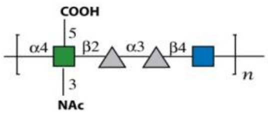 |
| 37 | 42 | Proteus mirabilis O3a, 3c (G1) | | 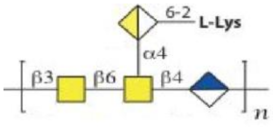 |
| 38 | 43 | Proteus mirabilis O8 (TG326) | | 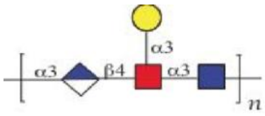 |

| | | | | |
|---|---|---|---|---|
| 39 | 44 | Proteus mirabilis O10 (HJ4320) | | 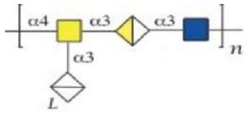 |
| 40 | 45 | Proteus mirabilis O29a, 29b (2002) | | 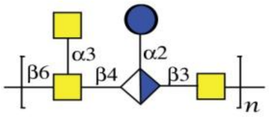 |
| 41 | 46 | Proteus mirabilis O50 (TG332) | | 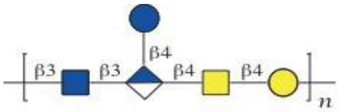 |
| 42 | 47 | Proteus mirabilis O54a, 54b (10704) | | 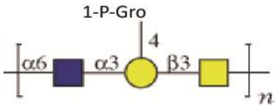 |
| 43 | 48 | Proteus mirabilis O57 (TG319) | | 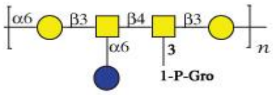 |
| 44 | 49 | Proteus penneri O8 (106) | | 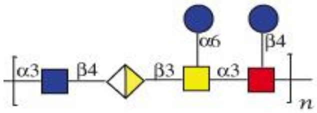 |
| 45 | 50 | Proteus penneri O64a, 64b, 64d (39) | | 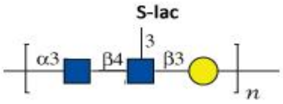 |
| 46 | 51 | Proteus penneri O66 (2) | | 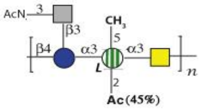 |
| 47 | 52 | Proteus penneri O69 (25) | | 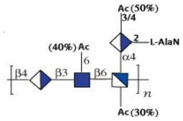 |
| 48 | 53 | Proteus penneri O71 (42) | | 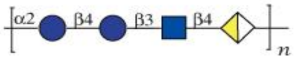 |
| 49 | 54 | Proteus penneri O72a, 72b (4) | | 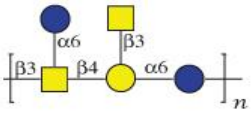 |
| 50 | 55 | Pseudomonas aerguinosa O2 (2a),2d,2f | IATS 10 , OPS | 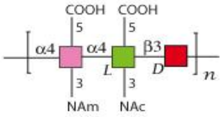 |
| 51 | 56 | Pseudomonas aerguinosa O2 2a,2b | IATS 16 OPS | 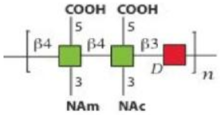 |

| | | | | |
|---|---|---|---|---|
| 52 | 57 | Pseudomonas aerguinosa O2 2a,2b,2e | IATS NO, OPS | 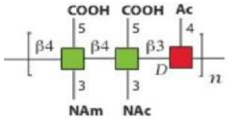 |
| 53 | 58 | Pseudomonas aerguinosa O2 2a,2d | IATS 5 OPS | 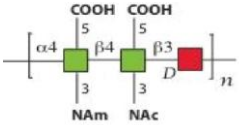 |
| 54 | 59 | Pseudomonas aerguinosa O2 Immuno 7 | IATS 18, OPS | 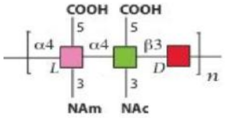 |
| 55 | 60 | Pseudomonas aerguinosa O3 3a,3b | IATS NO, OPS | 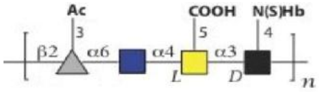 |
| 56 | 61 | Pseudomonas aerguinosa O3 3a,3b,3c | IATS 3, OPS | 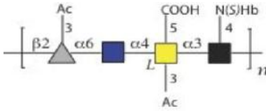 |
| 57 | 62 | Pseudomonas aerguinosa O3 3a,3d | IATS NO, OPS | 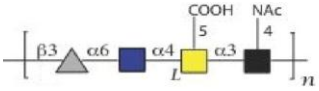 |
| 58 | 63 | Pseudomonas aerguinosa O4 4a,4c | IATS NO, OPS | 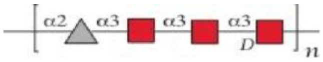 |
| 59 | 64 | Pseudomonas aerguinosa O6 6a | IATS 6, OPS | 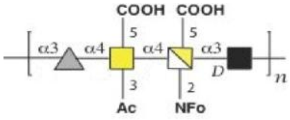 |
| 60 | 65 | Pseudomonas aerguinosa O6 6a,6c | IATS NO, OPS | 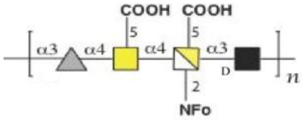 |
| 61 | 66 | Pseudomonas aerguinosa O6 Immuno 1 | IATS NO, OPS | 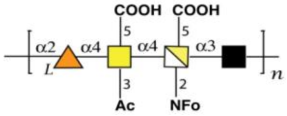 |
| 62 | 67 | Pseudomonas aerguinosa O7 7a,7b,7c | IATS 7,LPS | 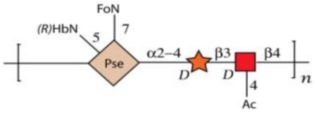 |
| 63 | 68 | Pseudomonas aerguinosa O7 7a,7b,7d | IATS 8,LPS | 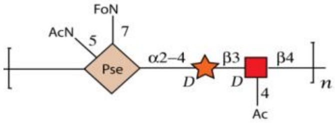 |
| 64 | 69 | Pseudomonas aerguinosa O7 7a,7d | IATS NO, LPS | 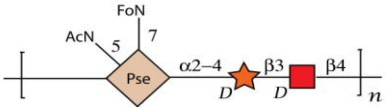 |

| | | | | |
|---|---|---|---|---|
| 65 | 71 | Pseudomonas aerguinosa O10 10a,10b | IATS 10, OPS | 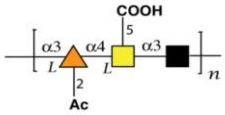  |
| 66 | 72 | Pseudomonas aerguinosa O10 10a,10c | IATS 19, OPS | 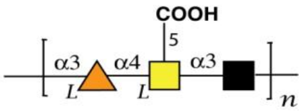  |
| 67 | 73 | Pseudomonas aerguinosa O11 11a,11b | IATS 11, OPS | 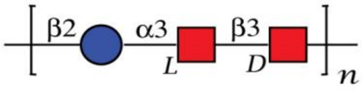  |
| 68 | 74 | Pseudomonas aerguinosa O12 12 | IATS 12, OPS Habs 12 | 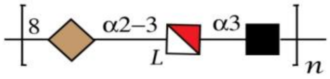  |
| 69 | 75 | Pseudomonas aerguinosa O13 13a,13c | IATS 14, OPS | 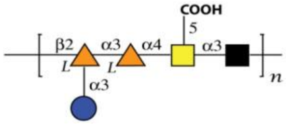  |
| 70 | 76 | Pseudomonas aerguinosa O14 14 | IATS 17,OPS Meitert X | 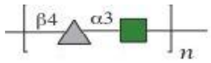  |
| 71 | 77 | Pseudomonas aerguinosa O15 15 | IATS 15, OPS | 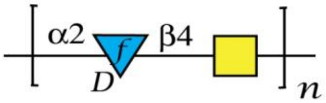  |
| 72 | 78 | Proteus vulgaris O1 (18984)* | LPS | 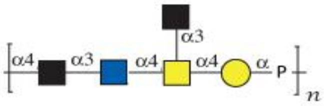  |
| 73 | 79 | Proteus vulgaris O4 (PrK 9/57) | OPS | 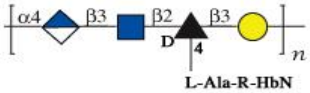  |
| 74 | 80 | Proteus vulgaris O12 (PrK 25/57) | OPS | 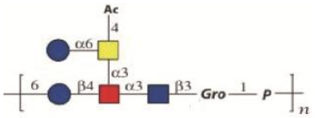  |
| 75 | 81 | Proteus vulgaris O13 (8344) | OPS | 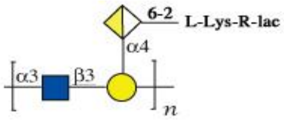  |
| 76 | 82 | Proteus vulgaris O15 (PrK 30/57) | OPS | 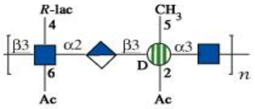  |
| 77 | 83 | Proteus vulgaris O17 (PrK 33/57) | OPS | 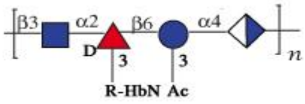  |

| | | | | |
|---|---|---|---|---|
| 78 | 84 | Proteus vulgaris O19a (PrK 37/57) | OPS | 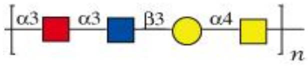 |
| 79 | 85 | Proteus vulgaris O21 (PrK 39/57)* | LPS | 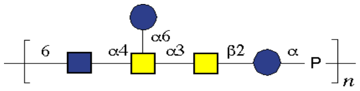 |
| 80 | 86 | Proteus vulgaris O22 (PrK 40/57) | OPS | 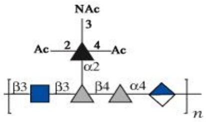 |
| 81 | 88 | Proteus vulgaris O25 (PrK 48/57) | OPS | 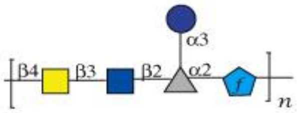 |
| 82 | 89 | Proteus vulgaris O34 (4669)* | LPS | 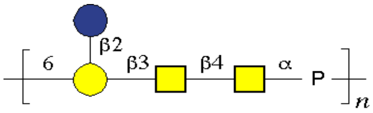 |
| 83 | 90 | Proteus vulgaris O37a,b (PrK 63/57) | OPS | 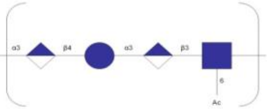 |
| 84 | 91 | Proteus vulgaris O37a,c (PrK 72/57) | OPS | 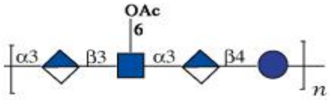 |
| 85 | 92 | Proteus vulgaris O44 (PrK 67/57) | OPS | 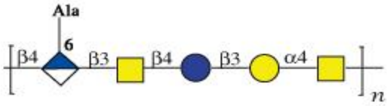 |
| 86 | 93 | Proteus vulgaris O45 (4680) | OPS | 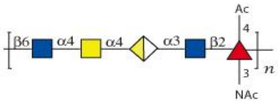 |
| 87 | 94 | Proteus vulgaris O53 (TG 276-10) | OPS | 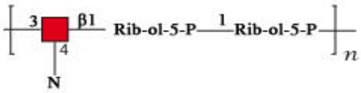 |
| 88 | 95 | Proteus vulgaris O54a,54c (TG 103) | OPS | 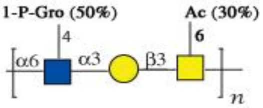 |
| 89 | 96 | Proteus vulgaris O55 (TG 155) | OPS | 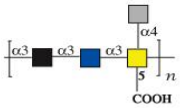 |
| 90 | 97 | Proteus vulgaris O65 (TG 251) | OPS | 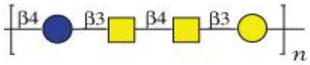 |

| | | | | |
|---|---|---|---|---|
| 91 | 98 | Proteus mirabilis O6 (PrK 14/57) | OPS | 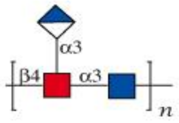  |
| 92 | 99 | Proteus mirabilis O11 (PrK 24/57) | OPS | 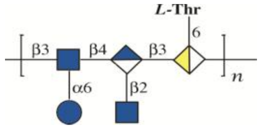  |
| 93 | 100 | Proteus mirabilis O13 (PrK 26/57) | OPS | 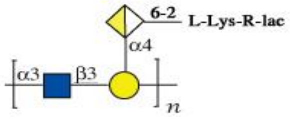  |
| 94 | 101 | Proteus mirabilis O14a,14b (PrK 29/57) | OPS | 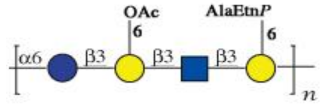  |
| 95 | 102 | Proteus mirabilis O16 (4652) | OPS | 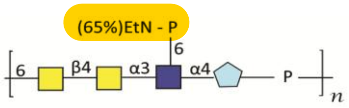  |
| 96 | 103 | Proteus mirabilis O17 (PrK 32/57) | OPS | 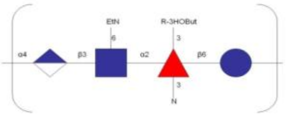 |
| 97 | 104 | Proteus mirabilis O23a,b,d (PrK 42/57) | OPS | 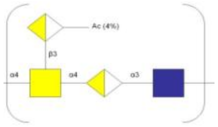 |
| 98 | 106 | Proteus mirabilis O26 (PrK 49/57) | OPS | 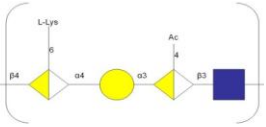 |
| 99 | 107 | Proteus mirabilis O27 (PrK 50/57) | OPS | 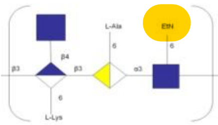 |
| 100 | 108 | Proteus mirabilis O28 (PrK 51/57) | OPS | 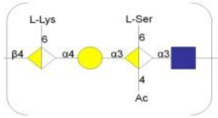  |
| 101 | 109 | Proteus mirabilis O29a (PrK 52/57) | OPS | 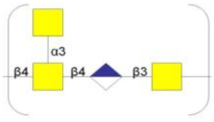  |
| 102 | 110 | Proteus mirabilis O40 (10703) | OPS | 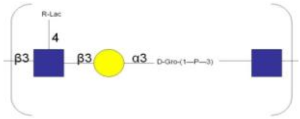  |
| 103 | 111 | Proteus mirabilis O41 (PrK 67/57) | OPS | 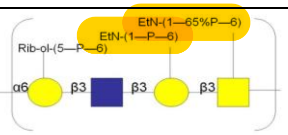  |

| | | | | |
|---|---|---|---|---|
| 104 | 112 | Proteus mirabilis O51 (19011)* | LPS | 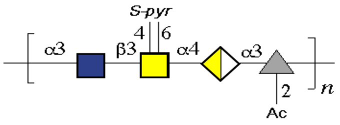  |
| 105 | 113 | Proteus mirabilis O74 (10705, OF) | OPS | 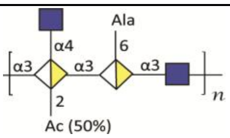  |
| 106 | 114 | Proteus mirabilis O75 (10702, OC) | OPS | 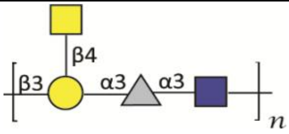  |
| 107 | 115 | Proteus mirabilis O77 (3 B-m) | OPS | 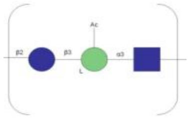 |
| 108 | 116 | Proteus penneri O31a (26) | OPS | 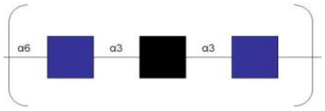  |
| 109 | 117 | Proteus penneri O52 (15) | OPS | 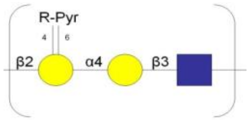  |
| 110 | 118 | Proteus penneri O58 (12) | OPS | 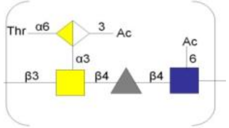  |
| 111 | 119 | Proteus penneri O59 (9) | OPS | 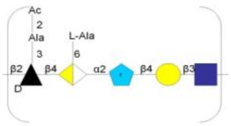  |
| 112 | 120 | Proteus penneri O61 (21) | OPS | 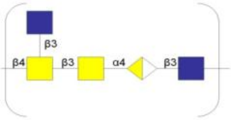  |
| 113 | 121 | Proteus penneri O62 (41) | OPS | 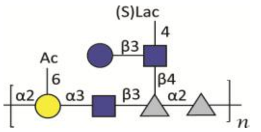  |
| 114 | 122 | Proteus penneri O63 (22) | OPS | 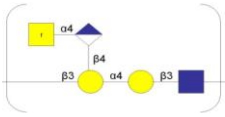  |
| 115 | 123 | Proteus penneri O64a,b,c (27) | OPS | 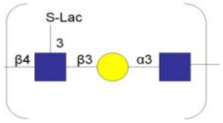  |
| 116 | 124 | Proteus penneri O65 (34) | OPS | 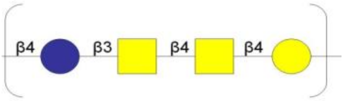  |

| | | | | |
|---|---|---|---|---|
| 117 | 125 | Proteus penneri O67 (8) | OPS | 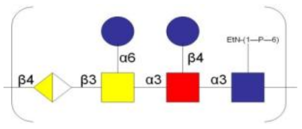 |
| 118 | 126 | Proteus penneri O68 (63) | OPS | 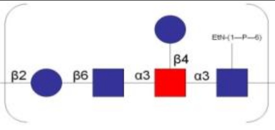 |
| 119 | 127 | Proteus penneri O70 (60) | OPS | 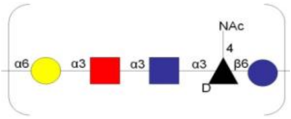 |
| 120 | 128 | Proteus penneri O73a,b (103) | OPS | 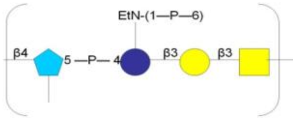 |
| 121 | 129 | Proteus myxofaciens O60 | OPS | 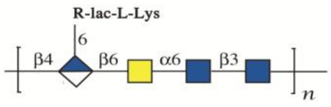 |
| 122 | 130 | Proteus O56 (genomospecies 4) | OPS | 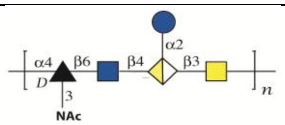 |
| 123 | 131 | Providencia stuartii O4 | OPS | 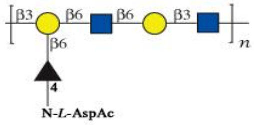 |
| 124 | 132 | Providencia stuartii O18 | OPS | 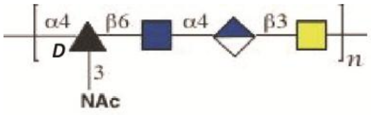 |
| 125 | 133 | Providencia stuartii O20* | LPS | 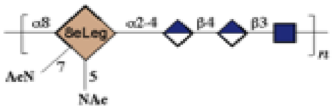 |
| 126 | 134 | Providencia stuartii O43 | OPS | 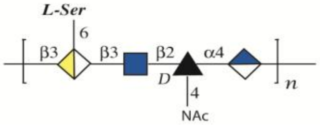 |
| 127 | 135 | Providencia stuartii O44 | OPS | 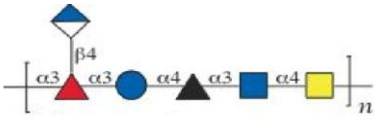 |
| 128 | 136 | Providencia stuartii O47 | OPS | 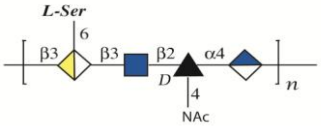 |
| 129 | 137 | Providencia stuartii O47, Core 9 | OPS | 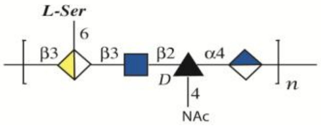 |

| | | | | |
|---|---|---|---|---|
| 130 | 138 | Providencia stuartii O49, Core 1 | OPS | 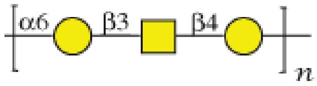 |
| 131 | 139 | Providencia stuartii O57 | OPS | 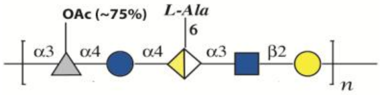 |
| 132 | 140 | Providencia alcalifaciens O5 | OPS | 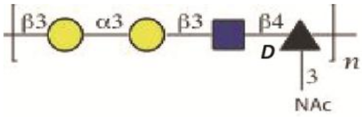 |
| 133 | 141 | Providencia alcalifaciens O6* | LPS | 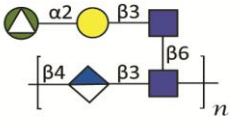 |
| 134 | 142 | Providencia alcalifaciens O19 | OPS | 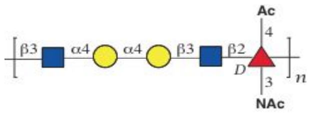 |
| 135 | 143 | Providencia alcalifaciens O19 | LPS | 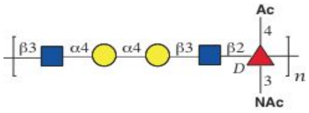 |
| 136 | 144 | Providencia alcalifaciens O19 | LPS/NaOH | 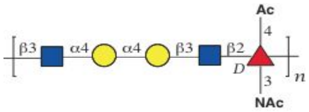 |
| 137 | 145 | Providencia alcalifaciens O21 | OPS | 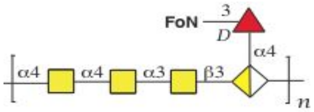 |
| 138 | 146 | Providencia alcalifaciens O23 | OPS | 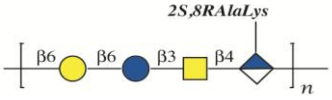 |
| 139 | 147 | Providencia alcalifaciens O27 | OPS | 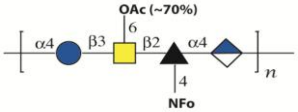 |
| 140 | 148 | Providencia alcalifaciens O29 | OPS | 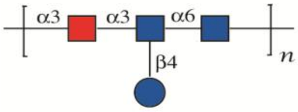 |
| 141 | 149 | Providencia alcalifaciens O30 | OPS | 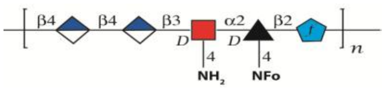 |
| 142 | 150 | Providencia alcalifaciens O32 | OPS | 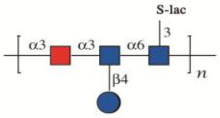 |

| | | | | |
|---|---|---|---|---|
| 143 | 151 | Providencia alcalifaciens O36* | LPS-NH4OH | 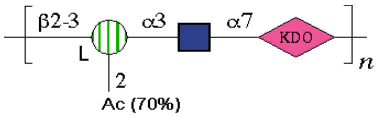 |
| 144 | 152 | Providencia alcalifaciens O39 | OPS | 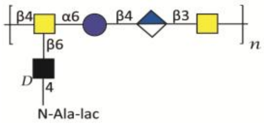 |
| 145 | 153 | Providencia rustigianii O14 | OPS | 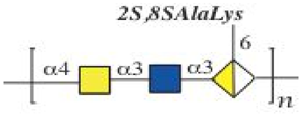 |
| 146 | 154 | Providencia rustigianii O16 | OPS | 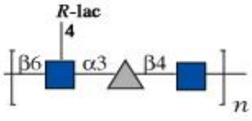 |
| 147 | 155 | Providencia rustigianii O34 | OPS | 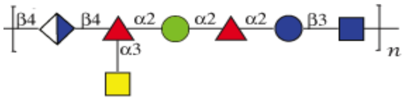 |
| 148 | 156 | Yersinia pestis, KM260(11)-∆0187 | LPS | 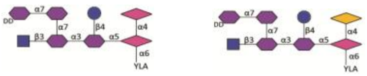 |
| 149 | 157 | Yersinia pestis, KM260(11)-∆0187 | Core oligo saccharide | 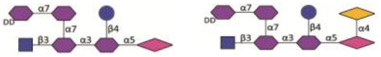 |
| 150 | 158 | Yersinia pestis, KM260(11)-∆rfe | LPS | 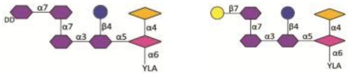 |
| 151 | 159 | Yersinia pestis, KM260(11)-∆rfe | Core oligo saccharide | 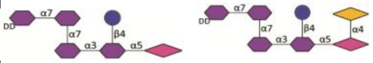 |
| 152 | 160 | Yersinia pestis, 1146-25 | LPS | 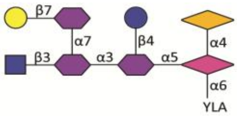 |
| 153 | 161 | Yersinia pestis 1146-25 | Core oligo saccharide | 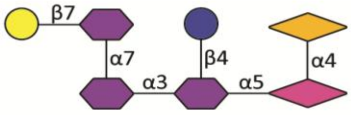 |
| 154 | 162 | Yersinia pestis, 1146-37 | LPS | 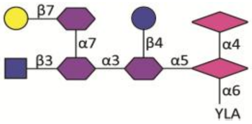 |
| 155 | 163 | Yersinia pestis, 1146-37 | Core oligo saccharide | 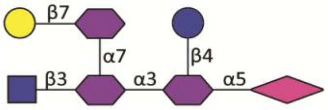 |

| | | | | |
|---|---|---|---|---|
| 156 | 164 | Yersinia pestis, KM218-37 | LPS | 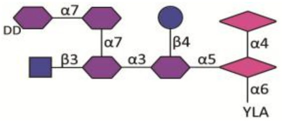  |
| 157 | 165 | Yersinia pestis, KM218-37 | Core oligo saccharide | 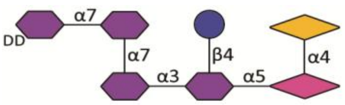  |
| 158 | 166 | Yersinia pestis, KM218-25 | LPS | 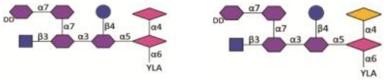  |
| 159 | 167 | Yersinia pestis, KM218-25 | Core oligo saccharide | 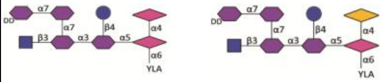  |
| 160 | 168 | Yersinia pestis, KM260(11)-ΔpmrF | LPS | 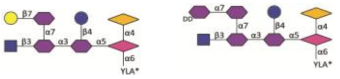  |
| 161 | 169 | Yersinia pestis, KM260(11)-ΔpmrF | Core oligo saccharide | 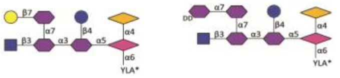  |
| 162 | 170 | Yersinia pestis, KM260(11)-Δ0186 | LPS | 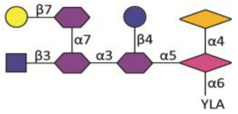  |
| 163 | 171 | Yersinia pestis, KM260(11)-Δ0186 | Core oligo saccharide | 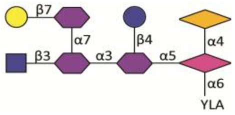  |
| 164 | 172 | Yersinia pestis, KM260(11)-ΔwaaQ | LPS | 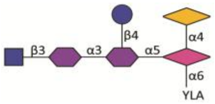  |
| 165 | 173 | Yersinia pestis, KM260(11)-ΔwaaQ | Core oligo saccharide | 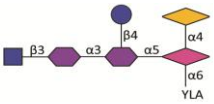  |
| 166 | 174 | Yersinia pestis, KM260(11)-ΔwaaL | LPS | 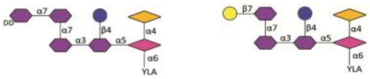  |
| 167 | 175 | Yersinia pestis, KM260(11)-25 | LPS | 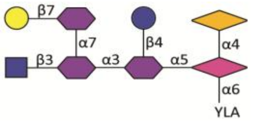  |
| 168 | 176 | Yersinia pestis, KM260(11)-25 | Core oligo saccharide | 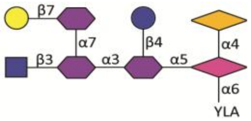  |

| | | | | |
|---|---|---|---|---|
| 169 | 177 | Yersinia pestis, KM260(11)-37 | Core oligo saccharide | 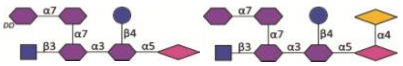 |
| 170 | 178 | Yersinia pestis, KIMD1-37 | Core oligo saccharide | 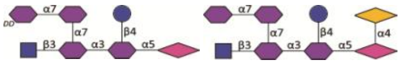 |
| 171 | 179 | Yersinia pestis, KIMD1-25 | Core oligo saccharide | 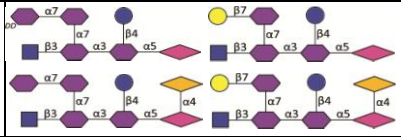 |
| 172 | 180 | Yersinia pestis, 11M-25 | LPS | 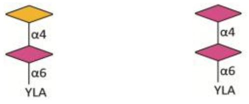 |
| 173 | 181 | Yersinia pestis, 11M-37 | LPS | 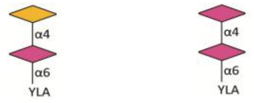 |
| 174 | 182 | Proteus vulgaris O23a, 23b, 23c (CCUG 10701) | OPS | 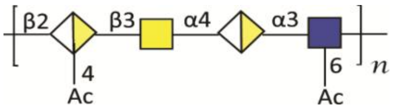 |
| 175 | 183 | Proteus vulgaris O24 (PrK 47/57) | LPSOH | 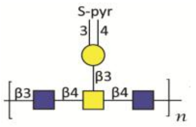 |
| 176 | 184 | Yersinia pestis KM260(11)-6C | LPS | |
| 177 | 185 | Yersinia pestis 260(11)-37C-186 | LPS | 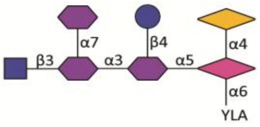 |
| 178 | 186 | Yersinia pestis 260(11)-37C-187 | LPS | 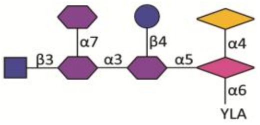 |
| 179 | 187 | Yersinia pestis 260(11)-37C-416 | LPS | 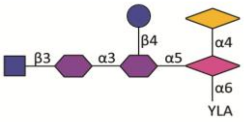 |
| 180 | 188 | Yersinia pestis 260(11)-37C-417 | LPS | |
| 181 | 189 | Yersinia pestis P-1680-25C | OS | |

| | | | | |
|---|---|---|---|---|
| 182 | 190 | Yersinia pestis P-1680-37C | LPS | |
| 183 | 191 | Yersinia pestis I-2377-25C | OS | |
| 184 | 192 | Yersinia pestis I-2377-37C | LPS | |
| 185 | 193 | Francisella novicida OPS | OPS | 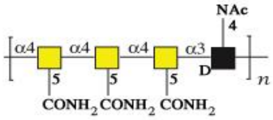 |
| 186 | 194 | Francisella tularensis OPS | OPS | 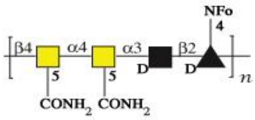 |
| 187 | 195 | Klebsiella O1 OPS | OPS | 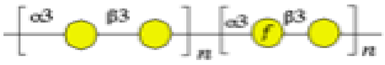 |
| 188 | 196 | Klebsiella O2a OPS | OPS | 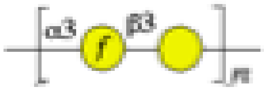 |
| 189 | 197 | Klebsiella O2ac OPS | OPS | 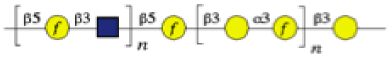 |
| 190 | 198 | Klebsiella O3 OPS | OPS | 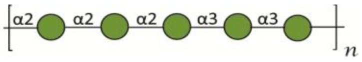 |
| 191 | 199 | Klebsiella O4 OPS | OPS | 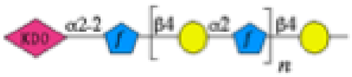 |
| 192 | 200 | Klebsiella O5 OPS | OPS | 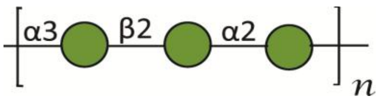 |
| 193 | 201 | Klebsiella O8 OPS | OPS | 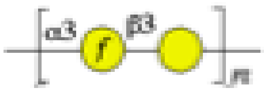 |
| 194 | 202 | Klebsiella O12 OPS | OPS | 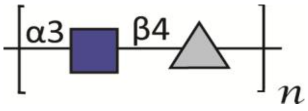 |

| | | | | |
|---|---|---|---|---|
| 195 | 203 | Shigella boydii type 1 | LPSOH | 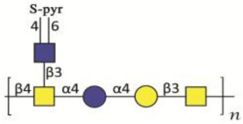 |
| 196 | 204 | Shigella boydii type 3 | OPS | 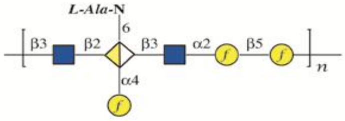 |
| 197 | 205 | Shigella boydii type 5 | OPS | 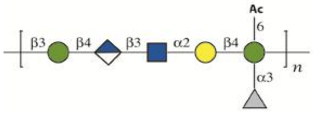 |
| 198 | 206 | Shigella boydii type 9 | OPS | 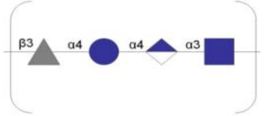 |
| 199 | 207 | Shigella boydii type 11 | OPS | 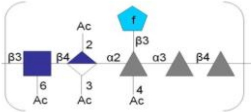 |
| 200 | 208 | Shigella boydii type 12 | OPS | 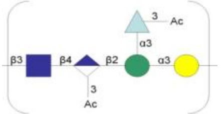 |
| 201 | 209 | Shigella boydii type 15 | OPS | 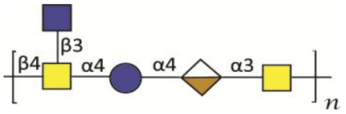 |
| 202 | 210 | Shigella boydii type 16 | OPS | 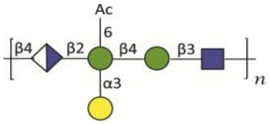 |
| 203 | 211 | Shigella boydii type 17 | OPS | 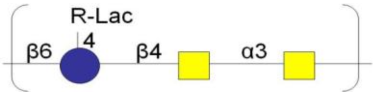 |
| 204 | 212 | Shigella boydii type 18 | OPS | 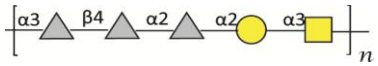 |
| 205 | 213 | Escherichia coli O49 | OPS | 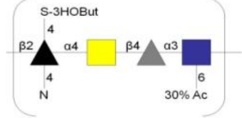 |
| 206 | 214 | Escherichia coli O52 | OPS | 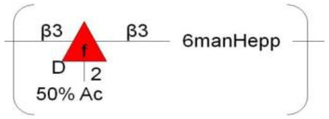 |
| 207 | 215 | Escherichia coli O58 | OPS | 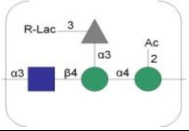 |

| | | | | |
|---|---|---|---|---|
| 208 | 216 | Escherichia coli O61 | LPSOH | 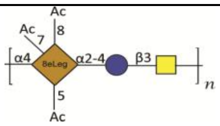  |
| 209 | 217 | Escherichia coli O73 | OPS | 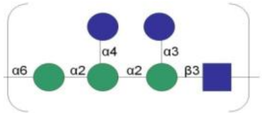  |
| 210 | 218 | Escherichia coli O112ab | OPS | 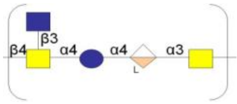  |
| 211 | 219 | Escherichia coli O118 | OPS | 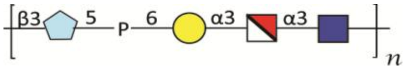  |
| 212 | 220 | Escherichia coli O125 | OPS | 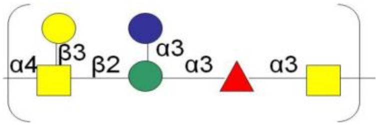  |
| 213 | 221 | Escherichia coli O151 | OPS | 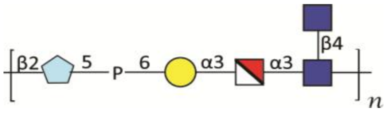  |
| 214 | 222 | Escherichia coli O168 | OPS | 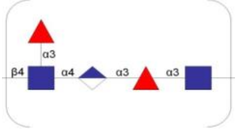  |
| 215 | 223 | Shigella dysenteriae type 2 | LPSOH | 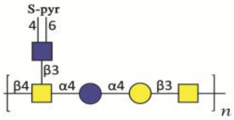  |
| 216 | 224 | Shigella dysenteriae type 4 | OPS | 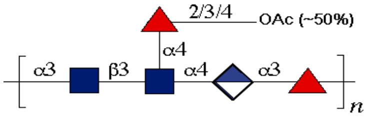  |
| 217 | 225 | Shigella dysenteriae type 5 | OPS | 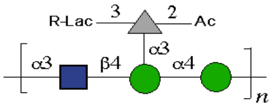  |
| 218 | 226 | Shigella dysenteriae type 6 SR-strain | SR-strain | 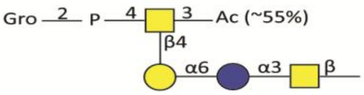  |
| 219 | 227 | Shigella dysenteriae type 7 | OPS | 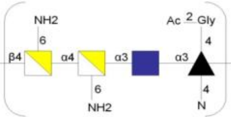  |
| 220 | 228 | Shigella dysenteriae type 8 (Russian) | OPS | 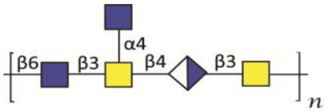  |

| | | | | |
|---|---|---|---|---|
| 221 | 229 | Shigella dysenteriae type 9 | OPS | 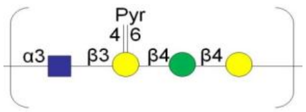 |
| 222 | 231 | Escherichia coli O111:B4 LPS-solution at 1 mg/mL | L5293-2ML (LPS) (Sigma) | 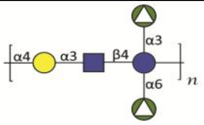 |
| 223 | 232 | Escherichia coli O26:B6 LPS-solution at 1 mg/mL | L5543-2ML (LPS) (Sigma) | 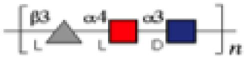 |
| 224 | 233 | Escherichia coli O55:B5 LPS-solution at 1 mg/mL | L5418-2ML (LPS) (Sigma) | 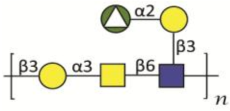 |
| 225 | 234 | Escherichia coli O127:B8 LPS-solution at 1 mg/mL | L5668-2ML (LPS) (Sigma) | 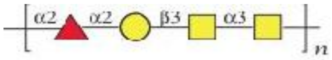 |
| 226 | 235 | Streptococcus pneumoniae type 1 (Danish type 1) | 161-X // Capsular PS | 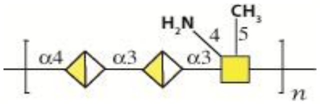 |
| 227 | 236 | Streptococcus pneumoniae type 2 (Danish type 2) | 165-X// Capsular PS | 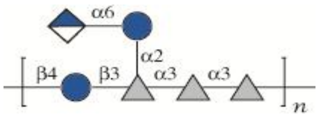 |
| 228 | 237 | Streptococcus pneumoniae type 3 (Danish type 3) | 169-X// Capsular PS | 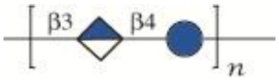 |
| 229 | 238 | Streptococcus pneumoniae type 4 (Danish type 4) | 173-X// Capsular PS | 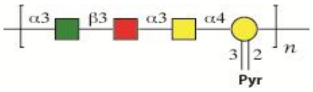 |
| 230 | 239 | Streptococcus pneumoniae type 5 (Danish type 5) | 177-X// Capsular PS | 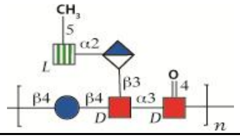 |
| 231 | 240 | Streptococcus pneumoniae type 8 (Danish type 8) | 185-X// Capsular PS | 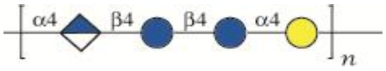 |
| 232 | 241 | Streptococcus pneumoniae type 9 (Danish type 9N) | 189-X// Capsular PS | 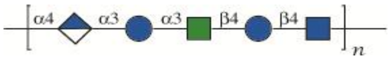 |
| 233 | 242 | Streptococcus pneumoniae type 12 (Danish type 12F) | 193-X// Capsular PS | 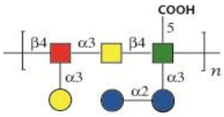 |

| | | | | |
|---|---|---|---|---|
| 234 | 243 | Streptococcus pneumoniae type 14 (Danish type 14) | 197-X// Capsular PS | 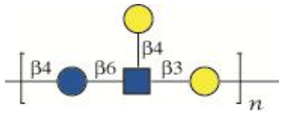 |
| 235 | 244 | Streptococcus pneumoniae type 17 (Danish type 17F) | 201-X// Capsular PS | 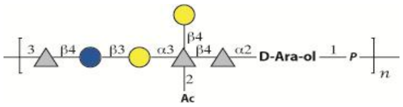 |
| 236 | 245 | Streptococcus pneumoniae type 19 (Danish type 19F) | 205-X// Capsular PS | 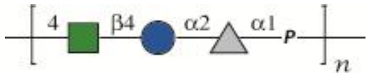 |
| 237 | 246 | Streptococcus pneumoniae type 20 (Danish type 20) | 209-X// Capsular PS | 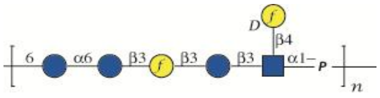 |
| 238 | 247 | Streptococcus pneumoniae type 22 (Danish type 22F) | 213-X// Capsular PS | 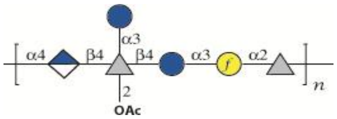 |
| 239 | 248 | Streptococcus pneumoniae type 23 (Danish type 23F) | 217-X// Capsular PS | 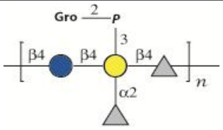 |
| 240 | 249 | Streptococcus pneumoniae type 26 (Danish type 6B) | 225-X// Capsular PS | 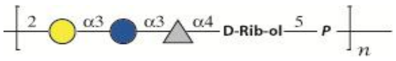 |
| 241 | 250 | Streptococcus pneumoniae type 34 (Danish type 10A) | 229-X// Capsular PS | 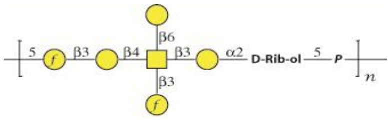 |
| 242 | 251 | Streptococcus pneumoniae type 43 (Danish type 11A) | 233-X// Capsular PS | 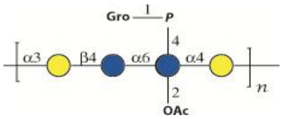 |
| 243 | 252 | Streptococcus pneumoniae type 51 (Danish type 7F) | 237-X// Capsular PS | 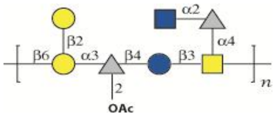 |
| 244 | 253 | Streptococcus pneumoniae type 54 (Danish type 15B) | 241-X// Capsular PS | 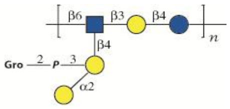 |
| 245 | 254 | Streptococcus pneumoniae type 56 (Danish type 18C) | 245-X// Capsular PS | 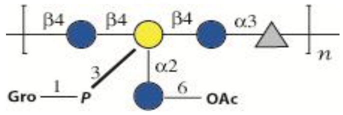 |
| 246 | 255 | Streptococcus pneumoniae type 57 (Danish type 19A) | 249-X// Capsular PS | 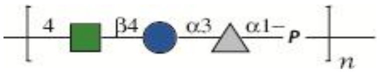 |

| | | | | |
|---|---|---|---|---|
| 247 | 256 | Streptococcus pneumoniae type 68 (Danish type 9V) | 253-X// Capsular PS | 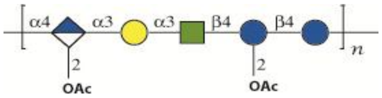 |
| 248 | 257 | Streptococcus pneumoniae type 70 (Danish type 33F) | 257-X// Capsular PS | 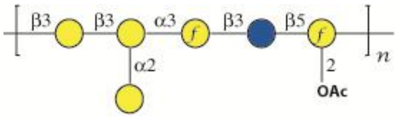 |
| 249 | 258 | Yersinia pestis KM218-6C | OS | 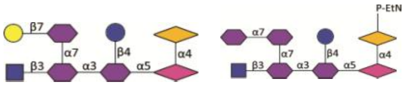 |
| 250 | 259 | Yersinia pestis KM260(11)-yjhW-6C | OS | |
| 251 | 260 | Yersinia pestis KM260(11)-wabD/waaL | OS | |
| 252 | 261 | Yersinia pestis KM260(11)-wabC/waaL | OS | |
| 253 | 262 | Yersinia pseudotuberculosis 85pCad-37C | OS | |
| 254 | 263 | Yersinia pseudotuberculosis 85pCad-20C | OS | |
| 255 | 264 | Yersinia pseudotuberculosis O:2a | PS | 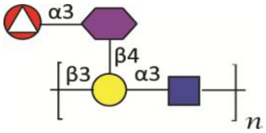 |
| 256 | 265 | Yersinia pseudotuberculosis O:2a-dhmA | PS | 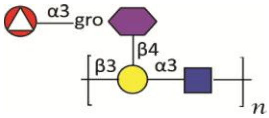 |
| 257 | 266 | Yersinia pseudotuberculosis O:2c | PS | 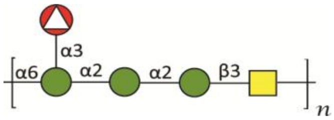 |
| 258 | 267 | Yersinia pseudotuberculosis O:3 | PS | 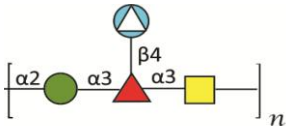 |
| 259 | 268 | Yersinia pseudotuberculosis O:4b | PS | 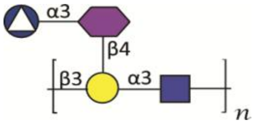 |

| | | | | |
|---|---|---|---|---|
| 260 | 269 | Proteus vulgaris O2 (OX2) | PS | 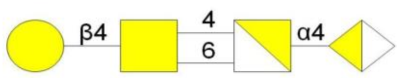  |
| 261 | 270 | Proteus mirabilis O3ab (S1959) | PS | 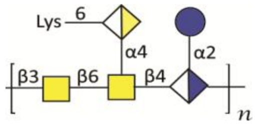  |
| 262 | 271 | Proteus mirabilis O5 (PrK 12/57) | PS | 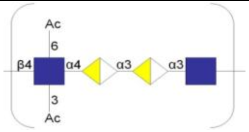  |
| 263 | 272 | Proteus mirabilis O9 (PrK 18/57) | PS | 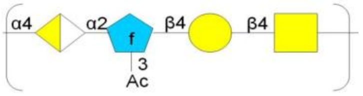  |
| 264 | 273 | Proteus mirabilis O11 (9B-m) | PS | 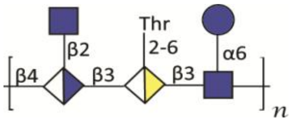  |
| 265 | 274 | Proteus penneri O17 (16) | PS | 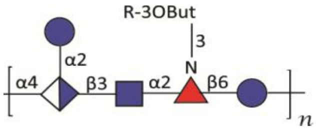  |
| 266 | 275 | Proteus mirabilis O18 (PrK 34/57) | LPSOH | 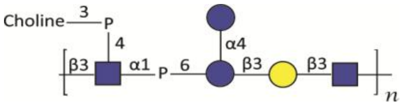  |
| 267 | 276 | Proteus mirabilis O20 (PrK 38/57) | LPSOH | 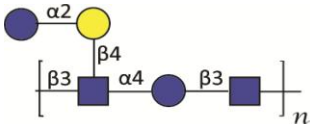  |
| 268 | 277 | Proteus penneri O31ab (28) | PS | 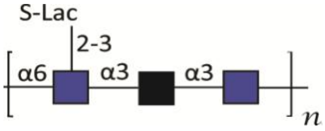  |
| 269 | 278 | Proteus mirabilis O33 (D52) | PS | 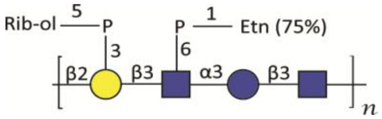  |
| 270 | 279 | Proteus mirabilis O43 (PrK 69/57) | PS | 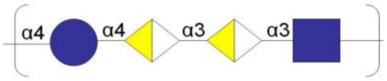  |
| 271 | 280 | Proteus vulgaris O47 (PrK 73/57) | Not stated | 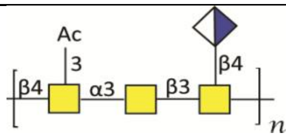  |
| 272 | 281 | Proteus mirabilis O49 (PrK 75/57) | PS | 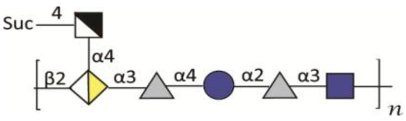  |

| | | | | |
|---|---|---|---|---|
| 273 | 282 | Proteus mirabilis O54ab (OE) | PS | 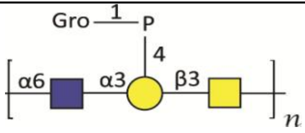  |
| 274 | 283 | Proteus penneri O73ac (75) | PS | 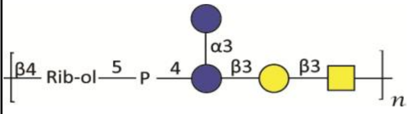  |
| 275 | 284 | Proteus vulgaris O76 (HSC438) | PS | 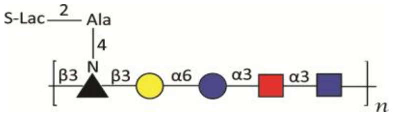  |
| 276 | 285 | Shigella flexneri type 1a | PS | 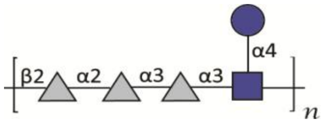  |
| 277 | 286 | Shigella flexneri type 1b | PS | 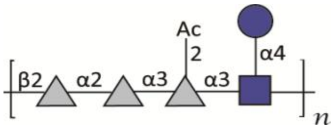  |
| 278 | 287 | Shigella flexneri type 2a | PS | 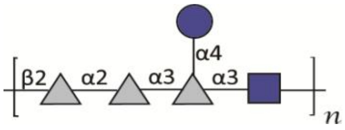  |
| 279 | 288 | Shigella flexneri type 2b | PS | 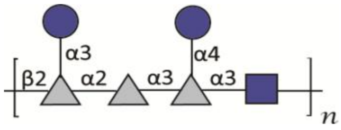  |
| 280 | 289 | Shigella flexneri type 3a | PS | 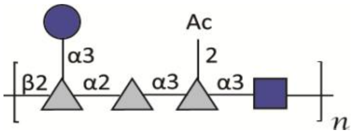  |
| 281 | 290 | Shigella flexneri type 3b | PS | 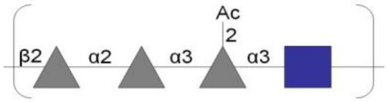  |
| 282 | 291 | Shigella flexneri type 4a | PS | 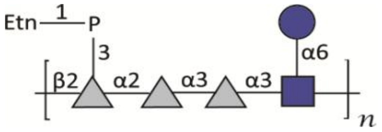  |
| 283 | 292 | Shigella flexneri type 4b | PS | 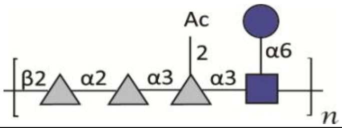  |
| 284 | 293 | Shigella flexneri type 5b | PS | 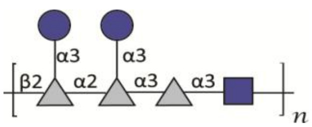  |
| 285 | 294 | Shigella flexneri type 6a | PS | 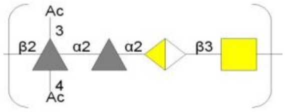  |

| | | | | |
|---|---|---|---|---|
| 286 | 295 | Shigella flexneri type 6 | PS | 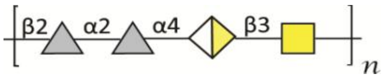 |
| 287 | 296 | Shigella flexneri type X | PS | 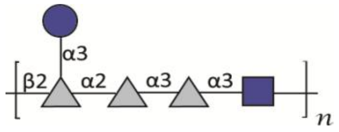 |
| 288 | 297 | Shigella dysenteriae type 1 | PS | 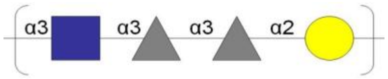 |
| 289 | 298 | Shigella boydii type 6 | PS | 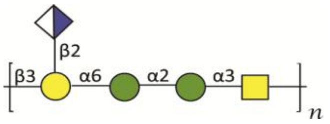 |
| 290 | 299 | Shigella boydii type 7 | PS | 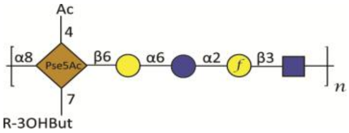 |
| 291 | 300 | Shigella boydii type 8 | PS | 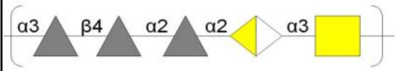 |
| 292 | 301 | Shigella boydii type 13 | LPSOH | 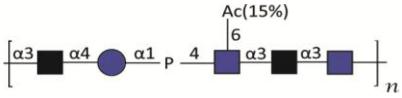 |
| 293 | 302 | Shigella boydii type 14 | LPSOH | 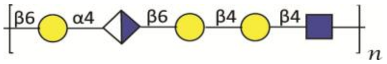 |
| 294 | 303 | Escherichia coli O71 | PS | 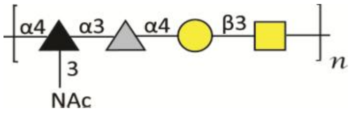 |
| 295 | 304 | Escherichia coli O85 | PS | 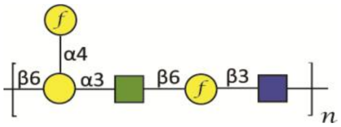 |
| 296 | 305 | Escherichia coli O99 | PS | 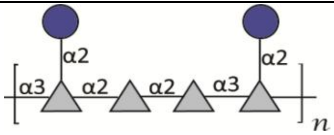 |
| 297 | 306 | Escherichia coli O145 | LPSOH | 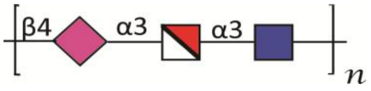 |
| 298 | 307 | Escherichia coli O107 | PS | 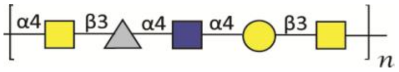 |

| | | | | |
|---|---|---|---|---|
| 299 | 308 | Salmonella enterica O17 | PS | 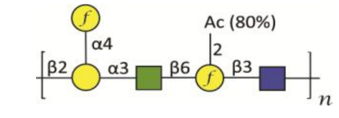 |
| 300 | 309 | Salmonella enterica O28 | PS | 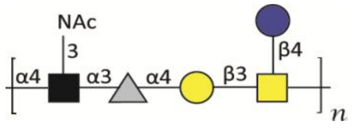 |
| 301 | 310 | Salmonella enterica O47 | PS | 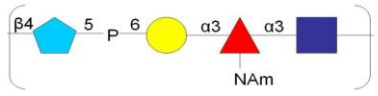 |
| 302 | 311 | Salmonella enterica O55 | PS | 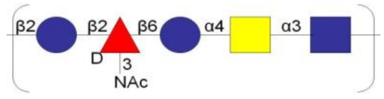 |
| 303 | 312 | Escherichia coli K92 | CPS | 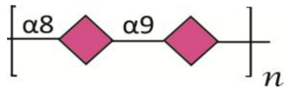 |
| 304 | 313 | Escherichia coli K5 | CPS | 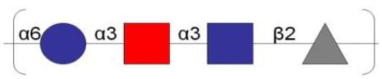 |
| 305 | 314 | Escherichia coli K13 | CPS | 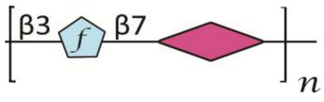 |
| 306 | 315 | Neisseria meningitidis Group C | CPS | 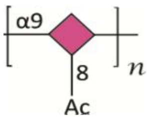 |
| 307 | 316 | Davanat | | 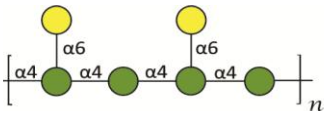 |
| 308 | 317 | Laminarin | | 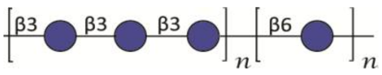 |
| 309 | 318 | Yeast Mannan | | |
| 310 | 319 | Escherichia coli O86 | | 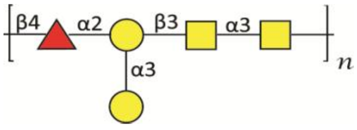 |
| 311 | 320 | Galactomannan DAVANT (160102) Pro-Pharmacenti | | 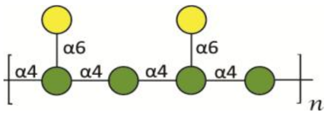 |

| | | | | |
|---|---|---|---|---|
| 312 | 321 | Yeast Mannan Sigma M-3640 | | |
| 313 | 322 | 1-2 Mannan Acetobacter methanolicus MB135 | | 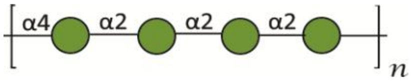 |

***Extended symbol nomenclature for bacterial glycans** (Reference: Stowell SR, Arthur CM, McBride R, Berger O, Razi N, Heimburg-Molinaro J, Rodrigues LC, Gourdine J-P, Noll AJ, von Gunten S, *et al*. 2014. Microbial glycan microarrays define key features of host-microbial interactions. *Nat. Chem. Biol.*, 10:470-476

SI-Spreadsheet-4

| Glycan name | Lectins | | | | | | | | |
|---|---|---|---|---|---|---|---|---|---|
| | MBL | ΔN-SPD | CL-K1 | CRP | FicH | Gal-3 | Gal-4 | ZG16P | PTX3 |
| POS1 | -19190.00 | -11343.70 | 1008.08 | -699.42 | -2634.42 | -6159.00 | 26.00 | 23611.58 | 57.94 |
| NEG | 226.67 | 5.50 | 40.00 | 1.00 | 1.00 | 52.00 | 1.00 | 1.00 | 1.55 |
| G0001 | 281.00 | 810.67 | 23.67 | 1.00 | 24.00 | 1.00 | 1.00 | 1.00 | 272.78 |
| G0002 | 1775.67 | 1.00 | 1.00 | 44.67 | 38.33 | 1.00 | 1.00 | 1.00 | 368.15 |
| G0003 | 520.33 | 499.00 | 117.00 | 1.00 | 1.00 | 1.00 | 1.00 | 1.00 | 838.33 |
| G0004 | 366.67 | 1.00 | 21.33 | 133.33 | 1.00 | 1.00 | 45.00 | 1.00 | 469.45 |
| G0005 | 59.33 | 1.00 | 1.00 | 1.00 | 1.00 | 1.00 | 121.00 | 1.00 | 120.49 |
| G0006 | 1017.67 | 510.00 | 96.67 | 1.00 | 1.00 | 1.00 | 18.00 | 2997.33 | 606.18 |
| G0007 | 459.67 | 1.00 | 1.00 | 1.00 | 1.00 | 64.00 | 49.00 | 1.00 | 255.80 |
| G0008 | 1592.33 | 334.00 | 112.00 | 8667.33 | 16.67 | 44.00 | 1.00 | 1.00 | 711.92 |
| G0009 | 325.33 | 219.00 | 107.67 | 367.33 | 1.00 | 80.00 | 203.00 | 1.00 | 480.52 |
| G0010 | 683.33 | 1.00 | 32.33 | 653.33 | 1.00 | 88.00 | 67.00 | 1.00 | 462.78 |
| G0011 | 271.00 | 8.00 | 122.33 | 1.00 | 1.00 | 1.00 | 1.00 | 1.00 | 591.41 |
| G0012 | 277.33 | 1.00 | 1.00 | 1.00 | 1.00 | 1.00 | 1.00 | 1.00 | 678.64 |
| G0013 | 267.33 | 1.00 | 1.00 | 1.00 | 5.00 | 5.00 | 111.00 | 1.00 | 393.31 |
| G0014 | 248.00 | 1.00 | 12.67 | 67.00 | 50.67 | 1.00 | 61.00 | 5140.00 | 671.23 |
| G0015 | 185.00 | 1.00 | 1.00 | 1.00 | 18.67 | 1.00 | 1.00 | 2741.33 | 215.86 |
| G0016 | 306.33 | 432.67 | 18.33 | 1.00 | 1.00 | 54.00 | 148.00 | 172.00 | 457.61 |
| G0017 | 141.33 | 229.00 | 97.00 | 1.00 | 1.00 | 125.00 | 154.00 | 1.00 | 464.27 |
| G0018 | 137.67 | 1.00 | 1.00 | 538.00 | 1.00 | 19.00 | 1.00 | 1.00 | 569.22 |
| G0019 | 3714.33 | 509.67 | 1.00 | 930.00 | 8.67 | 20.00 | 36.00 | 1.00 | 637.24 |
| G0020 | 107.33 | 1.00 | 1.00 | 9.33 | 16.67 | 1.00 | 1.00 | 1.00 | 820.58 |
| G0021 | 83.33 | 1.00 | 1.00 | 1.00 | 1.00 | 1.00 | 23.00 | 1.00 | 446.53 |
| G0022 | 356.00 | 159.33 | 3.33 | 1.00 | 1.00 | 1.00 | 31.00 | 1.00 | 385.15 |
| G0023 | 138.67 | 1.00 | 1.00 | 805.33 | 1.00 | 1.00 | 142.00 | 1028.00 | 767.34 |
| G0024 | 745.67 | 129.67 | 1.00 | 258.00 | 1.00 | 1.00 | 1.00 | 1.00 | 542.62 |
| G0025 | 263.33 | 341.67 | 89.67 | 1.00 | 17.00 | 98.00 | 1.00 | 1.00 | 215.14 |
| G0026 | 99.67 | 27.00 | 1.00 | 1.00 | 10.33 | 70.00 | 133.00 | 1.00 | 267.62 |
| G0027 | 1787.33 | 1017.00 | 144.67 | 1.00 | 1.00 | 1.00 | 1.00 | 1.00 | 741.46 |
| G0028 | 100.67 | 1.00 | 1.00 | 9160.00 | 1.00 | 1.00 | 1.00 | 1.00 | 751.07 |
| G0029 | 253.33 | 1.00 | 1.00 | 1.00 | 42.67 | 1.00 | 1.00 | 1.00 | 377.04 |
| G0030 | 488.67 | 421.33 | 1.00 | 249.33 | 1.33 | 1.00 | 139.00 | 1.00 | 654.96 |
| G0031 | 179.33 | 1.00 | 1.00 | 1.00 | 1.00 | 56.00 | 1.00 | 1.00 | 510.83 |

| | | | | | | | | | |
|---|---|---|---|---|---|---|---|---|---|
| G0032 | 251.33 | 557.33 | 1.00 | 1.00 | 2.67 | 1.00 | 9.00 | 1.00 | 445.03 |
| G0033 | 327.67 | 123.67 | 40.33 | 1.00 | 1.00 | 1.00 | 1.00 | 1.00 | 144.90 |
| G0034 | 358.33 | 18.33 | 1.00 | 1.00 | 1.00 | 79.00 | 1.00 | 1.00 | 219.58 |
| G0035 | 1.00 | 1520.00 | 400.33 | 1.00 | 6.67 | 27.00 | 1618.00 | 1413.00 | 16.98 |
| G0036 | 141.33 | 1.00 | 1.00 | 1.00 | 1.00 | 25.00 | 1.00 | 1.00 | 390.33 |
| G0037 | 71.33 | 1.00 | 1.00 | 1.00 | 1.00 | 100.00 | 47.00 | 1.00 | 674.20 |
| G0038 | 432.67 | 243.33 | 1.00 | 124.67 | 32.67 | 1.00 | 1.00 | 1.00 | 489.38 |
| G0039 | 70.67 | 1.00 | 2.33 | 1.00 | 1.00 | 1.00 | 1.00 | 1.00 | 736.29 |
| G0040 | 973.00 | 1.00 | 1.00 | 1.00 | 1.00 | 58.00 | 1.00 | 1.00 | 497.53 |
| G0041 | 925.33 | 1.00 | 1.00 | 1.00 | 1.00 | 1.00 | 44.00 | 1.00 | 693.41 |
| G0042 | 40.00 | 1.00 | 1.00 | 1.00 | 1.00 | 1.00 | 93.00 | 1.00 | 180.38 |
| G0043 | 165.00 | 106.67 | 1.00 | 1.00 | 1.00 | 104.00 | 41.00 | 1.00 | 249.89 |
| G0044 | 8.00 | 34.67 | 1.00 | 584.00 | 1.00 | 215.00 | 103.00 | 1.00 | 392.55 |
| G0045 | 7374.00 | 1081.00 | 109.33 | 1.00 | 1.00 | 1.00 | 1.00 | 1.00 | 804.31 |
| G0046 | 134.67 | 391.67 | 1.00 | 1.00 | 1.00 | 1.00 | 1.00 | 1.00 | 442.09 |
| G0047 | 1.00 | 20.33 | 1.00 | 1.00 | 11.67 | 1.00 | 43.00 | 1.00 | 170.04 |
| G0048 | 60.33 | 1.00 | 1.00 | 1.00 | 2.33 | 1.00 | 1.00 | 1.00 | 359.27 |
| G0049 | 136.67 | 1.00 | 1.00 | 1.00 | 1.00 | 1.00 | 1.00 | 4113.33 | 785.84 |
| G0050 | 1.00 | 1.00 | 1.00 | 1.00 | 1.00 | 1.00 | 1.00 | 1.00 | 606.20 |
| G0051 | 3306.33 | 373.33 | 1.00 | 1204.67 | 1.00 | 1.00 | 7.00 | 1.00 | 0.02 |
| G0052 | 173.00 | 1.00 | 1.00 | 1.00 | 13.67 | 107.00 | 15.00 | 1.00 | 227.70 |
| G0053 | 283.67 | 555.00 | 1.00 | 1.00 | 1.00 | 115.00 | 24.00 | 1.00 | 663.87 |
| G0054 | 1249.00 | 669.00 | 121.67 | 72.33 | 1.00 | 1.00 | 1.00 | 1.00 | 514.54 |
| G0055 | 50.33 | 22.33 | 10.33 | 1.00 | 2.33 | 24.00 | 61.00 | 1.00 | 314.92 |
| G0056 | 43.67 | 1.00 | 1.00 | 16727.00 | 1.67 | 10.00 | 1.00 | 1.00 | 526.35 |
| G0057 | 40.33 | 1.00 | 1.00 | 1.00 | 1.00 | 1.00 | 1.00 | 1.00 | 580.33 |
| G0058 | 136.00 | 1.00 | 1.00 | 1.00 | 16.33 | 46.00 | 16.00 | 1.00 | 697.86 |
| G0059 | 1972.00 | 51.33 | 1.00 | 293.67 | 20.33 | 39.00 | 1.00 | 1.00 | 817.59 |
| G0060 | 35.00 | 1.00 | 1.00 | 1.00 | 1.00 | 65.00 | 127.00 | 1.00 | 187.03 |
| G0061 | 2646.67 | 373.67 | 1.00 | 180.67 | 1.00 | 133.00 | 144.00 | 1.00 | 0.02 |
| G0062 | 322.00 | 485.33 | 118.00 | 1383.33 | 1.00 | 205.00 | 182.00 | 1.00 | 495.31 |
| G0063 | 372.00 | 1.00 | 1.00 | 693.33 | 8.00 | 1.00 | 1.00 | 428.00 | 475.37 |
| G0064 | 107.33 | 1.00 | 1.00 | 1.00 | 1.00 | 21.00 | 1.00 | 2312.00 | 309.01 |
| G0065 | 40.33 | 1.00 | 1.00 | 1.00 | 1.00 | 1.00 | 1.00 | 1.00 | 413.25 |
| G0066 | 1.00 | 1.00 | 1.00 | 1.00 | 1.00 | 1.00 | 42.00 | 1.00 | 155.27 |

| | | | | | | | | | |
|---|---|---|---|---|---|---|---|---|---|
| G0067 | 293.00 | 376.00 | 19.67 | 1.00 | 5.67 | 1.00 | 14.00 | 1.00 | 449.47 |
| G0068 | 3.67 | 1.00 | 12.00 | 1.00 | 1.00 | 1.00 | 32.00 | 1.00 | 67.26 |
| G0069 | 296.33 | 1.00 | 1.00 | 1.00 | 1.00 | 1.00 | 1.00 | 1.00 | 239.53 |
| G0070 | 86.67 | 240.33 | 119.67 | 1.00 | 18.33 | 115.00 | 18.00 | 1.00 | 269.11 |
| G0071 | 139.33 | 20.67 | 1.00 | 1.00 | 10.00 | 47.00 | 1.00 | 1.00 | 344.48 |
| G0072 | 39.00 | 18.33 | 1.00 | 1.00 | 20.00 | 1.00 | 1.00 | 1.00 | 708.19 |
| G0073 | 1.00 | 1.00 | 1.00 | 1.00 | 1.00 | 1.00 | 1.00 | 1.00 | 550.74 |
| G0074 | 30.33 | 1.00 | 1.00 | 1.00 | 1.00 | 1.00 | 1.00 | 1542.00 | 193.67 |
| G0075 | 2751.33 | 372.00 | 1.00 | 341.33 | 8.33 | 1.00 | 11.00 | 1.00 | 437.63 |
| G0076 | 101.33 | 1.00 | 1.00 | 1.00 | 1.00 | 1.00 | 1.00 | 1.00 | 764.40 |
| G0077 | 689.00 | 205.67 | 33.33 | 1.00 | 1.00 | 1.00 | 1.00 | 1.00 | 224.72 |
| G0078 | 793.00 | 1229.00 | 1.00 | 1.00 | 1.00 | 22.00 | 1.00 | 1.00 | 786.56 |
| G0079 | 173.00 | 1.00 | 1.00 | 1.00 | 1.00 | 133.00 | 1.00 | 984.00 | 184.80 |
| G0080 | 163.00 | 1.00 | 1.00 | 254.67 | 1.00 | 1.00 | 1.00 | 1.00 | 843.48 |
| G0081 | 82.67 | 1.00 | 1.00 | 1.00 | 1.00 | 9.00 | 1.00 | 1.00 | 776.20 |
| G0082 | 46.00 | 1.00 | 1.00 | 1.00 | 1.00 | 14.00 | 1.00 | 1.00 | 618.00 |
| G0083 | 263.00 | 272.00 | 1.00 | 1.00 | 1.00 | 1.00 | 1.00 | 1.00 | 243.93 |
| G0084 | 1.00 | 1.00 | 1.00 | 1.00 | 4.33 | 1.00 | 1.00 | 1.00 | 391.07 |
| G0085 | 741.00 | 227.00 | 1.00 | 1.00 | 7.00 | 1.00 | 30.00 | 1.00 | 581.06 |
| G0086 | 715.00 | 846.33 | 1.00 | 1.00 | 1.00 | 1.00 | 1.00 | 1.00 | 344.47 |
| G0087 | 79.00 | 1.00 | 1.00 | 1.00 | 1.00 | 1.00 | 2.00 | 1.00 | 687.49 |
| G0088 | 436.00 | 480.00 | 1.00 | 1.00 | 1.00 | 102.00 | 1.00 | 1.00 | 291.25 |
| G0089 | 215.33 | 1.00 | 1.00 | 1.00 | 10.00 | 59.00 | 1.00 | 1.00 | 313.43 |
| G0090 | 5410.33 | 714.67 | 1.00 | 1.00 | 11.00 | 12.00 | 1.00 | 1.00 | 618.00 |
| G0091 | 281.33 | 387.00 | 1.00 | 1.00 | 1.00 | 28.00 | 1.00 | 1.00 | 576.60 |
| G0092 | 87.33 | 1.00 | 1.00 | 1.00 | 1.00 | 1.00 | 1.00 | 1.00 | 342.26 |
| G0093 | 1.00 | 1.00 | 1.00 | 586.33 | 1.00 | 1.00 | 1.00 | 1.00 | 360.75 |
| G0094 | 1.00 | 1.00 | 1.00 | 1.00 | 1.00 | 1.00 | 1.00 | 1.00 | 885.64 |
| G0095 | 32.67 | 1.00 | 1.00 | 1.00 | 1.00 | 1.00 | 140.00 | 1.00 | 78.36 |
| G0096 | 160.00 | 31.67 | 42.33 | 11805.00 | 17.33 | 1.00 | 1.00 | 1.00 | 483.49 |
| G0097 | 300.67 | 74.67 | 1.00 | 7010.33 | 1.00 | 119.00 | 2.00 | 1.00 | 345.98 |
| G0098 | 138.67 | 449.00 | 22.00 | 5155.67 | 1.00 | 1.00 | 1.00 | 1.00 | 206.25 |
| G0099 | 85.00 | 251.00 | 35.67 | 11991.33 | 1.00 | 108.00 | 1.00 | 1.00 | 297.91 |
| G0100 | 24.67 | 1.00 | 1.00 | 1.00 | 1.00 | 1.00 | 1.00 | 1.00 | 74.64 |
| N010 | 1978.67 | 1.00 | 1.00 | 410.00 | 1.00 | 1.00 | 1.00 | 1.00 | 82.80 |

| | | | | | | | | | |
|---|---|---|---|---|---|---|---|---|---|
| N011 | 168.00 | 1.00 | 1.00 | 1.00 | 1.00 | 7.00 | 17.00 | 1.00 | 295.70 |
| N012 | 1.00 | 1.00 | 1.00 | 1.00 | 1.00 | 1.00 | 1.00 | 1.00 | 167.08 |
| N013 | 555.33 | 278.33 | 1.00 | 1.00 | 1.00 | 1.00 | 1.00 | 1.00 | 90.19 |
| N014 | 1253.67 | 1.00 | 1.00 | 1.00 | 2.33 | 1.00 | 1.00 | 1.00 | 125.69 |
| N015 | 456.33 | 402.67 | 1.00 | 1.00 | 26.67 | 28.00 | 1.00 | 1.00 | 156.70 |
| N020 | 3466.00 | 566.00 | 113.67 | 1.00 | 1.00 | 61.00 | 1.00 | 1.00 | 199.61 |
| N021 | 8.33 | 19.67 | 1.00 | 1.00 | 28.00 | 122.00 | 1.00 | 1.00 | 167.09 |
| N022 | 1.00 | 1.00 | 1.00 | 273.33 | 1.00 | 35.00 | 1.00 | 1.00 | 786.56 |
| N023 | 1.00 | 1.00 | 1.00 | 668.67 | 1.00 | 1.00 | 1.00 | 1.00 | 124.18 |
| N024 | 71.00 | 1.00 | 1.00 | 1.00 | 1.00 | 1.00 | 1.00 | 1.00 | 91.66 |
| N025 | 111.67 | 80.67 | 1.00 | 1.00 | 1.00 | 1.00 | 1.00 | 1.00 | 215.87 |
| N026 | 59.33 | 22.33 | 1.00 | 1.00 | 1.00 | 130.00 | 61.00 | 1.00 | 96.85 |
| N022G | 71.00 | 146.00 | 1.00 | 1.00 | 1.00 | 1.00 | 1.00 | 1.00 | 127.17 |
| N023G | 110.00 | 161.33 | 1.00 | 1.00 | 1.00 | 49.00 | 1.00 | 1.00 | 170.03 |
| N025G | 68.67 | 16.00 | 1.00 | 1.00 | 1.00 | 51.00 | 1.00 | 1.00 | 113.85 |
| N030 | 1772.00 | 103.67 | 1.00 | 1.00 | 1.00 | 1.00 | 1.00 | 1.00 | 372.59 |
| N210 | 1474.33 | 1.00 | 1.00 | 542.00 | 39.00 | 1.00 | 1.00 | 1.00 | 361.50 |
| N040 | 663.33 | 1.00 | 97.33 | 1.00 | 1.00 | 1.00 | 1.00 | 1.00 | 148.59 |
| N041 | 3820.67 | 612.00 | 1.00 | 1.00 | 1.33 | 3.00 | 1.00 | 1.00 | 236.55 |
| N042 | 58.33 | 1.00 | 2.33 | 1.00 | 9.67 | 1.00 | 1.00 | 1.00 | 239.52 |
| N043 | 206.00 | 734.67 | 1.00 | 1.00 | 1.00 | 1.00 | 1.00 | 1.00 | 227.70 |
| N044 | 744.00 | 653.00 | 1.00 | 1.00 | 5.00 | 1.00 | 1.00 | 1.00 | 184.81 |
| N045 | 295.67 | 128.67 | 1.00 | 1.00 | 1.00 | 38.00 | 1.00 | 1.00 | 207.00 |
| N050 | 2902.67 | 171.00 | 1.00 | 1.00 | 11.67 | 38.00 | 1.00 | 1.00 | 373.33 |
| N051 | 347.67 | 1.00 | 1.00 | 1.00 | 8.33 | 2.00 | 1.00 | 1.00 | 360.75 |
| N052 | 63.00 | 1.00 | 1.00 | 1.00 | 1.00 | 5.00 | 1.00 | 1.00 | 347.45 |
| N053 | 82.33 | 533.00 | 1.00 | 1.00 | 1.00 | 1.00 | 1.00 | 1.00 | 87.22 |
| N054 | 810.00 | 1.00 | 1.00 | 1.00 | 2.00 | 1.00 | 1.00 | 1.00 | 263.17 |
| N055 | 814.00 | 498.33 | 1.00 | 1.00 | 1.00 | 1.00 | 1.00 | 1.00 | 516.00 |
| TE001 | 187.67 | 233.67 | 1.00 | 10696.67 | 27.33 | 1.00 | 1.00 | 1.00 | 167.06 |
| TE002 | 46.00 | 1.00 | 1.00 | 1.00 | 10.33 | 73.00 | 71.00 | 1.00 | 193.70 |
| TE003 | 26.67 | 106.33 | 1.00 | 1.00 | 2.33 | 69.00 | 1.00 | 1.00 | 439.13 |
| TE004 | 82.67 | 1.00 | 1.00 | 1.00 | 1.00 | 167.00 | 1.00 | 1.00 | 338.57 |
| TE005 | 240.67 | 692.00 | 1.00 | 1.00 | 1.00 | 132.00 | 1.00 | 1.00 | 805.79 |
| TE006 | 775.00 | 415.33 | 1.00 | 1.00 | 5.67 | 1.00 | 1.00 | 1.00 | 510.06 |

| | | | | | | | | | |
|---|---|---|---|---|---|---|---|---|---|
| TE007 | 86.33 | 101.00 | 1.00 | 1.00 | 8.00 | 166.00 | 6.00 | 1.00 | 575.14 |
| TE008 | 251.33 | 41.00 | 1.00 | 1.00 | 1.00 | 1.00 | 1.00 | 1.00 | 306.05 |
| TE009 | 40.33 | 1.00 | 1.00 | 1.00 | 1.00 | 1.00 | 1.00 | 1.00 | 189.25 |
| TE010 | 1.00 | 1.00 | 1.00 | 1.00 | 1.00 | 1.00 | 1.00 | 1.00 | 510.07 |
| TE011 | 62.00 | 149.67 | 1.00 | 1.00 | 1.00 | 1.00 | 7.00 | 1.00 | 221.77 |
| TE012 | 48.33 | 43.00 | 1.00 | 1.00 | 1.00 | 88.00 | 1.00 | 1.00 | 411.02 |
| TE013 | 77.67 | 231.67 | 1.00 | 1.00 | 4.00 | 172.00 | 1.00 | 1.00 | 204.76 |
| TE014 | 200.67 | 486.67 | 1.00 | 1.00 | 1.00 | 169.00 | 1.00 | 1.00 | 73.19 |
| TE015 | 117.67 | 1.00 | 1.00 | 1.00 | 5.67 | 167.00 | 1.00 | 1.00 | 320.85 |
| TE016 | 168.00 | 1.00 | 1.00 | 1.00 | 1.00 | 132.00 | 1.00 | 1.00 | 379.99 |
| TE017 | 1.00 | 1.00 | 1.00 | 1.00 | 11.67 | 50.00 | 13.00 | 1.00 | 564.05 |
| TE018 | 21.67 | 1.00 | 1.00 | 1.00 | 1.00 | 1.00 | 1.00 | 1.00 | 329.71 |
| TE019 | 22.00 | 33.67 | 1.00 | 1.00 | 1.00 | 96.00 | 1.00 | 1.00 | 255.78 |
| TE020 | 25.33 | 1.00 | 1.00 | 1.00 | 5.00 | 165.00 | 1.00 | 1.00 | 171.50 |
| TE021 | 49.67 | 109.67 | 1.00 | 1.00 | 12.67 | 58.00 | 1.00 | 1.00 | 200.35 |
| TE022 | 139.33 | 260.00 | 1.00 | 1.00 | 1.00 | 12.00 | 1.00 | 1.00 | 430.24 |
| TE023 | 71.00 | 284.67 | 1.00 | 1.00 | 1.00 | 142.00 | 1.00 | 1.00 | 437.65 |
| TE024 | 1665.00 | 1.00 | 1.00 | 1.00 | 1.00 | 167.00 | 1.00 | 1.00 | 419.91 |
| TE025 | 5.00 | 5.00 | 3.33 | 1.00 | 4.00 | 297.00 | 1.00 | 1.00 | 134.55 |
| TE026 | 3126.00 | 1.00 | 1.00 | 1.00 | 1.00 | 346.00 | 36.00 | 1.00 | 131.60 |
| TE027 | 3.33 | 53.67 | 1.00 | 1.00 | 1.67 | 252.00 | 53.00 | 1.00 | 181.87 |
| TE028 | 1328.00 | 1.00 | 36.33 | 1.00 | 1.00 | 1262.00 | 107.00 | 1.00 | 298.68 |
| TE029 | 45.33 | 63.00 | 1.00 | 1.00 | 5.00 | 376.00 | 44.00 | 1.00 | 218.81 |
| TE030 | 2042.67 | 397.33 | 1.00 | 1.00 | 1.00 | 556.00 | 113.00 | 1.00 | 275.00 |
| TE031 | 113.00 | 1.00 | 1.00 | 1.00 | 15.33 | 607.00 | 237.00 | 1.00 | 490.86 |
| TE032 | 1055.00 | 1.00 | 1.00 | 1.00 | 4.00 | 1.00 | 1.00 | 1.00 | 506.41 |
| TE033 | 1.00 | 1.00 | 1.00 | 1.00 | 1.00 | 118.00 | 1.00 | 1.00 | 294.24 |
| TE034 | 6.67 | 1.00 | 1.00 | 1.00 | 1.00 | 154.00 | 1.00 | 1.00 | 110.87 |
| TE035 | 3574.33 | 469.67 | 32.00 | 1.00 | 1.00 | 166.00 | 1.00 | 1.00 | 147.10 |
| TE036 | 1.00 | 12.67 | 1.00 | 1.00 | 1.00 | 302.00 | 57.00 | 1.00 | 201.08 |
| TE037 | 5100.67 | 854.67 | 1.00 | 1.00 | 1.00 | 225.00 | 9.00 | 1.00 | 266.14 |
| TE038 | 124.67 | 166.00 | 1.00 | 1.00 | 12.00 | 392.00 | 1.00 | 1.00 | 255.80 |
| TE039 | 304.33 | 137.00 | 1.00 | 1.00 | 1.00 | 1.00 | 1.00 | 1.00 | 127.13 |
| TE040 | 1453.00 | 59.67 | 1.00 | 1.00 | 1.00 | 1.00 | 1.00 | 1.00 | 177.40 |
| TE041 | 2.00 | 44.67 | 1.00 | 1.00 | 1.00 | 99.00 | 4.00 | 1.00 | 152.30 |

| | | | | | | | | | |
|---|---|---|---|---|---|---|---|---|---|
| TE042 | 289.33 | 180.33 | 1.00 | 1.00 | 7.00 | 1.00 | 1.00 | 1.00 | 144.17 |
| TE043 | 78.33 | 141.00 | 3.00 | 1.00 | 1.00 | 334.00 | 141.00 | 1.00 | 61.34 |
| TE044 | 12.33 | 47.00 | 22.00 | 1.00 | 1.00 | 214.00 | 92.00 | 1.00 | 238.77 |
| TE045 | 443.00 | 254.33 | 1.00 | 1.00 | 1.00 | 1.00 | 1.00 | 1.00 | 337.10 |
| TE046 | 29.67 | 243.00 | 1.00 | 1.00 | 1.00 | 496.00 | 1.00 | 1.00 | 354.10 |
| TE047 | 7.33 | 1.00 | 19.00 | 1.00 | 19.67 | 530.00 | 322.00 | 1.00 | 323.81 |
| TE048 | 17.33 | 90.33 | 1.00 | 1.00 | 1.00 | 253.00 | 121.00 | 1.00 | 175.94 |
| TE049 | 146.00 | 42.00 | 1.00 | 1.00 | 4.67 | 1.00 | 1.00 | 1.00 | 251.33 |
| TE050 | 301.00 | 418.67 | 1.00 | 1.00 | 19.33 | 437.00 | 199.00 | 1.00 | 270.56 |
| H0100 | 3117.67 | 667.67 | 101.00 | 4.00 | 1.00 | 1.00 | 1.00 | 2226.67 | 801.35 |
| H0101 | 170.00 | 251.67 | 3.00 | 1.00 | 1.00 | 124.00 | 230.00 | 1.00 | 781.39 |
| H0103 | 102.33 | 44.33 | 1.00 | 122.00 | 1.00 | 1.00 | 1.00 | 1.00 | 780.66 |
| H0105 | 1079.67 | 1.00 | 31.33 | 1.00 | 2.33 | 137.00 | 104.00 | 1.00 | 390.34 |
| H0106 | 763.00 | 1.00 | 1.00 | 1.00 | 1.33 | 1.00 | 1.00 | 1.00 | 252.83 |
| H0200 | 400.33 | 322.00 | 24.00 | 1.00 | 1.00 | 116.00 | 56.00 | 1.00 | 217.35 |
| H0201 | 56.00 | 52.33 | 1.00 | 694.00 | 9.33 | 14.00 | 1.00 | 1.00 | 647.59 |
| H0202 | 188.67 | 260.00 | 1.00 | 531.33 | 1.00 | 1.00 | 1.00 | 1.00 | 601.75 |
| H0203 | 562.67 | 503.33 | 1.00 | 942.67 | 1.00 | 1.00 | 1.00 | 1.00 | 796.91 |
| H0204 | 1827.00 | 536.67 | 98.00 | 1.00 | 3.00 | 245.00 | 333.00 | 1.00 | 674.94 |
| H0205 | 1.00 | 37.33 | 5.67 | 1.00 | 1.00 | 1.00 | 1.00 | 1.00 | 630.58 |
| H0207 | 63.33 | 1.00 | 1.00 | 1.00 | 1.00 | 1.00 | 84.00 | 1.00 | 564.79 |
| H0208 | 531.67 | 1.00 | 1.00 | 1.00 | 1.00 | 2387.00 | 369.00 | 1.00 | 579.58 |
| H0209 | 2145.00 | 616.67 | 118.00 | 100.00 | 3.00 | 1.00 | 1.00 | 685.33 | 515.28 |
| H0210 | 916.00 | 129.00 | 1.00 | 707.33 | 1.00 | 1.00 | 106.00 | 1.00 | 458.35 |
| H0300 | 3813.67 | 595.67 | 1.00 | 1.00 | 1.00 | 12.00 | 6.00 | 1.00 | 263.19 |
| H0301 | 569.33 | 591.00 | 1.00 | 1.00 | 1.00 | 1.00 | 1.00 | 1.00 | 814.66 |
| H0303 | 1613.67 | 618.67 | 60.33 | 1.00 | 1.00 | 1.00 | 11.00 | 1.00 | 683.81 |
| H0304 | 2068.67 | 1.00 | 30.33 | 1.00 | 1.00 | 90.00 | 30.00 | 1.00 | 625.42 |
| H0305 | 38.00 | 1.00 | 1.00 | 1.00 | 1.00 | 61.00 | 7.00 | 1.00 | 604.72 |
| H0306 | 61.00 | 1.00 | 1.00 | 1.00 | 1.00 | 21.00 | 97.00 | 1.00 | 737.79 |
| H0307 | 85.67 | 366.00 | 6.67 | 1.00 | 6.00 | 1.00 | 98.00 | 1.00 | 652.76 |
| H0400 | 111.00 | 1.00 | 3.67 | 1.00 | 3.67 | 10.00 | 54.00 | 3597.33 | 375.54 |
| H0402 | 330.33 | 98.67 | 11.67 | 1.00 | 1.00 | 43.00 | 19.00 | 5055.33 | 345.98 |
| H0403 | 247.00 | 207.00 | 27.67 | 1.00 | 1.00 | 14.00 | 1.00 | 1.00 | 611.36 |
| H0404 | 348.33 | 289.67 | 50.67 | 1.00 | 3.00 | 1.00 | 19.00 | 1.00 | 717.09 |

| | | | | | | | | | |
|---|---|---|---|---|---|---|---|---|---|
| H0405 | 220.33 | 81.67 | 1.00 | 266.67 | 1.00 | 6.00 | 1.00 | 1.00 | 645.37 |
| H0406 | 156.00 | 1.00 | 1.00 | 1.00 | 1.00 | 1.00 | 1.00 | 1.00 | 636.51 |
| H0407 | 129.67 | 113.33 | 1.00 | 1.00 | 1.00 | 133.00 | 1.00 | 1.00 | 637.26 |
| H0408 | 252.67 | 417.33 | 1.00 | 1.00 | 1.67 | 69.00 | 1.00 | 1.00 | 802.84 |
| H0409 | 246.67 | 114.00 | 1.00 | 9313.67 | 1.00 | 764.00 | 1.00 | 3597.33 | 606.18 |
| H0410 | 139.33 | 307.67 | 31.67 | 1.00 | 6.00 | 121.00 | 1.00 | 1.00 | 712.66 |
| H0411 | 174.33 | 214.00 | 421.33 | 1.00 | 1.00 | 1.00 | 1.00 | 1.00 | 246.91 |
| H0500 | 185.67 | 206.00 | 19.33 | 1.00 | 1.00 | 1.00 | 1.00 | 1.00 | 705.26 |
| H0503 | 194.33 | 80.00 | 1.00 | 141.33 | 1.00 | 1.00 | 1.00 | 1.00 | 561.84 |
| H0504 | 168.67 | 1.00 | 2.00 | 1.00 | 1.00 | 1.00 | 1.00 | 1.00 | 805.05 |
| H0505 | 50.67 | 1.00 | 1.00 | 1.00 | 1.00 | 1.00 | 53.00 | 1.00 | 621.72 |
| H0600 | 81.00 | 109.67 | 1.00 | 1.00 | 13.67 | 153.00 | 242.00 | 1.00 | 436.91 |
| H0601 | 229.67 | 101.33 | 1.00 | 1.00 | 1.00 | 106.00 | 132.00 | 1.00 | 405.13 |
| H0602 | 1266.33 | 750.67 | 171.67 | 1.00 | 1.00 | 132.00 | 26.00 | 1.00 | 552.97 |
| H0603 | 2926.67 | 260.33 | 23.33 | 1.00 | 2.00 | 168.00 | 137.00 | 1.00 | 643.16 |
| H0604 | 59.00 | 25.00 | 1.00 | 44.33 | 1.00 | 41.00 | 4.00 | 1.00 | 582.52 |
| H0606 | 144.67 | 219.67 | 80.33 | 1.00 | 1.00 | 9.00 | 1.00 | 1.00 | 629.87 |
| H0608 | 403.67 | 144.33 | 63.33 | 1.00 | 1.00 | 1.00 | 1.00 | 342.67 | 743.70 |
| H0609 | 218.33 | 255.67 | 13.67 | 1.00 | 1.00 | 1.00 | 15.00 | 1.00 | 634.29 |
| H0610 | 1075.33 | 480.67 | 55.67 | 1.00 | 1.00 | 1.00 | 1.00 | 1.00 | 533.76 |
| H0700 | 109.33 | 1.00 | 1.00 | 1.00 | 107.33 | 417.00 | 523.00 | 1.00 | 491.60 |
| H0701 | 2371.00 | 875.67 | 76.00 | 1.00 | 1.00 | 237.00 | 22.00 | 1.00 | 586.97 |
| H0800 | 118.67 | 1.00 | 1.00 | 1.00 | 1.00 | 481.00 | 137.00 | 1.00 | 505.66 |
| H0900 | 306.00 | 1.00 | 1.00 | 1.00 | 1.00 | 145.00 | 147.00 | 1.00 | 710.42 |
| L1001 | 96.00 | 215.33 | 1.00 | 1.00 | 1.00 | 1.00 | 1.00 | 1.00 | 711.90 |
| L1002 | 183.00 | 1.00 | 1.00 | 276.67 | 1.00 | 51.00 | 1.00 | 1.00 | 776.23 |
| L1003 | 184.67 | 351.33 | 1.00 | 1.00 | 1.00 | 6.00 | 16.00 | 1.00 | 462.79 |
| L1011 | 183.00 | 178.33 | 50.33 | 1.00 | 1.00 | 1.00 | 1.00 | 1028.00 | 591.43 |
| L1012 | 74.67 | 1.00 | 1.00 | 1.00 | 1.00 | 1.00 | 1.00 | 1.00 | 384.41 |
| L1013 | 273.33 | 375.00 | 74.33 | 1.00 | 1.00 | 1.00 | 1.00 | 1.00 | 152.29 |
| L1021 | 1.00 | 1.00 | 1.00 | 1.00 | 1.00 | 1.00 | 80.00 | 1.00 | 245.43 |
| L1022 | 110.00 | 1.00 | 1.00 | 1491.67 | 1.00 | 1.00 | 380.00 | 1.00 | 775.46 |
| L1023 | 18.33 | 80.67 | 1.00 | 1.00 | 1.00 | 1.00 | 59.00 | 1.00 | 493.83 |
| L1201 | 50.00 | 1.00 | 17.00 | 1.00 | 1.00 | 1.00 | 1.00 | 1.00 | 628.38 |
| L1202 | 137.33 | 144.67 | 25.67 | 1.00 | 1.00 | 27.00 | 1.00 | 1.00 | 626.88 |

| | | | | | | | | | |
|---|---|---|---|---|---|---|---|---|---|
| L1203 | 105.33 | 133.33 | 24.67 | 1.00 | 2.67 | 1.00 | 1.00 | 1.00 | 549.26 |
| L1204 | 110.00 | 1.67 | 1.00 | 1.00 | 1.00 | 1.00 | 1.00 | 1.00 | 705.98 |
| L1205 | 230.33 | 411.00 | 36.00 | 1.00 | 1.00 | 3.00 | 1.00 | 1.00 | 534.46 |
| L1206 | 128.67 | 1.00 | 1.00 | 1.00 | 1.00 | 3.00 | 1.00 | 1.00 | 245.43 |
| L1207 | 106.00 | 1.00 | 1.00 | 1.00 | 1.00 | 7.00 | 1.00 | 1.00 | 258.02 |
| L1209 | 74.00 | 7.33 | 1.00 | 1.00 | 1.00 | 1.00 | 1.00 | 1.00 | 445.06 |
| L1211 | 22.00 | 1.00 | 1.00 | 314.33 | 1.00 | 1.00 | 1.00 | 1.00 | 617.29 |
| L1212 | 67.33 | 1.00 | 70.00 | 1.00 | 1.00 | 1.00 | 1.00 | 1.00 | 405.87 |
| L1213 | 26.00 | 1.00 | 15.33 | 1.00 | 1.00 | 1.00 | 1.00 | 1.00 | 476.81 |
| L1214 | 58.00 | 1.00 | 1.00 | 1.00 | 1.00 | 1.00 | 1.00 | 1.00 | 628.36 |
| L1215 | 132.33 | 140.00 | 1.00 | 1.00 | 1.00 | 1.00 | 1.00 | 1.00 | 444.28 |
| L1216 | 65.67 | 1.00 | 1.00 | 1.00 | 1.00 | 1.00 | 1.00 | 1.00 | 422.85 |
| L1221 | 105.67 | 1.00 | 1.00 | 1.00 | 1.00 | 1.00 | 68.00 | 1.00 | 674.20 |
| L1222 | 85.67 | 49.00 | 2.67 | 1.00 | 1.00 | 1.00 | 45.00 | 1.00 | 416.93 |
| L1225 | 53.00 | 1.00 | 22.67 | 1.00 | 1.00 | 1.00 | 41.00 | 1.00 | 440.60 |
| L1226 | 129.67 | 118.00 | 1.00 | 1.00 | 1.00 | 1.00 | 1.00 | 1.00 | 378.51 |
| L2000 | 1535.33 | 481.67 | 54.00 | 1.00 | 1.00 | 148.00 | 1.00 | 1.00 | 192.22 |
| L2100 | 50.67 | 12.67 | 1.00 | 1.00 | 1.00 | 182.00 | 1.00 | 1.00 | 555.91 |
| L2101 | 69.33 | 144.67 | 42.00 | 1.00 | 18.67 | 206.00 | 8.00 | 1.00 | 301.60 |
| L2102 | 149.67 | 1.00 | 1.00 | 843.33 | 1.00 | 148.00 | 1.00 | 1.00 | 616.52 |
| L2111 | 259.00 | 1.00 | 1.00 | 1.00 | 1.00 | 65.00 | 1.00 | 170.67 | 360.02 |
| L2112 | 34.00 | 129.33 | 1.00 | 1.00 | 1.00 | 152.00 | 1.00 | 1.00 | 353.36 |
| L2113 | 25.67 | 1.00 | 1.00 | 1.00 | 1.00 | 179.00 | 1.00 | 1.00 | 248.40 |
| L2121 | 76.00 | 135.67 | 1.00 | 1.00 | 1.00 | 65.00 | 1.00 | 1.00 | 630.58 |
| L2122 | 103.67 | 135.00 | 1.00 | 1.00 | 1.00 | 52.00 | 1.00 | 1.00 | 448.00 |
| L2103 | 103.33 | 1.00 | 1.00 | 1.00 | 1.00 | 189.00 | 80.00 | 641.33 | 401.42 |
| L2104 | 409.33 | 303.33 | 52.00 | 1.00 | 1.00 | 140.00 | 1.00 | 1.00 | 339.32 |
| L2131 | 42.00 | 77.33 | 42.67 | 1.00 | 1.00 | 1.00 | 1.00 | 1.00 | 502.70 |
| L2132 | 132.33 | 145.00 | 32.33 | 1.00 | 1.00 | 1.00 | 1.00 | 1.00 | 250.60 |
| L2133 | 112.33 | 187.00 | 11.67 | 1.00 | 1.00 | 1.00 | 1.00 | 1.00 | 192.21 |
| L2191 | 392.67 | 1.00 | 43.67 | 1.00 | 1.00 | 269.00 | 1.00 | 1.00 | 310.49 |
| L2192 | 70.33 | 1.00 | 44.00 | 1.00 | 1.00 | 144.00 | 6.00 | 1.00 | 403.64 |
| L2200 | 2607.33 | 211.67 | 112.00 | 1.00 | 1.00 | 207.00 | 20.00 | 1.00 | 379.24 |
| L2300 | 6.00 | 46.00 | 1.00 | 1.00 | 1.00 | 361.00 | 43.00 | 1.00 | 728.91 |
| L2301 | 16.00 | 1.00 | 1.00 | 1.00 | 1.00 | 340.00 | 76.00 | 1.00 | 236.57 |

| | | | | | | | | | |
|---|---|---|---|---|---|---|---|---|---|
| L2302 | 40.67 | 39.00 | 35.67 | 1.00 | 1.00 | 214.00 | 1.00 | 1.00 | 320.84 |
| L2311 | 1.00 | 15.00 | 4.33 | 1.00 | 26.67 | 141.00 | 1.00 | 1.00 | 277.98 |
| L2312 | 1.00 | 19.00 | 30.33 | 127.33 | 1.00 | 112.00 | 1.00 | 1.00 | 466.46 |
| L2303 | 55.00 | 1.00 | 1.00 | 1.00 | 16.00 | 173.00 | 1.00 | 1.00 | 331.20 |
| L2304 | 240.67 | 63.67 | 48.67 | 359.33 | 1.00 | 1.00 | 1.00 | 1.00 | 357.06 |
| L2391 | 1880.33 | 263.67 | 136.33 | 85.33 | 1.00 | 1.00 | 1.00 | 1.00 | 311.99 |
| L2392 | 1.00 | 22.67 | 79.67 | 111.33 | 1.00 | 448.00 | 74.00 | 1.00 | 575.16 |
| L2900 | 62.67 | 1.33 | 1.00 | 1.00 | 1.00 | 141.00 | 242.00 | 1.00 | 523.42 |
| L2911 | 49.67 | 56.00 | 1.00 | 1.00 | 1.00 | 154.00 | 1.00 | 1.00 | 523.40 |
| L2912 | 50.33 | 74.67 | 45.00 | 1.00 | 1.00 | 112.00 | 13.00 | 1.00 | 429.52 |
| L2913 | 77.00 | 136.00 | 16.00 | 1.00 | 1.00 | 66.00 | 1.00 | 1.00 | 285.36 |
| L3100 | 108.67 | 1.00 | 34.67 | 1.00 | 1.00 | 1.00 | 1.00 | 1.00 | 498.25 |
| L3101 | 12.67 | 1.00 | 40.67 | 1.00 | 1.00 | 1.00 | 46.00 | 1.00 | 337.82 |
| L3102 | 21.67 | 98.67 | 1.00 | 1.00 | 1.00 | 1.00 | 73.00 | 1.00 | 263.17 |
| L3111 | 1.00 | 30.33 | 1.33 | 1.00 | 4.67 | 1.00 | 90.00 | 1.00 | 192.22 |
| L3112 | 44.33 | 57.67 | 91.33 | 1.00 | 1.00 | 85.00 | 158.00 | 1.00 | 396.26 |
| L3113 | 72.33 | 27.67 | 25.33 | 1.00 | 1.00 | 82.00 | 93.00 | 1.00 | 316.43 |
| L3103 | 58.67 | 284.00 | 41.67 | 1.00 | 3.67 | 44.00 | 44.00 | 1.00 | 450.96 |
| L3200 | 53.33 | 100.00 | 47.33 | 1.00 | 1.00 | 87.00 | 198.00 | 86.00 | 470.16 |
| L3201 | 55.67 | 32.33 | 1.00 | 1.00 | 3.00 | 1.00 | 1.00 | 1.00 | 325.28 |
| L3202 | 7.00 | 5.33 | 11.33 | 1.00 | 1.00 | 14.00 | 53.00 | 1.00 | 532.99 |
| L3211 | 25.33 | 62.67 | 14.00 | 1.00 | 1.00 | 1.00 | 487.00 | 1.00 | 687.50 |
| L3212 | 18.33 | 1.00 | 1.00 | 1.00 | 1.00 | 70.00 | 121.00 | 1.00 | 612.09 |
| L3213 | 42.00 | 13.33 | 1.00 | 1.00 | 9.00 | 74.00 | 192.00 | 1.00 | 148.60 |

SI-Spreadsheet 5

### Glycan 300 Structure Symbols

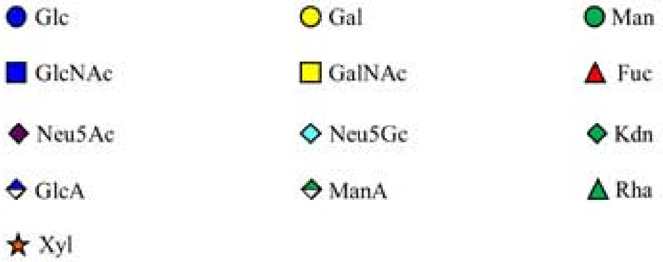


### Glycan 100 Glycans

#### A. Linkers

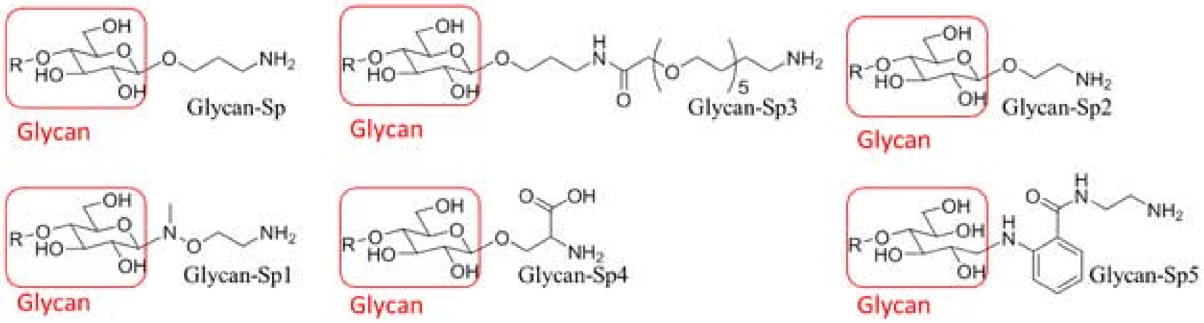


#### B. Glycan 100 Structures

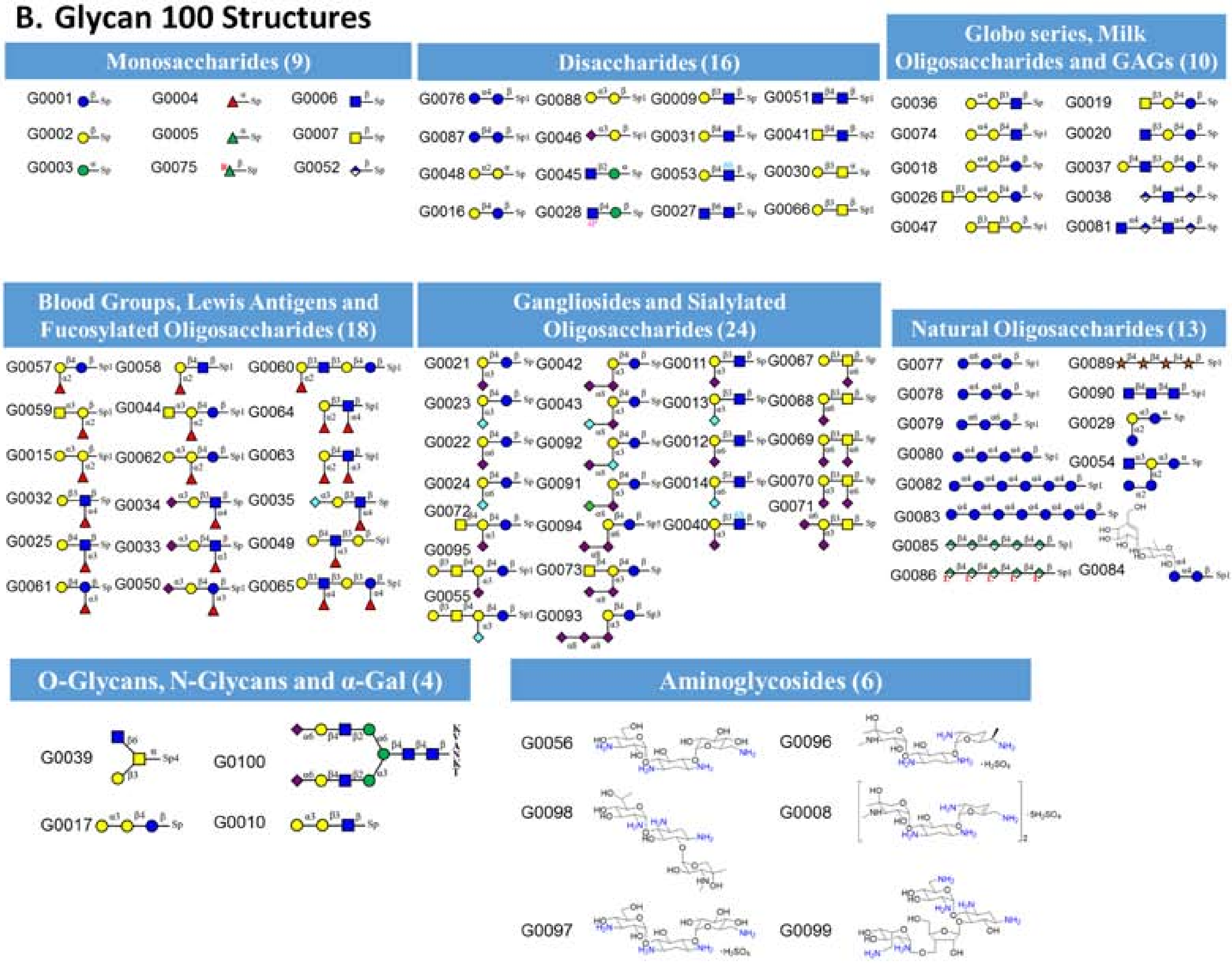

### N-Glycans

#### A. Linkers

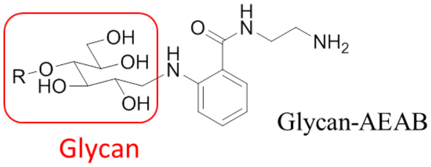


#### B. N-Glycan Structures

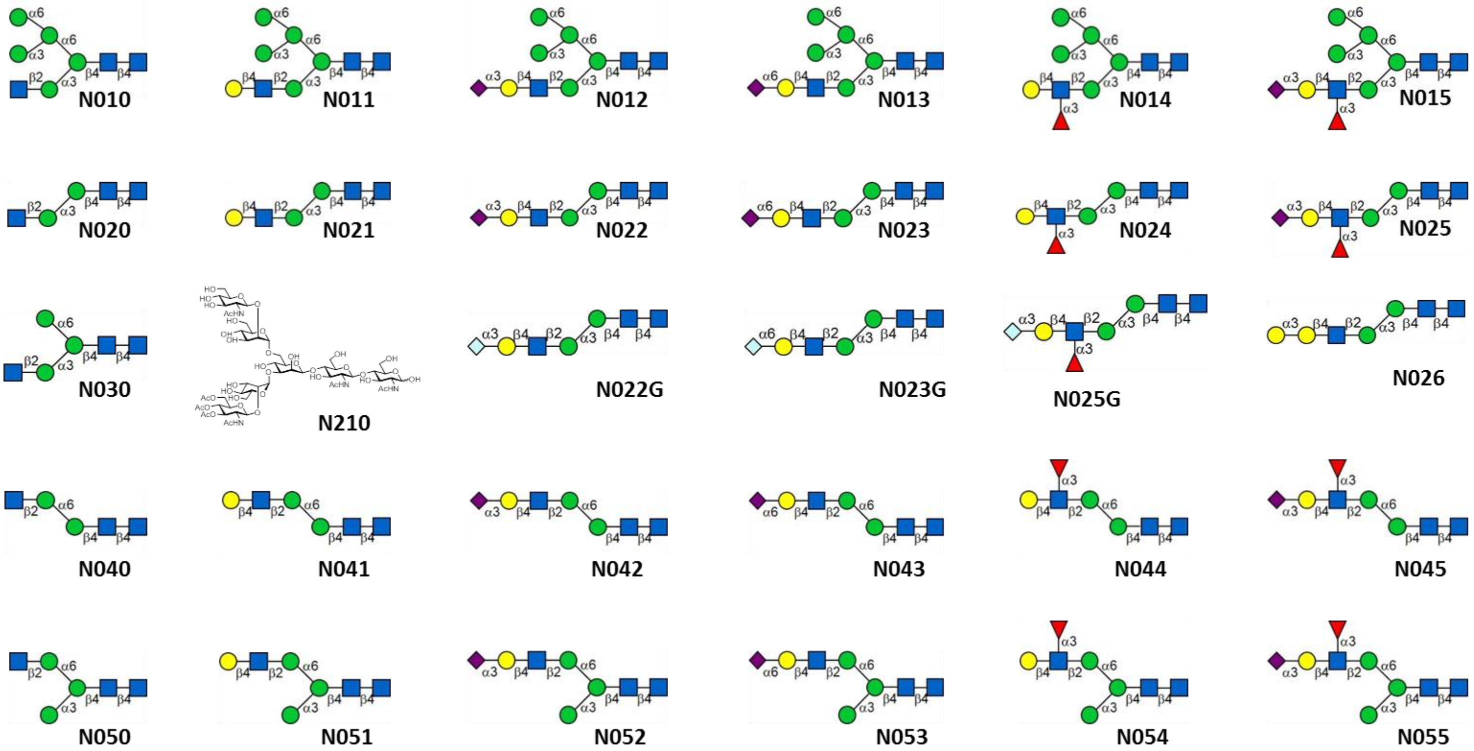

### Glycolipid Glycans

#### A. Linkers

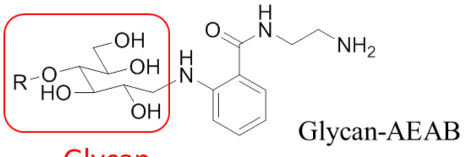


#### B. Glycolipid Glycan Structures

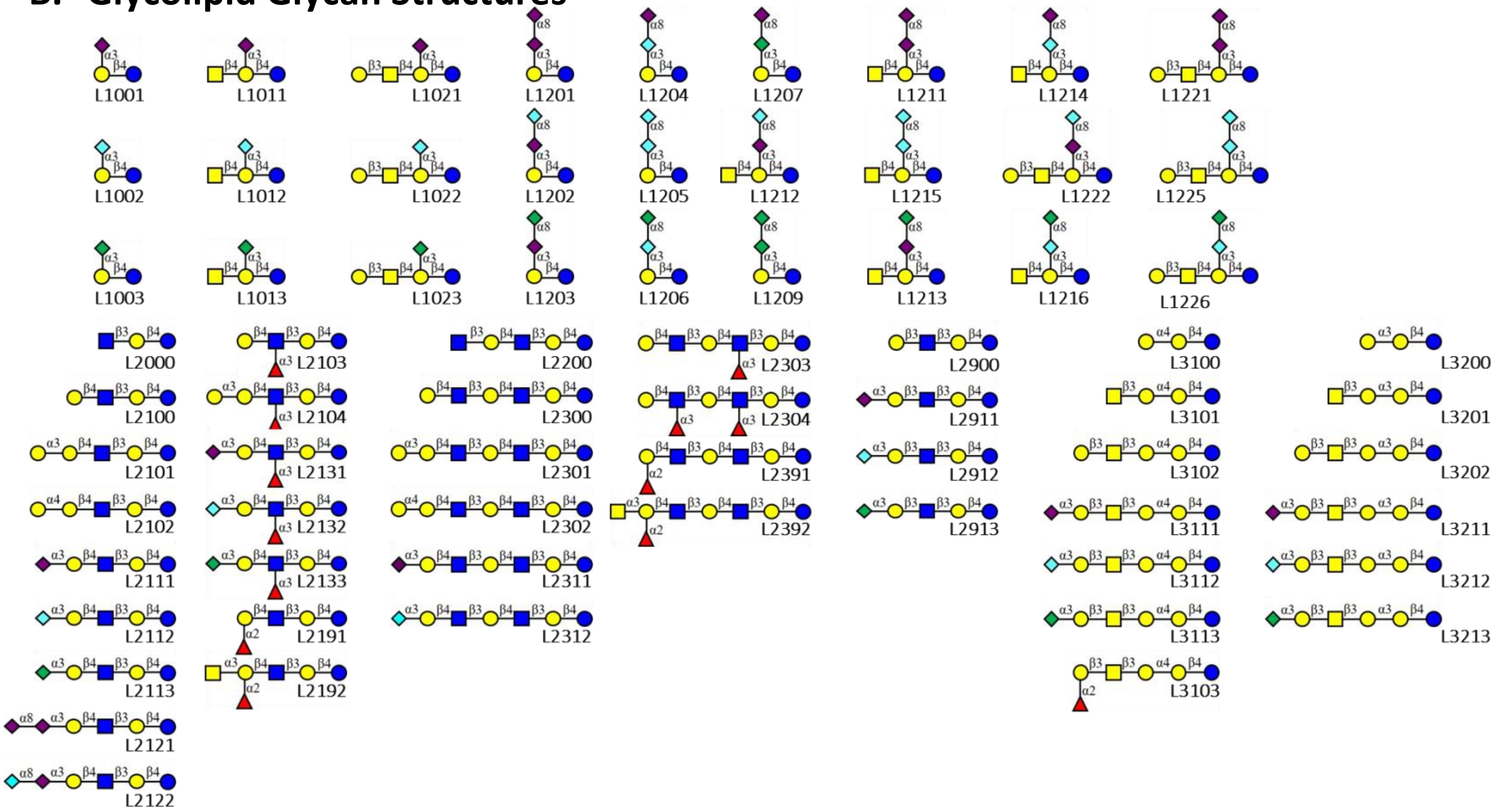

### Human Milk Oligosaccharides

#### A. Linkers

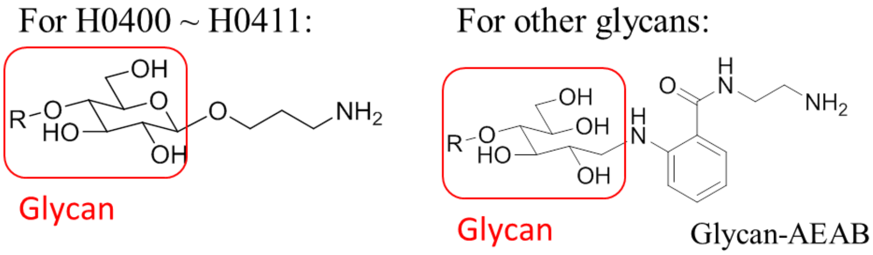


#### B. Human Milk Oligosaccharide Structures

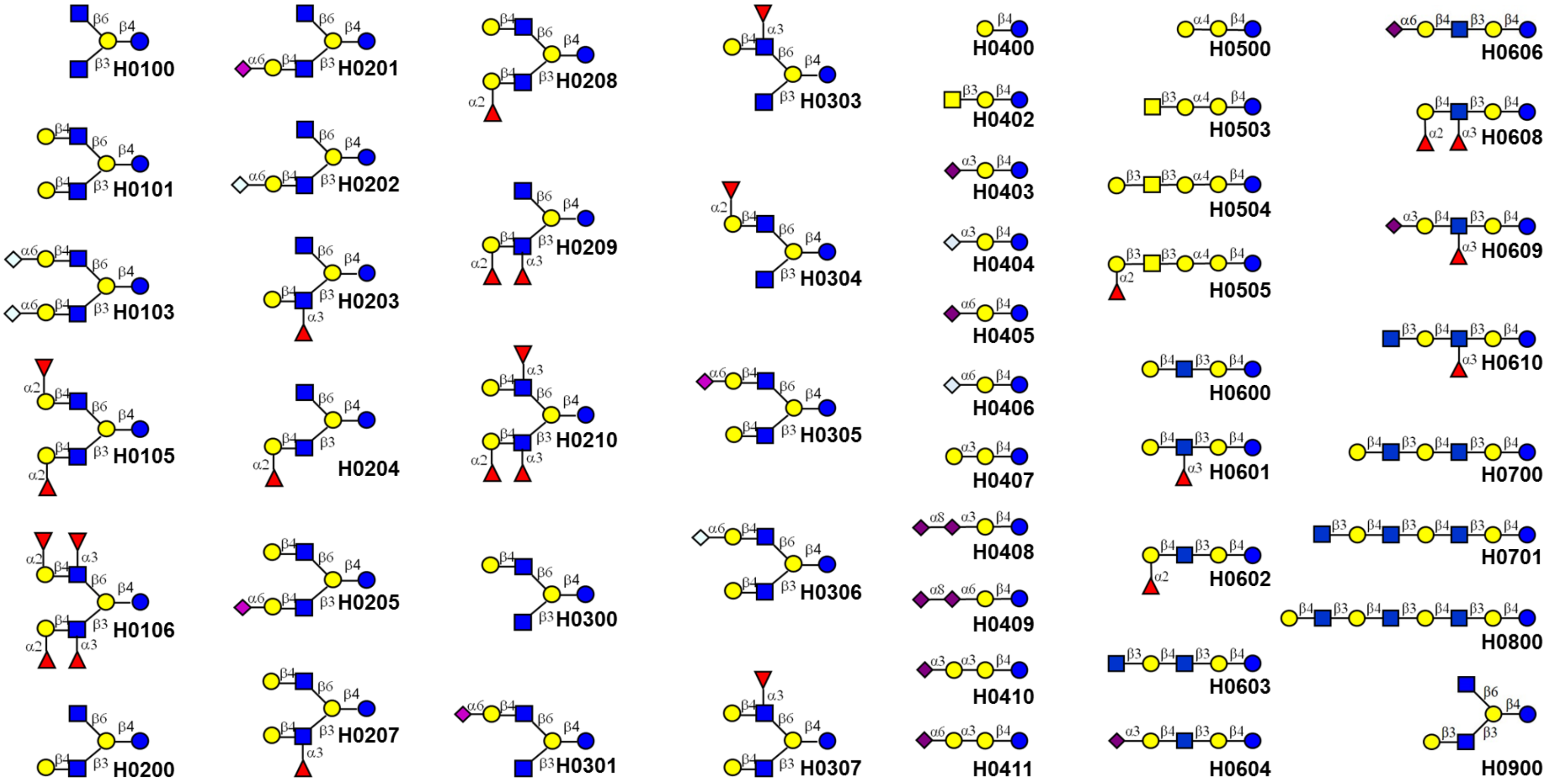

#### Tandem Epitopes

##### A. Linkers

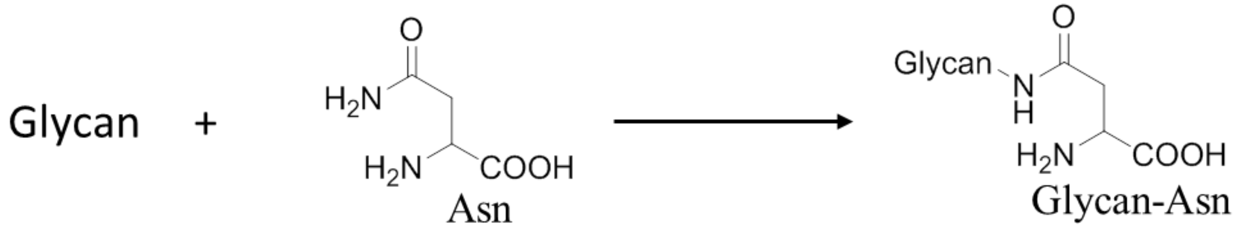


##### B. Tandem Epitope Structures

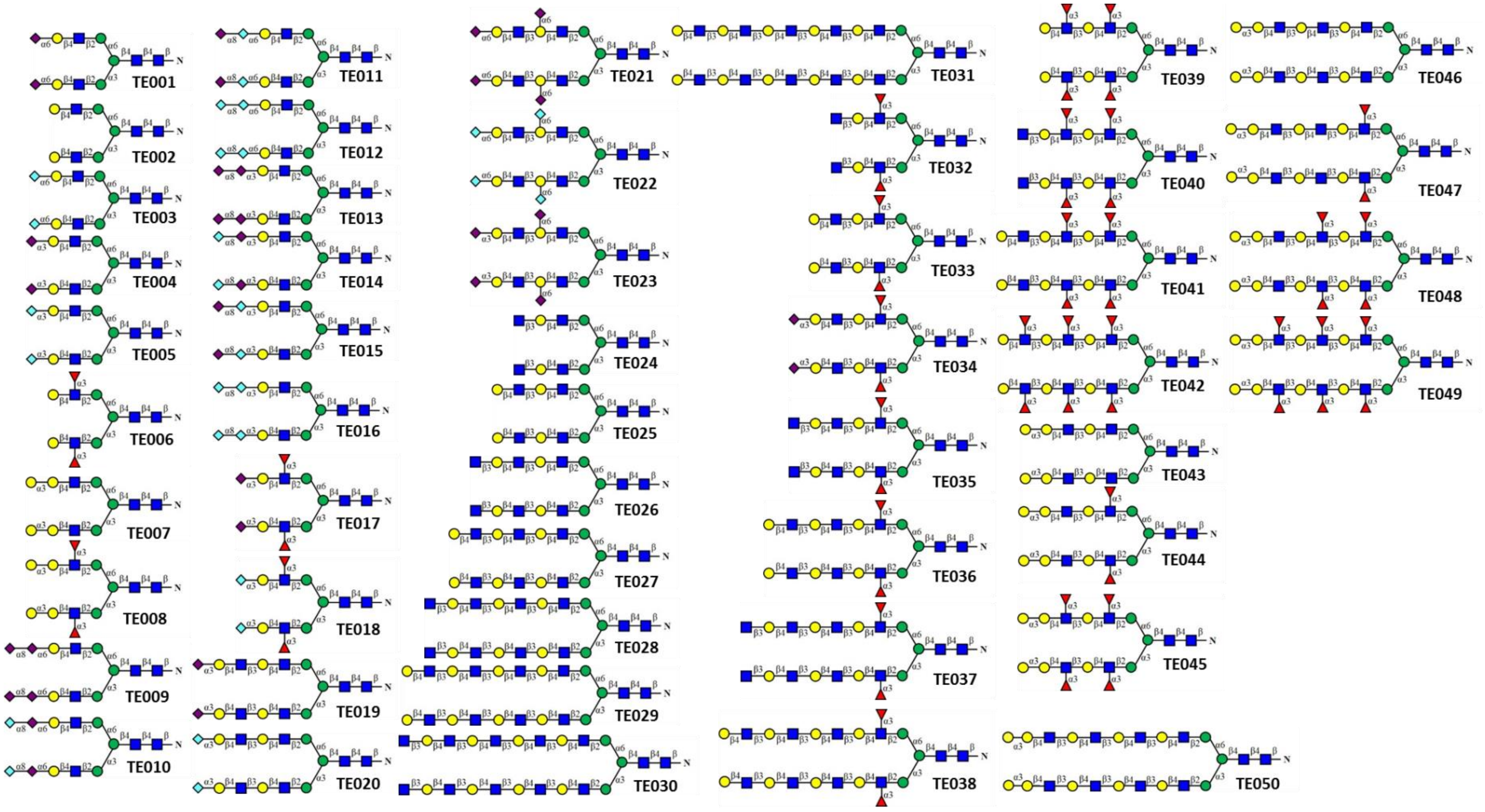
